# A gut-secreted peptide controls arousability through modulation of dopaminergic neurons in the brain

**DOI:** 10.1101/2020.08.31.275552

**Authors:** Iris Titos, Dragana Rogulja

## Abstract

Since sensory information is always present in the environment, animals need to internally regulate their responsiveness to fit the context. During sleep, the threshold for sensory arousal is increased so that only stimuli of sufficient magnitude can cross it. The mechanisms that make arousability flexible are largely mysterious, but they must integrate sensory information with information about physiology. We discovered a gut-to-brain signaling pathway that uses information about ingested nutrients to control arousability from sleep, without affecting sleep duration. Protein ingestion causes endocrine cells in the *Drosophila* gut to increase production of CCHa1, a peptide that decreases sensory responsiveness. CCHa1 is received by a small group of brain dopaminergic neurons whose activity gates behavioral responsiveness to mechanical stimulation. These dopaminergic neurons innervate the mushroom body, a brain structure involved in determining sleep duration. This work describes how the gut tunes arousability according to nutrient availability, allowing deeper sleep when dietary proteins are abundant. It also suggests that behavioral flexibility is increased through independent tuning of sleep depth and duration.

---

Arousability is dynamically regulated, allowing context-appropriate behavioral responsiveness and facilitating deep sleep. Flies share with mammals all the behavioral hallmarks of sleep, including decreased responsiveness to stimuli^1,2^. While work in flies led to identification of neural circuits and conserved genes that regulate sleep duration^3^, much less is known about the regulation of sleep depth^4–8^. Since the restorative function of sleep depends on its quality and not just duration^9^, it is important to identify the molecular and circuit mechanisms that set arousal threshold.

We probed the arousal threshold of sleeping flies with mechanical stimulation. In our setup, loudspeakers convert a tunable electrical signal into 2-second-long vibrations every 30-40 minutes throughout the night (intervals are randomized to minimize habituation). Responsiveness to vibrations is automatically tracked for individual animals (**Extended Data Fig. 1a,b**, and Experimental Procedures). We determined the amplitudes sufficient to wake up either ~20% or ~95% of control flies and used these settings to screen for genetic manipulations that increased or decreased sensory responsiveness. We used elav-Gal4, a driver mainly expressed in neurons^10^, to target ~3,400 genes with RNAi (the screen was unbiased). The screen produced both hyper- and hypo-arousable phenotypes (**Fig. 1a**) (**Extended Data Table 1**). A Gene Ontology (GO) term analysis (see Experimental Procedures) revealed diversity in biological processes, molecular functions and cellular localization for the genes whose mRNA knockdown produced arousability phenotypes (**Extended Data Fig.1c**). We focused on the neuropeptide CCHamide-1 and its receptor CCHa1R, since both have strong phenotypes and belong to the same pathway. CCHamides (CCHa1 and CCHa2) are recently discovered neuropeptides with functions related to feeding and growth regulation (CCHa2), and olfactory responsiveness and circadian behavior (CCHa1)^11–16^. CCHa1R is a G protein-coupled receptor homologous to the mammalian Gastrin-Releasing Peptide Receptor^16^ which regulates itch sensation^17^ and responsiveness to social interactions^18,19^.

**Figure 1.**
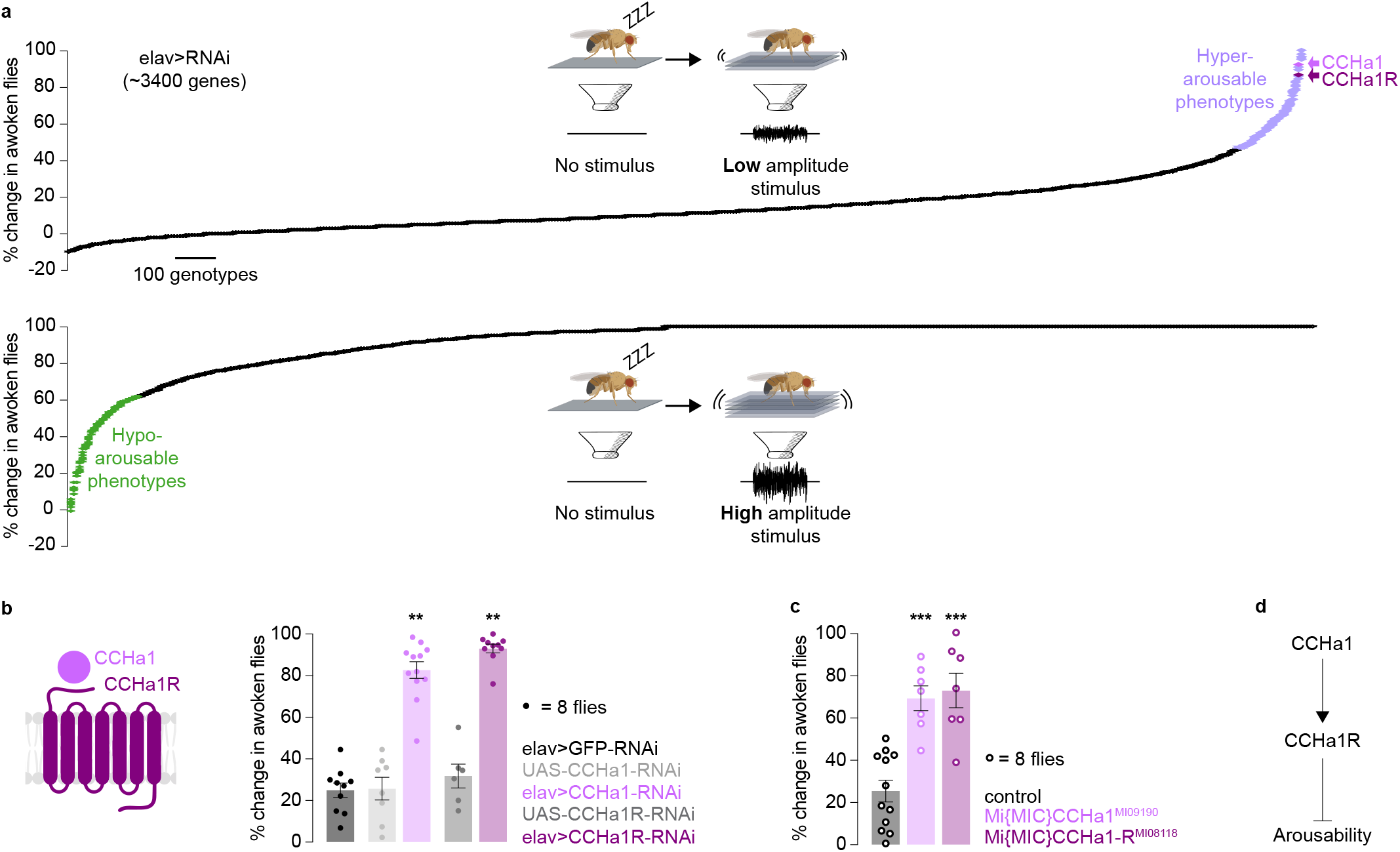
CCHa1 and CCHa1R suppress arousability. **(a)** Results of a screen for genes that regulate arousability. Low amplitude stimulus uncovers hyper-responsive phenotypes, while high amplitude stimulus uncovers hypo-responsive phenotypes (phenotypes exceed two standard deviations from the mean). **(b)** elav-Gal4-driven RNAi against CCHa1 (elav>CCHa1-RNAi) or its receptor (elav>CCHa1R-RNAi) increases arousability of sleeping animals. **(c)** CCHa1 and CCHa1R mutant flies are easier to wake up during sleep than control flies. **(d)** Schematic: CCHa1 and CCHa1R suppress arousability. In all panels, mean and S.E.M. are depicted; sample sizes and statistical analyses are shown in Extended Data Table 2.

RNAi against either CCHa1 or CCHa1R led to hyper-arousability, with ~85% of animals waking up in response to low-amplitude stimulation that woke up only ~20% of the parental controls (**Fig. 1b**, elav>CCHa1-RNAi and elav>CCHa1R-RNAi). We noticed that either the peptide or the receptor depletion caused flies to respond more readily to stimulation even when awake (as reported by locomotor response upon stimulation), without increasing basal locomotion (**Extended Data Fig. 2a**). If undisturbed, CCHa1-R-knockdown animals slept less than the controls and CCHa1 knockdown flies had more fragmented sleep (**Extended Data Fig. 2a**). The specificity of RNAi for CCHa1 and CCHa1R (and not CCHa2 and CCHa2R) was shown with quantitative RT-PCR (**Extended Data Fig. 2b**). Additional RNAi lines (**Extended Data Fig 2c,d**), as well as transgenic flies containing the MiMIC constructs from the Gene Disruption Project (Mi{MIC}CCHa1^MI09190^ and Mi{MIC}CCHa1R^MI08118^ that affect CCHa1 and CCHa1R^20^ respectively) (**Fig. 1c and Extended Data Fig. 2e**), confirmed the role for these genes in arousability. Total sleep was somewhat reduced. A conditional approach allowed depletion of CCHa1 and CCHa1R during adulthood, which was sufficient to cause hyper-arousability (**Extended Data Fig. 2f**). We concluded that CCHa1, likely signaling through CCHa1R, suppresses sensory arousability in adults (**Fig. 1d**).

To find the relevant CCHa1-producing cells, we raised antibodies against CCHa1 (using a previously validated epitope^21^) and expressed CCHa1-RNAi with spatially restricted Gal4s. The antibody signal agreed with previous reports^13,16,21^ (**Fig. 2a**) and with the pattern of CCHa1-promoter-driven GFP (**Extended Data Fig. 3a**). The signal was abolished in elav>CCHa1-RNAi flies (**Fig. 2a**) and in homozygous CCHa1 mutants (**Extended Data Fig. 3a**). From ~100 Gal4 lines, we found one that recapitulated the phenotype obtained with elav-Gal4 (**Fig. 2b**). This line did not noticeably change, or even overlap with, the CCHa1 signal in the nervous system (EECG>CCHa1-RNAi in **Fig. 2a**, EECG>GFP in **Extended Data Fig. 3b**), indicating that the hyper-arousability phenotype seen with CCHa1 depletion may have originated in a non-neuronal tissue. CCHa1 was previously detected in a posterior subpopulation of enteroendocrine cells^21^ a type of secretory cell sparsely distributed throughout the gut^22^ (**Extended Data Fig. 3c**). Enteroendocrine cells and neurons share developmental programs^22^, which could explain why elav-Gal4, the pan-neuronal driver we used for RNAi expression, is also expressed in enteroendocrine cells^23^ (**Extended Data Fig. 3d**). The CCHa1 signal in the gut indeed disappeared when CCHa1-RNAi was driven by elav-Gal4 (**Fig. 2a and Extended Data Fig. 3e)**. The restricted driver that produced the arousal phenotype without altering neuronal CCHa1 levels also depleted CCHa1 from the gut (**Fig. 2a,b and Extended Data Fig. 3e**). We call this driver EECG-Gal4 (<u>E</u>ntero<u>E</u>ndocrine <u>C</u>CHa1-positive cells in the <u>G</u>ut – Gal4). The gut-specific CCHa1 depletion (EECG>CCHa1-RNAi) led to increased arousability both during sleep and wakefulness (**Fig. 2b and Extended Data Fig. 3f**), similarly to what was seen with elav-Gal4 (**Fig. 1b and Extended Data Fig. 2a**). All the phenotypes observed with EECG-Gal4 were reproduced with a conditional version of pros-Gal4^24^, (**Fig. 2a,b and Extended Data Fig. 3b,e and g**) a classic driver for enteroendocrine cells. Depletion of CCHa1 from the gut had no effect on daily sleep amount, or on how much food animals ingested (**Extended Data Fig. 3f, g and h**). As there was no change in sleep duration, we could study specifically how gut-produced CCHa1 regulates arousability (i.e. sleep depth).

**Figure 2.**
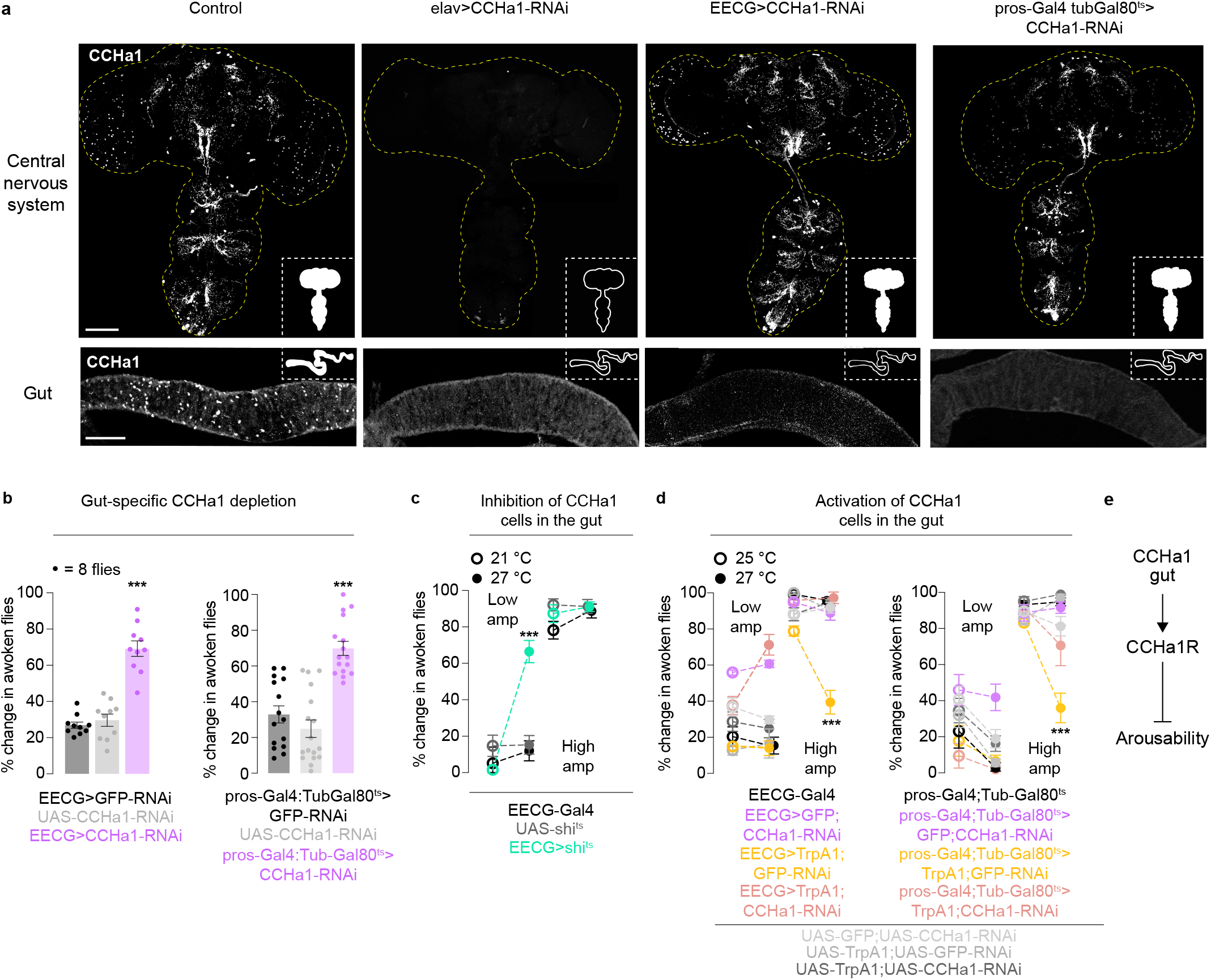
CCHa1 signals from the gut to regulate arousability. **(a)** Antibodies against CCHa1 reveal sparse expression in the nervous system and the posterior midgut. CCHa1 expression is abolished with elav-Gal4-driven CCHa1-RNAi (elav>CCHa1-RNAi). CCHa1-RNAi driven in enteroendocrine cells (EECG>CCHa1-RNAi and pros-Gal4:TubGal80^ts^>CCHa1-RNAi) does not affect expression of CCHa1 in the nervous system, but decreases it in the gut. **(b)** Knockdown of CCHa1 in the gut decreases arousal threshold. **(c)** Inhibiting EECGs using shi^ts^ increases arousability. **(d)** Activating enteroendocrine cells using EECG-Gal4 or pros-Gal4:TubGal80^ts^ to drive TrpA1 decreases arousability in a CCHa1-dependent manner. The temperature for this experiment needs to be at least 25 °C in order to ensure CCHa1-RNAi expression (Gal4 activity is low at 21 °C). **(e)** CCHa1 from the gut suppresses arousability. In all panels, mean and S.E.M. are depicted; sample sizes and statistical analyses are in Extended Data Table 2. Scale bars: 100 μm.

Mammalian enteroendocrine cells are electrically excitable^25^. In *Drosophila*, activation of enteroendocrine cells increases intracellular calcium and promotes vesicle release^26^. We used neuronal activation and silencing tools to ask whether changing the activity of enteroendocrine cells can also impact arousability. Conditional silencing (with a temperature-sensitive allele of a dynamin homolog shibire, shi^ts27^) or conditional activation (with a temperature-sensitive cation channel TrpA1^28^) produced strong and opposite phenotypes (**Fig. 2c,d**). Silencing EECGs phenocopied CCHa1 depletion, with animals reacting more readily to vibrations (**Fig. 2c and Extended Data Fig. 4a)**. When enteroendocrine cells were activated (using EECG-Gal4 or pros-Gal4;Tub-Gal80^ts^) animals were harder to arouse with vibrations (**Fig. 2d**). The phenotype was suppressed if the cells could not produce CCHa1, demonstrating that CCHa1 mediates the effect of enteroendocrine cells on arousability (**Fig. 2d and Extended Data Fig. 4b and 5**). Activation of enteroendocrine cells reduced total sleep in a CCHa1-independent manner (**Extended Data Fig. 4b and 5**) suggesting that non-CCHa1 cells labeled with those Gal4s could have a role in regulating sleep duration. We concluded that enteroendocrine cells use the peptide CCHa1 to control arousability (**Fig. 2e**).

Gut cells are positioned to absorb nutrients and assess food quality, and peptides produced and secreted by the enteroendocrine cells could be used to propagate this information^14,22,29^. Food satiety may increase not only sleep duration^30–32^ but also sleep depth by suppressing arousability. We asked if the activity of enteroendocrine cells, and their production of CCHa1, changes in response to three basic dietary components: sugar, fat and protein. To measure cell activity, we used pros-Gal4 to drive CaLexA^33^ (**Fig. 3a**), a GFP-based reporter previously shown to register calcium changes in the fat body, neurons and enteroendocrine cells^26^. Supplementing standard fly food with sugars (glucose, galactose or fructose) or fat (coconut oil, propionic acid or hexanoic acid) had no effect at the tested concentrations; in contrast, with peptone as an additional protein source, the activity of CCHa1-positive enteroendocrine cells was increased, as were their CCHa1 levels (**Fig. 3b,c and Extended Data Fig. 6a**). The magnitude of the response depended on how long the flies had access to supplemented food, since the increase in CCHa1 levels was stronger after 24 hours than after 6 hours of peptone feeding (**Extended Data Fig. 6b**). Not all cells with high CCHa1 levels showed evidence of activation, and some activated cells did not label for CCHa1 (**Fig. 3b**); the latter observation suggests that other peptides made in this area of the gut could signal protein ingestion and affect behavior.

**Figure 3.**
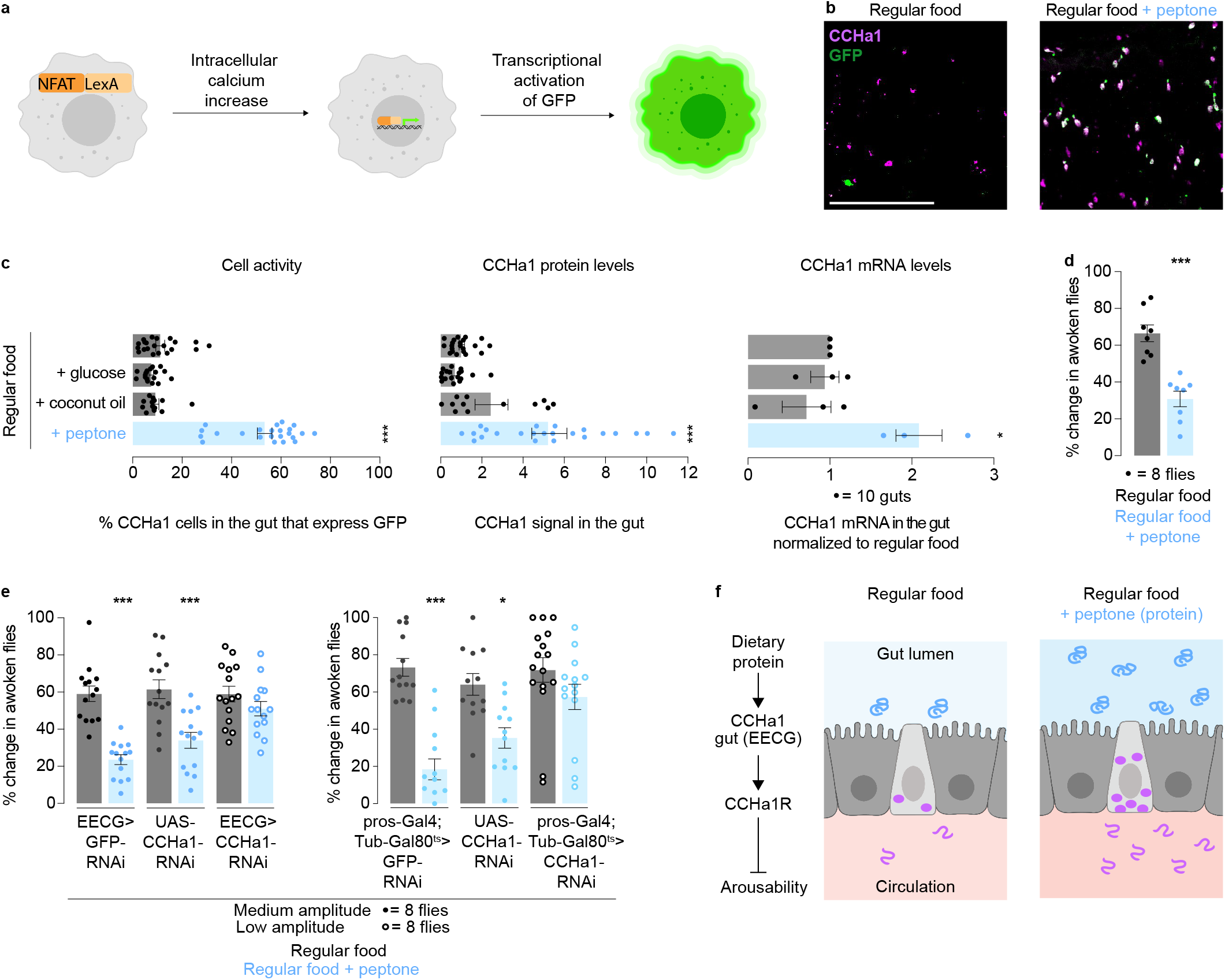
CCHa1 in enteroendocrine cells responds to dietary proteins. **(a)** The CaLexA method detects calcium-dependent cell activation. **(b)** CCHa1 mRNA (measured by qRT-PCR, each dot represents 10 guts) and protein levels, and CaLexA-dependent GFP signal, increase in the posterior midgut when flies are fed protein-enriched food. **(c)** Supplementing regular food with protein (peptone), but not fat (coconut oil) or sugar (glucose), activates enteroendocrine cells and increases CCHa1 protein and mRNA levels. **(d)** Protein-enriched diet decreases arousability. **(e)** Knocking down CCHa1 specifically in the gut with EECG-Gal4 or pros-Gal4:Tub-Gal80^ts^ diminishes the peptone-induced changes in arousability. **(f)** Schematic: protein ingestion activates CCHa1-producing cells and increases the neuropeptide levels to suppress arousability. In all panels, mean and S.E.M. are depicted; sample sizes and statistical analyses are in Extended Data Table 2. Scale bars: 100 μm.

Peptones are made by partial hydrolysis of dietary proteins and contain polypeptides and amino acids. To investigate if specific peptone components activate CCHa1-producing enteroendocrine cells, we supplemented regular food with different combinations of amino acids, matching their respective concentrations in peptone. Adding single essential vs non-essential or biochemically-related amino acids to the food did not recapitulate the CCHa1 increase observed with peptone (**Extended Data Fig. 6c**). However, the mix of all amino acids (which matched the total amino acid concentration without changing their individual representation with respect to peptone) strongly induced CCHa1 production (**Extended Data Fig. 6c**). The most likely explanation is that EECGs do not receive signals from specific amino acids but monitor whether food is generally protein-rich (**Extended Data Fig. 6d**), and adjust arousability accordingly. Peptone supplementation indeed suppressed sensory arousal; only half as many wild-type flies fed peptone-supplemented food woke up in response to vibrations compared to flies fed regular food (**Fig. 3d**). Peptone feeding had a modest effect on responsiveness during wakefulness, basal locomotion, sleep duration and feeding (**Extended Data Fig. 6e and f**). To determine whether CCHa1 mediates the effect of peptone on arousal, we repeated the feeding experiments in animals lacking CCHa1 in the gut. Because removal of CCHa1 by itself causes hyper-arousability, lower amplitude stimulation was used in animals lacking the peptide (EECG>CCHa1-RNAi and pros-Gal4:TubGal80^ts^>CCHa1-RNAi) than in the parental controls. This way, baseline arousability (on regular food) could be matched across genotypes (**Fig. 3e and Extended Data Fig. 6g**). CCHa1 knockdown weakened the effect of peptone on arousal (**Fig. 3e and Extended Data Fig. 6g**), though incomplete phenotype suppression reinforces the idea that dietary proteins activate multiple signaling pathways. Together, our results show that ingested proteins activate CCHa1-producing cells in the gut to suppress arousability (**Fig. 3f**).

Peptides produced by enteroendocrine cells can either signal through synapses or be secreted into circulation^22^. CCHa1-producing cells reside in a poorly innervated region^22^, suggesting that CCHa1 works as a hormone. We looked for CCHa1-receiving cells by knocking down the receptor with different Gal4 drivers and testing arousability. Flies were not hyper-arousable when CCHa1R was knocked down in cholinergic (Cha-Gal4), octopaminergic (Tdc2-Gal4), glutamatergic (vGlut-Gal4), or circadian (Tim-Gal4) neurons (**Extended Data Fig. 7a**). Though one dopaminergic driver (TH-Gal4) produced no phenotype, another dopaminergic driver with a broader expression pattern (Ddc-Gal4) produced a strong phenotype (**Fig. 4a**). TH-Gal4 covers the majority of dopamine neurons but spares the PAM cluster, labeled by Ddc-Gal4^34^. Flies were indeed hyper-arousable when CCHa1R-RNAi was driven in the ~87 neurons^34^ that comprise the PAM dopaminergic cluster in each brain hemisphere (PAM-Gal4) (**Fig. 4a**). Arousability during wakefulness was weakly increased, while basal locomotion and sleep duration were unaffected by these manipulations (**Extended Data Fig. 7b**). Two different Gal4s expressed under the control of CCHa1R regulatory elements label the PAM dopaminergic cluster (**Extended Data Fig. 7c**), agreeing with the notion that the receptor is expressed there.

**Figure 4.**
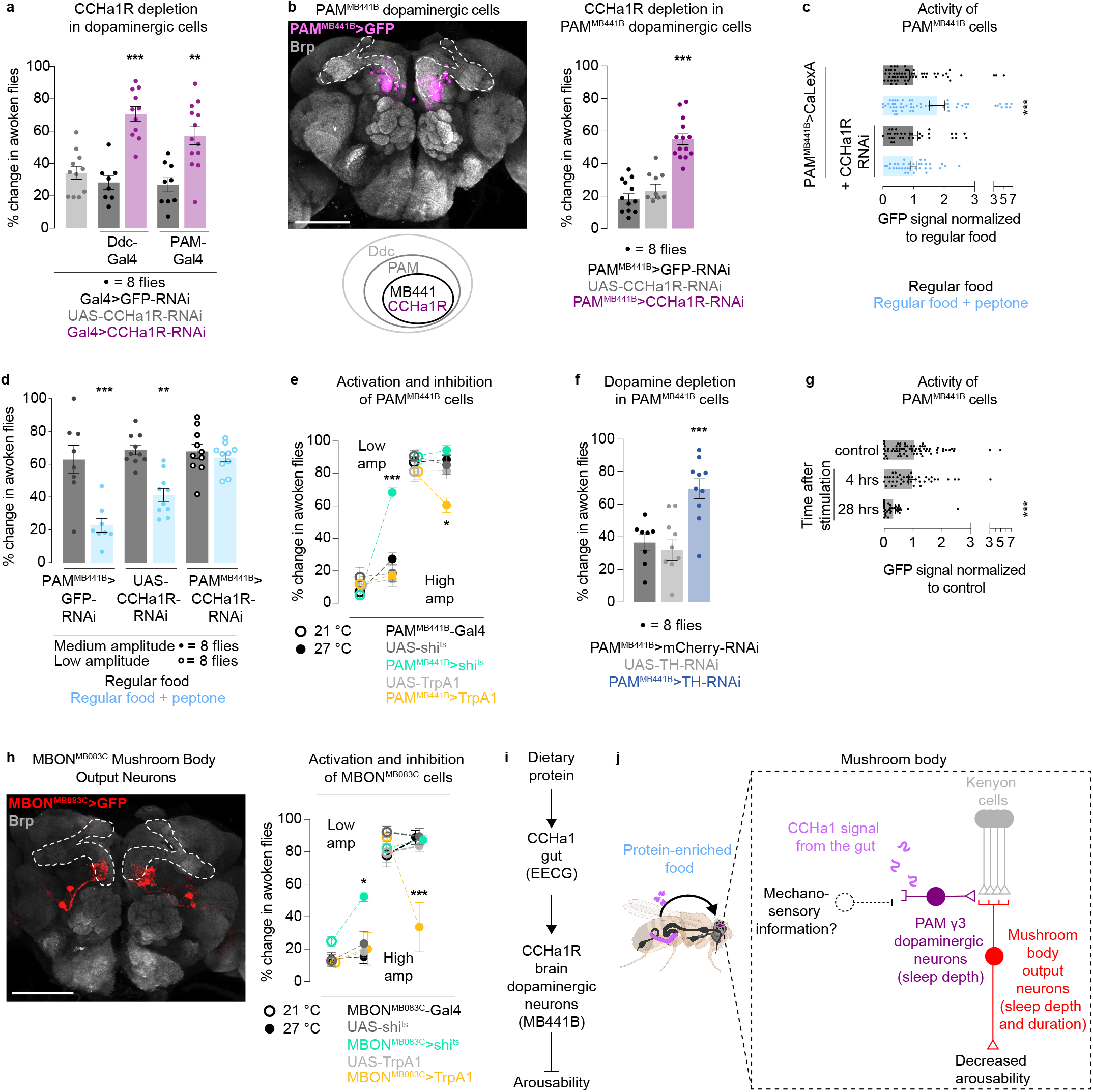
CCHa1 regulates arousability through dopaminergic neurons in the brain. **(a)** Knockdown of the CCHa1 receptor using dopaminergic drivers increases arousability from sleep. **(b)** PAM^MB441B^-Gal4 expression in the brain visualized with a membrane-targeted GFP. Presynaptic protein Brp marks the neuropil. Dashed white line marks the mushroom bodies. Schematic of the expression pattern of the Gal4s used in this figure. Knocking down CCHa1 receptor in PAM^MB441B^ neurons increases arousability from sleep. **(c)** Protein-enriched diet activates PAM^MB441B^ neurons, as visualized with a calcium-dependent GFP reporter CaLexA, in a CCHa1R-dependent manner. **(d)** Knocking down CCHa1R in PAM^MB441B^ neurons diminishes the peptone-induced changes in arousability. **(e)** Inhibition (with shi^ts^) and activation (with TrpA1) of PAM^MB441B^ neurons increases and decreases arousability, respectively. **(f)** Knocking down tyrosine hydroxylase (TH) in PAM^MB441B^ neurons increases arousability from sleep. **(g)** Mechanical vibrations reduce the activity of PAM^MB441B^ neurons, as visualized with a calcium-dependent GFP reporter CaLexA 4 and 28 hours after exposure to mechanosensory stimuli. **(h)** Mushroom body output neurons from the γ3 compartment, MBON^MB083C^. Dashed white line indicates mushroom bodies. Inhibition of MBON^MB083C^ with shi^ts^ increase arousability in response to vibrations, while activation with TrpA1 suppresses arousability. **(i)** CCHa1 levels in the gut increase in response to dietary protein and target a sparse group of dopaminergic neurons in the brain to decrease arousability. **(j)** A model for adjustment of sensory gating by gut-brain interactions. In all panels, mean and S.E.M. are depicted; sample sizes and statistical analyses are shown in Extended Data Table 2. Scale bars: 100 μm.

Thanks to a large-scale mapping and tool-generating effort^35^, access to subpopulations of PAM neurons is possible, which allowed us to further narrow down the CCHa1 target cells. Subsets of PAM neurons have been implicated in regulating sleep duration and food foraging^36–38^. We found that depleting CCHa1R from a small PAM subpopulation labeled by PAM^MB441B^-Gal4^35^ caused flies to wake up more easily (**Fig. 4b**). PAM^MB441B^-Gal4 labels ~8-10 neurons per hemisphere and has no expression in the gut (**Fig. 4b and Extended Data Fig. 7d**). Recently, those neurons were found to express CCHa1R^39^. Depletion of CCHa1R from PAM^MB441B^ neurons did not change sleep duration or feeding, but it decreased basal locomotion (**Extended Data Fig. 7e,f**). In contrast to all other manipulations described so far, depleting CCHa1R from this narrow dopaminergic cell population did not affect responsiveness of awake flies (**Extended Data Fig. 7e**). This suggests that CCHa1 is received by a separate population of CCHa1R-expressing dopaminergic neurons in the PAM cluster to regulate arousability during wakefulness, meaning that sensory gating during sleep and wakefulness can be uncoupled.

If brain dopaminergic neurons receive CCHa1 from the gut, their activity should change in response to inputs that activate the CCHa1-producing enteroendocrine cells, in a CCHa1R-dependent manner; further, behavioral effects stemming from increased CCHa1 production should be abolished by PAM^MB441B^-specific CCHa1R depletion. In agreement, peptone supplementation nearly doubled the calcium-activated GFP signal in PAM^MB441B^ neurons (**Fig. 4c**), and this was dependent on CCHa1R: PAM^MB441B^-specific CCHa1R depletion prevented the neurons from being activated by dietary peptone (**Fig. 4c**). This manipulation also abolished the effect of peptone feeding on arousability (which was specific for sleep state) (**Fig. 4d and Extended Data Fig. 7g**). Next, we investigated how the activity of PAM^MB441B^ neurons affects sensory responsiveness. Conditional silencing of these neurons caused hyper-arousability, while their conditional activation caused hypo-arousability, specifically during sleep (**Fig. 4e** and **Extended Data Fig. 8a**). Sleep duration and locomotion were unaffected (**Extended Data Fig. 8a**). Knocking down Tyrosine Hydroxylase (TH, the rate-limiting enzyme in dopamine synthesis) in PAM^MB441B^ neurons caused hyper-arousability during sleep (**Fig. 4f**), arguing that dopamine mediates the effect of PAM^MB441B^ neurons on arousal. Again, sleep duration and responsiveness during wakefulness were unchanged (**Extended Data Fig. 8b**). Dopamine, perhaps the best known arousal-promoting molecule^40–43^ can therefore suppress arousal by acting on specific synapses.

The PAM dopaminergic neurons, including the MB441B population, project to the mushroom body, a brain structure that receives sensory inputs from different modalities^44–49^ and regulates olfaction, memory and sleep duration^38,50–55^. PAM neurons modulate synapses between Kenyon cells (mushroom body intrinsic neurons) and the mushroom body output neurons (MBONs)^56^ to contextualize and drive appropriate behavioral responses^38,56^. So far, we talked about a physiological signal (CCHa1) acting on modulatory neurons (dopaminergic PAMs). But where does the sensory information itself fit in, where is it received in the system? Kenyon cells carry olfactory and visual information^49,57^ but there is no evidence yet that they receive mechanical inputs. The larval^58^ and adult^59^ PAM neurons on the other hand are known to receive such information. We targeted CaLexA to PAM^MB441B^ neurons and exposed flies to mechanical stimuli used to test arousal threshold (**Fig. 4g**). The GFP signal in the PAM^MB441B^ cell bodies was quantified 4 hours after stimulation (which should be enough time for detecting an increase in GFP signal), in case neurons are activated, as well as 28 hours after stimulation (which should be enough time for GFP degradation and signal decrease, in case neuronal activity is reduced^60^). CaLexA reported that PAM^MB441B^ neurons are inhibited in response to vibrations (**Fig. 4g**). These results suggest that PAM^MB441B^ neurons could regulate arousal by integrating mechanosensory information with a physiological signal from the gut. Arousability is attenuated across sensory modalities during sleep^33^; an outstanding question is whether different senses can be tuned independently. To address this, we asked whether the CCHa1 pathway regulates arousability in response to other sensory stimuli besides mechanical vibrations. We built a setup for rapid temperature increase that was effective at waking flies up with heat (35-40 °C) (**Extended Data Fig. 9a**). Unlike for vibrations, the CCHa1 pathway had no bearing on the ability of thermal stimuli to elicit arousal (**Extended Data Fig. 9b**). In agreement with behavioral data, there was no change in the activity of PAM^MB441B^ neurons when flies were exposed to thermal stimuli (**Extended Data Fig. 9c**). Gating of different sensory modalities is therefore modular, perhaps allowing greater flexibility in adjusting behaviors to external and internal circumstances.

PAM^MB441B^ neurons project to the γ3 compartment of the mushroom body^35^. In each brain hemisphere, two MBONs (labeled by MBON^MB083C^-Gal4 which shows no expression in the gut, (**Fig. 4h and Extended Data Fig. 9d**)) receive information from this compartment and transmit it to other brain centers^35,38^. Conditional silencing of the MBON^MB083C^ caused flies to be easily awoken with vibrations, while conditional activation had the opposite effect (**Fig. 4h**). Responsiveness was not changed in awake animals (**Extended Data Fig. 9e**). In contrast to the upstream CCHa1 pathway components we described, MBON^MB083C^ also have a role in determining sleep duration: changing their activity can bidirectionally modulate the amount of sleep (**Extended Data Fig. 9e**). Other studies have reported that altering the activity of Kenyon cells or MBON can impact sleep duration^3^. Our results point to convergence of circuits that control sleep amount and sleep depth. They also specifically demonstrate that dietary proteins boost production of CCHa1 in the gut, and that CCHa1 targets dopaminergic neurons in the brain to suppress arousability (**Fig. 4i**).

Animals need to decide which of the many environmental cues are worth responding to. Elevated arousal threshold is a core feature of sleep^61^ and the mechanisms that allow flexible arousability are likely conserved. In mammals, the flow of sensory information is thought to be gated by the thalamus^57,62^ but olfactory information can be modulated despite not passing through this brain region^63^. This suggests that different sensory modalities can be regulated independently, a view supported by our results. Multiple sensory streams are received by the mushroom body^35,49^. This brain structure is organized in a modular way^35^, which might be particularly suited for modular processing of sensory cues. Dopamine itself is a key regulator of arousal across species^64^ but there is little understanding of how the dopaminergic system itself is regulated. We describe a pathway that informs specific dopaminergic neurons, which seem to also receive sensory information, about ingested nutrients, thereby tuning them to physiology (**Fig. 4j**). The logic of the system has similarities with the dopaminergic tuning of sexual arousal by signals from the reproductive periphery^65,66^. Abundance of molecules from food could signal an opportunity to sleep more deeply since there is no need to forage; peptides like CCHa1 generally have longer-lasting effects than classical neurotransmitters^67^ and are ideal for regulating states like sleep that can last for hours^3^. Nutritional status is a critical variable for many behavioral outcomes so there are likely to be different signaling axes like the one we describe, affecting multiple behaviors. The molecular and functional diversity of gut endocrine cells is increasingly appreciated as fundamental for regulating processes other than digestion^22^. The gut influences behavior more profoundly than previously appreciated through peptide signals that can diffuse to the brain or through direct connections with the nervous system (notably via the vagus nerve^68^). Our study adds to the emerging knowledge on the basic mechanisms of gut-brain interactions^22^. Recent evidence links gut dysfunction with neurological diseases like the autism spectrum disorders, Parkinson’s and Alzheimer’s disease^69^. Together, these studies support the centuries-old idea that digestive and nervous systems are critically linked in normal physiology and disease^70^.

## ACKNOWLEDGEMENTS

We thank our lab and Michael Crickmore’s lab for advice and comments on the manuscript. Ofer Mazor and Pavel Gorelik assisted in building the arousal threshold setup. Benjamin Sanchez developed the software used to control mechanical stimuli and analyze the behavioral responsiveness. Keishi Nambara and Alexa Soares assisted with the screen. For fly stocks, we thank Michael Crickmore, Nicholas Stavropoulos, Kyunghee Koh and Norbert Perrimon. We thank Pedro Saavedra for assistance with the quantitative RT-PCR. I.T. was supported by the Mahoney and Brooks postdoctoral fellowships. D.R. is a New York Stem Cell – Robertson investigator. This work was supported by the New York Stem Cell Foundation, the NIH (DP2 OD022385) and the Pew Scholars Program in the Biomedical Sciences.

## AUTHOR CONTRIBUTIONS

I.T. and D.R. designed the study. I.T. performed the experiments. I.T. and D.R. analyzed the data and wrote the paper.

## EXPERIMENTAL PROCEDURES

### Fly stocks

Flies were grown in 12 hour light: 12 hour dark cycles at 25 °C unless indicated otherwise. Experiments were performed in Percival Scientific Inc. (DR36VL model) incubators. Stocks used include: wild type iso31, elav-Gal4, UAS-TrpA1 and UAS-Shi^ts^ (gift from Kyunghee Koh), elav-Gal4;UAS-Dcr2 (gift from Nicholas Stavropoulos), PAM-Gal4 (BDSC #41347), UAS-CCHa1-RNAi (BDSC #57562), UAS-CCHa1-RNAi #2 (VDRC #104074), UAS-CCHa1R-RNAi (BDSC #51168), UAS-CCHa1R-RNAi #2 (VDRC #103055), UAS-GFP-RNAi (BDSC #41552), EECG-Gal4 (BDSC #49037), Tdc2-Gal4 (BDSC #9313), vGlut-Gal4 (BDSC #26160), MB441B-split-Gal4 (BDSC #68251), MB083-split-Gal4 (BDSC #68287/68262), CaLexA (BDSC #66542), UAS-nlsGFP (BDSC #4776), UAS-TH-RNAi (BDSC #25796), UAS-Dcr2 (BDSC #24650), Mi{MIC}CCHa1^MI09190^ (BDSC #51261), Mi{MIC}CCHa1R^MI08118^ (BDSC #44750), TI{2A-GAL4}CCHa1-R[2A-GAL4] (BDSC #84601), P{GMR23H07-GAL4}attP2 (BDSC #47498), and UAS-CD8GFP, UAS-nlsLacZ, TH-Gal4, ddc-Gal4 and Cha-Gal4 (gift from Michael Crickmore), Tim-Gal4 (gift from Michael Young) and pros-Gal4 (gift from Norbert Perrimon). All the stocks used for behavioral experiments were outcrossed at least 5 times to an isogenised (iso31) genetic background. In the case of UAS-CCHa1-RNAi and UAS-CCHa1R-RNAi the UAS-GFP-RNAi from the same library was used as a control.

The RNAis against CCHa1R target non-overlapping sequences: VDRC #103055 targets CCHa1R coding sequence nucleotides 16 to 333, and BDSC #51168 targets CCHa1R coding sequence nucleotides 375 to 397. The RNAis used for CCHa1 target sequences that partially overlap: VDRC #104974KK targets CCHa1 coding sequence nucleotides 241 to 539 and BDSC #57562 targets CCHa1 coding sequence nucleotides 525 to 547.

For all the Gal4/UAS experiments, heterozygous controls were used: (UAS crossed to the Gal4 genetic background, and Gal4 crossed to the UAS genetic background).

### Fly media

Flies were raised on cornmeal-agar medium^71^. Briefly, our regular food contained 7.3% agar (Spectrum), 12.7% yeast (MP Biomedicals), 56% sugar (Domino) and 24% cornmeal (Bunge). These ingredients were heated until the temperature reached 97 °C while stirring. After the temperature of the solution decreased to 75 °C, 2.5% Tegosept, 23.8 g/L Ethanol (Koptic #V1001) and water to compensate for evaporation were added.

### Arousal threshold screen

UAS-RNAi lines were from Bloomington *Drosophila* Stock Center, Kyoto Stock Center of *Drosophila* Genomics and Genetic Resources, Vienna *Drosophila* Resource Center and TRiP library generously shared by Norbert Perrimon’s laboratory. These lines were selected randomly (the screen was unbiased) and crossed to flies expressing elav-Gal4. 8 males, 3-5 days old, were tested in the system described below. Each dot represents the behavior of 8 flies. In our experiments, the flies were exposed to low amplitude stimulation (0.2V) on the first night, and high amplitude stimulation (2V) the second night.

### Analysis of screen results

For the genes whose disruption in flies caused hypo- or hyper-arousability phenotypes gene ontology analysis was performed. We used the GO Term Mapper (http://go.princeton.edu/cgi-bin/GOTermMapper) to bin individual GO categories into more general terms, to facilitate the analysis. In order to normalize the different GO terms taking into account their representation in the genome, for every GO term the % of hits containing that term in our screen was normalized to the % of genes in the whole genome that contain that term. The data are represented on a log10 scale.

### Locomotor activity and sleep monitoring

Locomotor activity was recorded using the *Drosophila* Activity Monitoring system (DAM3, Trikinetics, Waltham, MA). Flies were individually housed in 65 mm-long glass tubes (TriKinetics) filled with ~40 mm of cornmeal-agar food, unless otherwise indicated. This left ~25 mm of space for flies to walk back and forth in. An infrared light beam bisects this space and gets interrupted when flies walk through; this is automatically scored as movement. Sleep was defined as five minutes of inactivity^1,2^. Sleep quantification was performed with a custom MATLAB software (available on github at https://github.com/CrickmoreRoguljaLabs/SleepAnalysis).

### Arousal threshold mechanical setup

DAMs were mounted on custom platforms positioned on top of large bass loudspeakers (381 mm, Delta 15 LFA, Eminence, Eminence, KY). Each speaker/platform holds up to 8 DAMs. To stimulate flies, a custom program (Good Morning, available on github at https://github.com/IrisTitos/ATanalysis/blob/master/Goodmorning.md) written in MATLAB (The MathWorks, Natick, MA) generates a multisine signal (40 Hz - 200 Hz) using a multifunction device (USB-6211, National Instruments, Austin, TX). The signal is fed into an amplifier (SLA-1, Applied Research and Technology, Rochester, NY) connected to the speaker. Using a range of frequencies should avoid frequency-specificity of the response. With this setup, the mechanical vibrations of the loudspeaker translate into up and down movements of the platform on which DAMs are mounted. Low power accelerometers (ADXL335, Analog Devices, Norwood, MA) measure movements of individual DAMs, at different positions on the platform, to ensure that vibrations are uniform across all monitors. In our experiments, 2 second vibrations were delivered at random intervals every 30-40 minutes, starting 1.5 hrs after the lights turn off (for 10 hours during the night (dark phase)).

These data are processed in MATLAB together with locomotor data from DAMs (available on github at https://github.com/IrisTitos/ATanalysis/blob/master/ReadDamsPullDown). Only flies that were asleep (more than 5 minutes of locomotor inactivity) at the moment that the stimulus was applied were considered when arousability from sleep was calculated. If locomotor activity was triggered within the 3 minutes following stimulus application, the fly was considered awoken by vibrations. To control for the increased chance of spontaneous awakenings in genetic manipulations that cause a decrease in sleep duration or increase in sleep fragmentation, we normalized for spontaneous awakenings: % change in awoken flies = 100*(% flies that wake up - % spontaneous awakenings) / (100 - % spontaneous awakenings). To calculate spontaneous awakenings, the activity data from the same night and the same flies used to calculate the “% flies that wake up” was analyzed with a modified timestamp file in which the stimulation timings are shifted by 10 minutes preceding the stimulation. Next, we performed the same analysis previously done with the original timestamp; only flies that were asleep at the modified timestamp were considered for analysis, and flies that produced locomotor activity within the 3 minutes following the modified timestamp were considered to wake up.

To quantify the percentage of flies that react to stimuli during wakefulness we used the same flies and timestamps used to calculate the “% change in awoken flies” since our analysis was post-hoc. In this case, only flies that were awake immediately before the stimulation were taken into account. For those flies we calculated the average activity during the 3 minutes prior to stimulation, and the average activity during the 3 minutes after the stimulation. Flies were considered reacting to stimuli if the average activity after the stimulation was higher than prior to it.

### The setup for probing arousal threshold with temperature

To probe arousability with temperature, we used a flat multi-beam activity monitor (DAM5M Trikinetics) in which the fly-containing tubes lay horizontally. A heating pad (McMaster-Carr 35765K469, Elmhurst, IL) controlled by a 1/16 DIN Ramp/Soak Controller (Omega CN 7800, Norwalk, CT) covers the surface of the monitor, Custom-made software (MATLAB) turns on the heating pad, increasing the temperature to 40 °C (unless otherwise indicated) for 2 minutes, at random intervals (30-40 minutes) during the night. The same protocol as described for the mechanical setup was used to calculate the % of flies awoken by heat.

### Feeding experiments

Male flies, 3 to 5 days old, were transferred from regular food to different food conditions approximately 10 hrs after light turning on (i.e. CT10 in 12:12 light-dark cycles), unless otherwise indicated. After 24 hours, they were dissected or examined for behavior. The cornmeal-agar food was supplemented with 10% coconut oil (Spectrum essentials #L215562P-006), 1% propionic acid (Sigma-Aldrich #P1386), 0.5% hexanoic acid (Sigma-Aldrich #P21530), 200 mM glucose (Sigma-Aldrich #158968-1KG), 200 mM Fructose (Sigma-Aldrich #F0127), 200 mM Galactose (Sigma-Aldrich #G0750), 10% peptone (Apex #20-260), and the single amino acids (Sigma-Aldrich LAA21-1KT) or water as a control.

### Food intake measurement

3-5 day old male flies were transferred from regular food to regular food, or to regular food supplemented with peptone containing 2% (wt/vol) FD&C Blue #1 (Spectrum #FD110). After 24 hours, flies were frozen at −20 °C. Frozen flies were decapitated (to avoid eye the pigment interfering with the measurement), aliquoted in groups of 5 into Eppendorf tubes containing 50 μL of 1% Triton X-100 1X-PBS and homogenized with a motorized pestle (Argos #9950-901). Samples were centrifuged and supernatant was measured at 660 nm (A660) in a NanoDrop 2000 Spectrophotometer (Thermo Fisher Scientific). A standard curve was generated using serial dilutions of the blue dye and the sample dye concentration was extrapolated from that curve.

### Immunohistochemistry

Adult males were anesthetized with carbon dioxide, washed briefly with 70% ethanol (Koptic #V1001) and dissected in 1X-PBS (Phosphate Buffered Saline 10X SeraCare #5460-0030). Guts were fixed overnight at 4 °C in PBS with 4% paraformaldehyde (PFA, Electron Microscopy Sciences #15710). Dissected brains and ventral nervous systems were fixed for 30 minutes in 4% PFA at room temperature. Tissues were washed 3 times for 20 minutes in 1X-PBS containing 0.2% Triton X-100 (Amresco #M143), and then blocked for 2 hours in 1X PBS, 0.2%Triton X-100, 2% bovine serum albumin (BSA Gemini #700-101P) at room temperature. Samples were incubated with primary antibodies overnight (except for Bruchpilot (Brp) stainings, which were incubated for ~48 hours at 4 °C in 1X PBS, 0.2% Triton X-100, 2% BSA at the indicated dilutions). Tissues were washed 3 times for 20 minutes in 1X-PBS, 0.2% Triton X-100 and then incubated with secondary antibodies for 2 hours at room temperature at the indicated dilutions (except for Bruchpilot stainings that were incubated for ~48 hours at 4 °C). Samples were then washed 3 times for 20 minutes in 1X-PBS, 0.2% Triton X-100, and mounted in-between glass slides and coverslips (Electron Microscopy Sciences #72230-01 and 64321-10) in Prolong Gold Antifade medium (Invitrogen #1942345).

Rabbit anti-CCHa1 antibodies were raised against the peptide QIDADNENYSGYELT^21^ by Genscript and affinity purified by the company.

Primary antibodies used: mouse anti-Bruchpilot (1:7, DSHB nc82), rabbit anti-CCHa1 (1:50), chicken anti-GFP (1:1000, Aves #GFP-1020), mouse anti-prospero (1:50, DSHB pros) and mouse anti-tyrosine hydroxylase (TH) (1:1000 ImmunoStar #22941).

Secondary antibodies used: Alexa Fluor 488 goat anti-chicken (1:1000 Life technologies #A11039), Alexa Fluor 488 goat anti-rabbit (1:1000 Life technologies #A11034), Alexa Fluor 647 goat anti-rabbit (1:1000 Life technologies #A21244) Alexa Fluor 647 donkey anti-mouse (1:1000 Life technologies #A31571).

### Fluorescence microscopy

Fluorescent images were acquired with a Leica SP8 confocal microscope. All the imaging conditions remained constant within each experiment, only the Z start and end settings were adjusted individually for each sample. The average distance between Z-stacks was 100 microns.

### Image quantification

Quantification of CCHa1 and GFP signals was performed with Fiji. For every image, a Z-stack summation projection of the whole gut was obtained. To quantify CCHa1 antibody staining in the gut, CCHa1 signal was used to define the region of interest (ROI). The average pixel intensity and area of that ROI was measured. A region with similar area adjacent to the ROI was selected to measure the average background pixel intensity. The amount of CCHa1 was quantified by subtracting the background from the CCHa1 signal and multiplying by the CCHa1 ROI area.

For Fig. 3c and Extended Data Fig. 6a, the percentage of activated CCHa1 cells in the gut was calculated based on CaLexA-dependent GFP transcription. First, we quantified how many CCHa1-expressing cells were present in the posterior midgut using the “counter” function of Fiji. Then, we counted how many of those cells also have GFP signal. To determine the % of CCHa1 cells that were activated, we used the formula: (CCHa1 and GFP-positive cells/all CCHa1-positive cells)*100.

To quantify GFP intensity in Fig. 4c,g and Extended Data Fig. 9c, we projected the summation of the Z-stacks containing the cells of interest. Each cell was manually selected and the average pixel intensity of the GFP signal was quantified. A region next to the cells of interest but containing no GFP positive cells was used to measure the background signal. To calculate the GFP signal we used this formula: (GFP average pixel intensity from the cell – GFP background average pixel intensity)

### Quantitative RT-PCR

For each condition, 2 whole flies, 5 brains or 10 guts of 3-5 days old males were dissected and immediately transferred into 100 μl of ice-cold TRIzol (Thermo fisher #15596026) one by one, flash-frozen on dry ice and stored at −80 °C. To extract RNA, samples were homogenized with a pestle (Argos #9950-901) for 20 seconds on ice before adding 200 μl of TRIzol and 60 μl of chloroform. Samples were then centrifuged for 15 minutes at 15000 rpm 4 °C. The supernatant was collected and diluted in an equal amount of ethanol 99% (Koptec #V1001), mixed and transferred to a purification column (Zymo kit Direct-zol R2061). The manufacturer’s instructions for RNA purification and subsequent DNAse treatment were followed (DNAse Turbo DNA-free kit AM1907). RNA was quantified using NanoDrop. 500 ng of RNA were used for the retrotranscriptase reaction (Biorad iScript cDNA Synthesis #1708890). DNA product was diluted 1:10 and used to set up quantitative PCR reactions using Sybr Green (Biorad #170-8880). Primers were obtained from ID Technologies:

CCHa1 (F-ACTGACGTCGGACAATTTGC and R-ACACGAATGTCCGTATTCCA)^13^
CCHa1R (F-GTTCCAAACACCTACATTTTATCAC and R-CGGATAATGCAGTCAGCGTA)^13^
CCHa2 (F-AAACAGCAACAGCAGCAAAC and R-AGGACCACGGTGCAGATAAC)^13^
CCHa2R (F-CATACCCAACACATACATTCTTTC and R-GAAAGGGCGGTCAGTGTAAA)^13^
rp49 (F-ATCGGTTACGGATCGAACAA and R-GACAATCTCCTTGCGCTTCT)

### Conditional experiments

Adult-specific knockdown of CCHa1 or its receptor was achieved using a temperature-sensitive allele of Gal80, a Gal4 inhibitor (Gal80^ts^). Flies were raised at 21 °C, temperature at which Gal80^ts^ is stable and interferes with Gal4 function. Two days after eclosion, temperature was raised to 29 °C, temperature at which Gal80^ts^ is degraded and the Gal4 driver becomes functional.

For TrpA1 and shi^ts^ experiments, flies were raised at 21 °C, temperature at which TrpA1 and shi^ts^ are not active. Flies were transferred to 27 °C to analyze behavior. In Figures 2d, Extended Data Figure 4b, and Extended Data Figure 5, the experiment needs to be done at least at 25°C to ensure CCHa1-RNAi expression, since the Gal4-UAS system depends on temperature (Gal4 expression is higher at higher temperatures). TrpA1 has been reported to be active at temperatures above 25 °C, as low as 26 °C^12^. We cannot completely discard that some mild TrpA1 activation is occurring at 25 °C. However, it does not seem to be enough to cause behavioral phenotype since we do not observe any arousal threshold phenotypes at 25 °C when EECG-Gal4 drives TrpA1, while arousal is strongly decreased at 27 °C.

### Statistical analysis

Statistical analyses were performed using GraphPad Prism software 7 (GraphPad Software Inc., San Diego, CA). Except for the screen data, all experiments were done 3 times independently. Data was tested for normality and all the statistical tests were two-sided All data are presented as mean ± S.E.M. Please see Table S1 for sample sizes, tests and P values.

## DATA AND SOFTWARE AVAILABILITY

All data and materials are available upon request.

## EXTENDED DATA

**Extended Data Figure 1. Experimental setup and analysis of the genes that regulate arousability, uncovered in our screen. (a)** Picture of the setup used for mechanical stimulation. Custom software generates 2-second-long multi-sine (40-200 Hz) signals that are converted into mechanical stimuli (vibrations). Vibrations are given at random intervals centered on 30-40 minutes throughout the night. DAM monitors are mounted on custom-built platforms on top of the loudspeakers. Vibrations produced by the loudspeaker are translated into vibrations of the DAMs. Accelerometers placed on top of the DAMs measure the stimulus intensities, to ensure that all flies are stimulated equally regardless of their position on the platform. **(b)** Each fly is housed in a glass tube, with ~25 mm of space for movement. Movement is detected by the interruption of the infra-red light beam and recorded every minute using the DAM system (Trikinetics, Waltham, MA). **(c)** Analysis of the gene ontology terms for Biological process, Molecular function and Cellular component for the genes whose disruption produced hypo- or hyper-arousability phenotypes. We used the GO Term Mapper (http://go.princeton.edu/cgi-bin/GOTermMapper) to bin individual GO categories into more general terms, to facilitate analysis. We represent a GO term enrichment in our screen results compared to the relative weight of that GO term in the whole genome. To do this, we calculated the % of hits in our screen that had each GO term and normalized to the % of genes in the whole genome that contain that term. The data are represented on a log10 scale.

**Extended Data Figure 2. CCHa1 and CCHa1R mutants and RNAi lines. (a)** elav-Gal4-driven RNAi against CCHa1 (elav>CCHa1-RNAi) or its receptor (elav>CCHa1R-RNAi) increases arousal in awake flies. elav>CCHa1-RNAi also causes fragmented sleep (number of sleep episodes is slightly increased, while their duration is decreased); elav>CCHa1R-RNAi decreases total sleep duration due to a decrease in sleep bout length. The last panel shows sleep amount throughout the day, in 30-minute intervals. The white box at the top indicates day period (lights on) while black box indicates night period (lights off). **(b)** qRT-PCR shows that elav>CCHa1-RNAi depletes CCHa1 in the whole fly while it does not affect CCHa2. Each dot represents a sample of 2 whole flies. CCHa1R but not CCHa2R is depleted in fly brains when CCHa1R-RNAi is driven by elav-Gal4. Each dot represents a sample of 5 brains. **(c)** Arousal during sleep, total sleep duration, arousal during wakefulness, and basal locomotor activity data for additional CCHa1- and CCHa1R-RNAi lines. **(d)** qRT-PCR shows that elav>CCHa1-RNAi#2 depletes CCHa1 in the whole fly while it does not affect CCHa2. Each dot represents a sample of 2 whole flies. CCHa1R but not CCHa2R is depleted in fly brains when CCHa1R-RNAi#2 is driven by elav-Gal4. Each dot represents a sample of 5 brains. **(e)**Mutants for CCHa1 or its receptor increase arousability during wakefulness and reduce total sleep amount. The last panel shows sleep amount throughout the day in 30-minute intervals. The white box at the top indicates day period (lights on), the black box indicates night period (lights off). **(f)** Conditional, adult-specific knockdown of CCHa1 or its receptor was achieved using a temperature-sensitive allele of Gal80, a Gal4 inhibitor (elav-Gal4;Tub-Gal80^ts^). Flies were raised at 21 °C, temperature at which Gal80^ts^ is stable and interferes with Gal4 function. Two days after eclosion, temperature was raised to 29 °C, temperature at which Gal80^ts^ is degraded and the Gal4 driver becomes functional. Conditional knockdown phenocopied the results seen with elav-Gal4. In all panels, mean and S.E.M. are depicted; sample sizes and statistical analyses are in Extended Data Table 2.

**Extended Data Figure 3. CCHa1 produced in the gut regulates arousability but not sleep duration. (a)** Mi{MIC}CCHa1^MI09190^ contains the GFP open reading frame inserted downstream of the CCHa1 5’ UTR. Heterozygotes for the Mi{MIC}CCHa1^MI09190^ construct show a GFP pattern of expression that overlaps with the CCHa1 antibody. Homozygotes for Mi{MIC}CCHa1^MI09190^ mutation show the same GFP expression pattern as heterozygotes but do not show any CCHa1 antibody signal since both copies of the gene are mutated. **(b)** nls::GFP expression pattern driven by EECG-Gal4 and pros-Gal4 in the central nervous system shows no co-localization with antibodies against CCHa1. **(c)** CCHa1 in the gut co-localizes with prospero, a marker of enteroendocrine cells. **(d)** elav-Gal4-driven nlsLacZ shows that the driver labels CCHa1-producing enteroendocrine cells in the posterior midgut. **(e)** CCHa1 signal in the gut is abolished when CCHa1-RNAi is expressed with elav-Gal4, EECG-Gal4, or pros-Gal4:TubGal80^ts^. Rectangles indicate the regions shown in Fig. 2a. (**f)** When CCHa1 is knocked down using EECG-Gal4, responsiveness in awake flies is increased but there are no changes in basal locomotion, sleep duration, sleep bout number and length. The last panel shows sleep amount throughout the day, in 30-minute intervals. The white box at the top indicates day period (lights on) while black box indicates night period (lights off). **(g)** When CCHa1 is knocked down using pros-Gal4:TubGal80^ts^, responsiveness in awake flies is increased but there are no changes in basal locomotion, sleep duration, sleep bout number and length. The last panel shows sleep amount throughout the day, in 30-minute intervals. The white box at the top indicates day period (lights on) while black box indicates night period (lights off)**. (h)** Knocking down CCHa1 in the gut does not affect the feeding behavior of flies as measured with blue dye. Each dot represents the average of three experiments, each done with 10 flies. In all panels, mean and S.E.M. are depicted; sample sizes and statistical analyses are in Extended Data Table 2. Scale bars: 100 μm.

**Extended Data Figure 4. Enteroendocrine cells (labeled with EECG-Gal4 driver) regulate arousability through CCHa1. (a)**Inactivation of CCHa1-expressing cells in the gut (EECG>shi^ts^) increases responsiveness in awake flies but does not affect basal locomotion, sleep duration, sleep bout number and length. The last panel shows sleep amount throughout the day, in 30-minute intervals. The white box at the top indicates day period (lights on) while black box indicates night period (lights off). **(b)** Activation of CCHa1-expressing cells in the gut (EECG>TrpA1) decreases responsiveness in awake animals in a CCHa1-dependent manner, and decreases sleep independently of CCHa1. The experiment needs to be done at least at 25 °C to ensure CCHa1-RNAi expression (see methods). The last panel shows sleep amount throughout the day, in 30-minute intervals. The white box at the top indicates day period (lights on) while black box indicates night period (lights off). The statistical significances between the experimental and its two parental controls at the same temperature are shown in the figure. In all panels, mean and S.E.M. are depicted; sample sizes and statistical analyses are in Extended Data Table 2.

**Extended Data Figure 5. Enteroendocrine cells (labeled with pros-Gal4;TubGal80^ts^ driver) regulate arousability through CCHa1.** Activation of CCHa1-expressing cells in the gut (pros-Gal4;TubGal80^ts^>TrpA1) decreases responsiveness in awake animals in a CCHa1-dependent manner, and decreases sleep. The experiment needs to be done at least at 25 °C to ensure CCHa1-RNAi expression (see methods). The last panel shows sleep amount throughout the day, in 30-minute intervals. The white box at the top indicates day period (lights on) while black box indicates night period (lights off). The statistical significances between the experimental genotypes and their parental controls at the same temperature is shown in the figure. In all panels, mean and S.E.M. are depicted; sample sizes and statistical analyses are in Extended Data Table 2.

**Extended Data Figure 6. Effects of dietary protein on CCHa1 levels, sleep, and arousal. (a)** Supplementing regular food with other sources of sugar (galactose and fructose) or fat (propionic acid or hexanoic acid) does not activate enteroendocrine cells or increase CCHa1 protein levels. Access to peptone-supplemented food leads to CCHa1 increase in the gut. The longer the flies are fed peptone, the higher the CCHa1 levels. **(c)** Supplementation with a mix of amino acids phenocopies the effect of peptone on arousal. Supplementation with individual amino acids or biochemically-defined amino acid groups does not recapitulate the effect of peptone on CCHa1levels in the gut; total amino acid concentration is lower in this case. **(d)** Plot of the same CCHa1 signal in the gut shown in (c), according to the amount of amino acid added. **(e)** Peptone supplementation slightly increases sleep, decreases responsiveness to stimulation during wakefulness and slightly reduces basal locomotion. Sleep bout number or length are not affected by peptone supplementation. The last panel shows sleep amount throughout the day in 30-minute intervals, The white box at the top indicates day period (lights on) while black box indicates night period (lights off). **(f)** Supplementing regular food with peptone does not affect the feeding behavior of flies, as reported by quantification of blue dye ingested with the food. Each dot represents the average of three experiments, each done with 10 flies. **(g)**Knocking down CCHa1 specifically in the gut with EECG-Gal4 or pros-Gal4:TubGal80^ts^ diminishes the peptone-induced changes in arousal during wakefulness. In all panels, mean and S.E.M. are depicted; sample sizes and statistical analyses are in Extended Data Table 2.

**Extended Data Figure 7. CCHa1 regulates arousal through dopaminergic neurons in the brain. (a)** From the indicated Gal4 drivers, only elav-Gal4 produces a phenotype when used to knock down CCHa1 receptor. **(b)** Responsiveness in awake flies, basal locomotor activity and total daily sleep duration when the CCHa1 receptor is knocked down with Ddc-Gal4 or PAM-Gal4. Two different Gal4s expressed under the CCHa1R regulatory elements (P{GMR23H07-GAL4}attP2 and TI{2A-Gal4}CCHa1-R2A-Gal4) show expression in the PAM cluster of dopaminergic neurons. **(d)** PAM^MB441B^ driver shows no expression in the gut. **(e)** Knocking down CCHa1 receptor in the PAM^MB441B^ dopaminergic neurons does not affect arousal during wakefulness, sleep duration, sleep bout duration or sleep bout number, but it reduces basal locomotion. The last panel shows sleep amount throughout the day, in 30-minute intervals. The white box at the top indicates day period (lights on) while black box indicates night period (lights off). **(f)** Knocking down CCHa1R in PAM^MB441B^ neurons does not affect feeding behavior as reported by quantification of blue dye ingested with the food. Each dot represents the average of three experiments, each done with 10 flies. **(g)** Knocking down CCHa1R in PAM^MB441B^ neurons does not alter the peptone-induced changes in arousal during wakefulness. In all panels, mean and S.E.M. are depicted; sample sizes and statistical analyses are in Extended Data Table 2.

**Extended Data Figure 8. PAM^MB441B^ neurons regulate arousal through dopamine. (a)** Activation or inhibition of PAM^MB441B^ neurons does not affect arousability during wakefulness, sleep duration, basal locomotion, sleep bout number or sleep bout length. The last panel shows sleep amount throughout the day, in 30-minute intervals. The white box at the top indicates day period (lights on) while black box indicates night period (lights off). **(b)** Knocking down TH in PAM^MB441B^ neurons does not change arousability in awake flies, sleep duration, basal locomotor activity, sleep bout number or sleep bout length. The last panel shows sleep amount throughout the day, in 30-minute intervals. The white box at the top indicates day period (lights on) while black box indicates night period (lights off). In all panels, mean and S.E.M. are depicted; sample sizes and statistical analyses are in Extended Data Table 2.

**Extended Data Figure 9. MBONs integrate information relevant for sleep depth and duration. (a)** Picture of the setup used for thermal stimulation. The percentage of flies that wake up increases as the stimulating temperature goes up (35 vs 40 °C). **(b)** Responsiveness to heating during sleep or wakefulness is unaffected by peptone feeding, CCHa1 knockdown in the gut, or CCHa1R knockdown in PAM^MB441B^ cells. **(c)** Thermal stimulation does not affect the activity of PAM^MB441B^ neurons, as visualized with a calcium-dependent GFP reporter CaLexA. **(d)** MBON^MB083C^-Gal4 driver shows no expression in the fly gut. **(e)**Arousability during wakefulness, sleep duration, basal locomotion or sleep bout length are not altered by changing the activity of MBON^MB083C^ neurons. Inhibition and activation of MBON^MB083C^ slightly increase and decrease sleep bout length, respectively. The last panel shows sleep amount throughout the day, in 30-minute intervals. The white box at the top indicates day period (lights on) while black box indicates night period (lights off). In all panels, mean and S.E.M. are depicted; sample sizes and statistical analyses are in Extended Data Table 2.

**Extended Data Table 1. List of genes from the screen that when knocked down using elav-Gal4 show hypo or hyper-arousable phenotypes**

**Extended Data Table 2. Sample sizes and statistical analysis**

