## Supplementary figures and images for "A gut-secreted peptide controls arousability through modulation of dopaminergic neurons in the brain"

### Extended data figure 1

Extended Data Figure 1

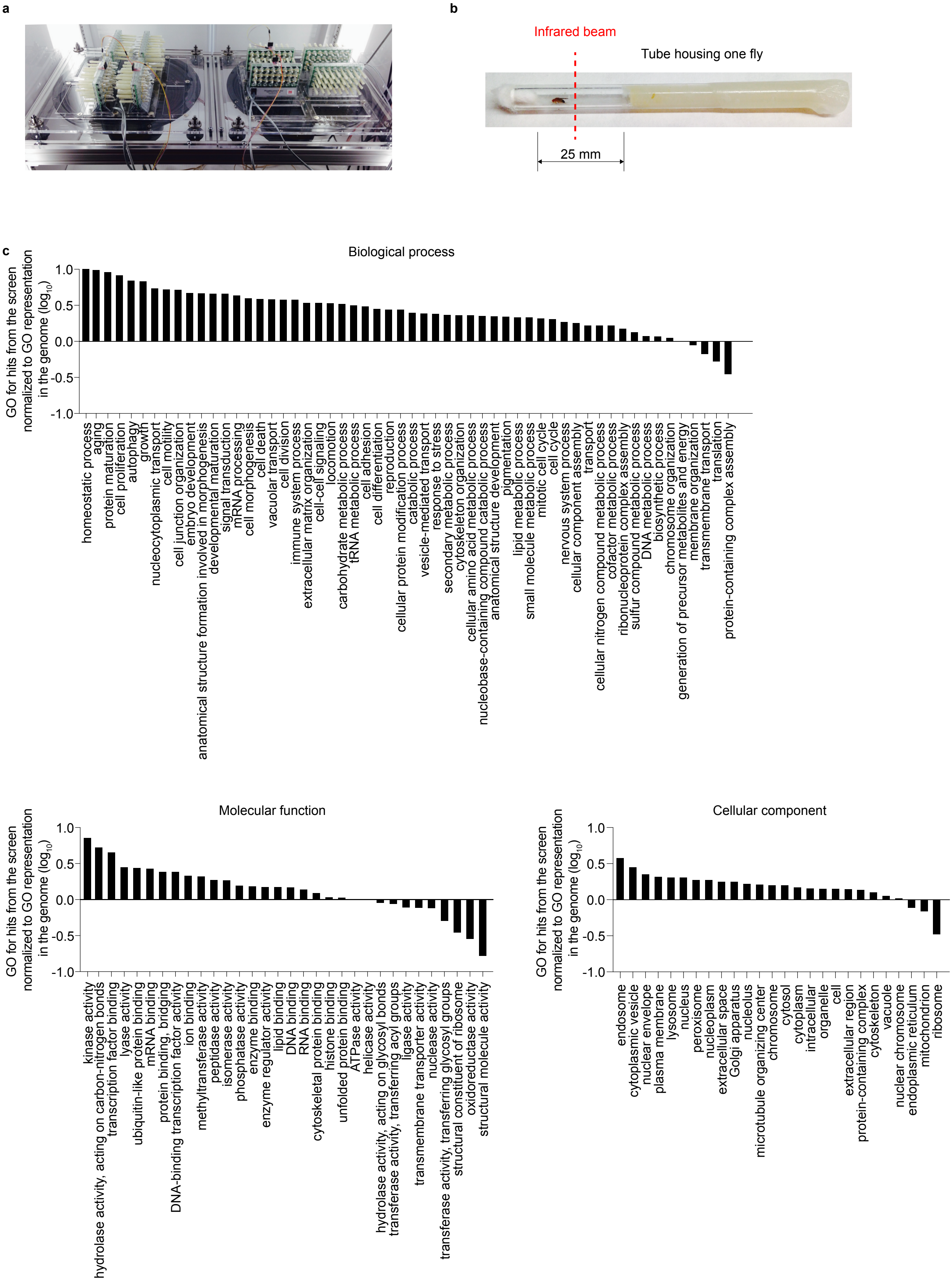

### Extended data figure 2

Extended Data Figure 2

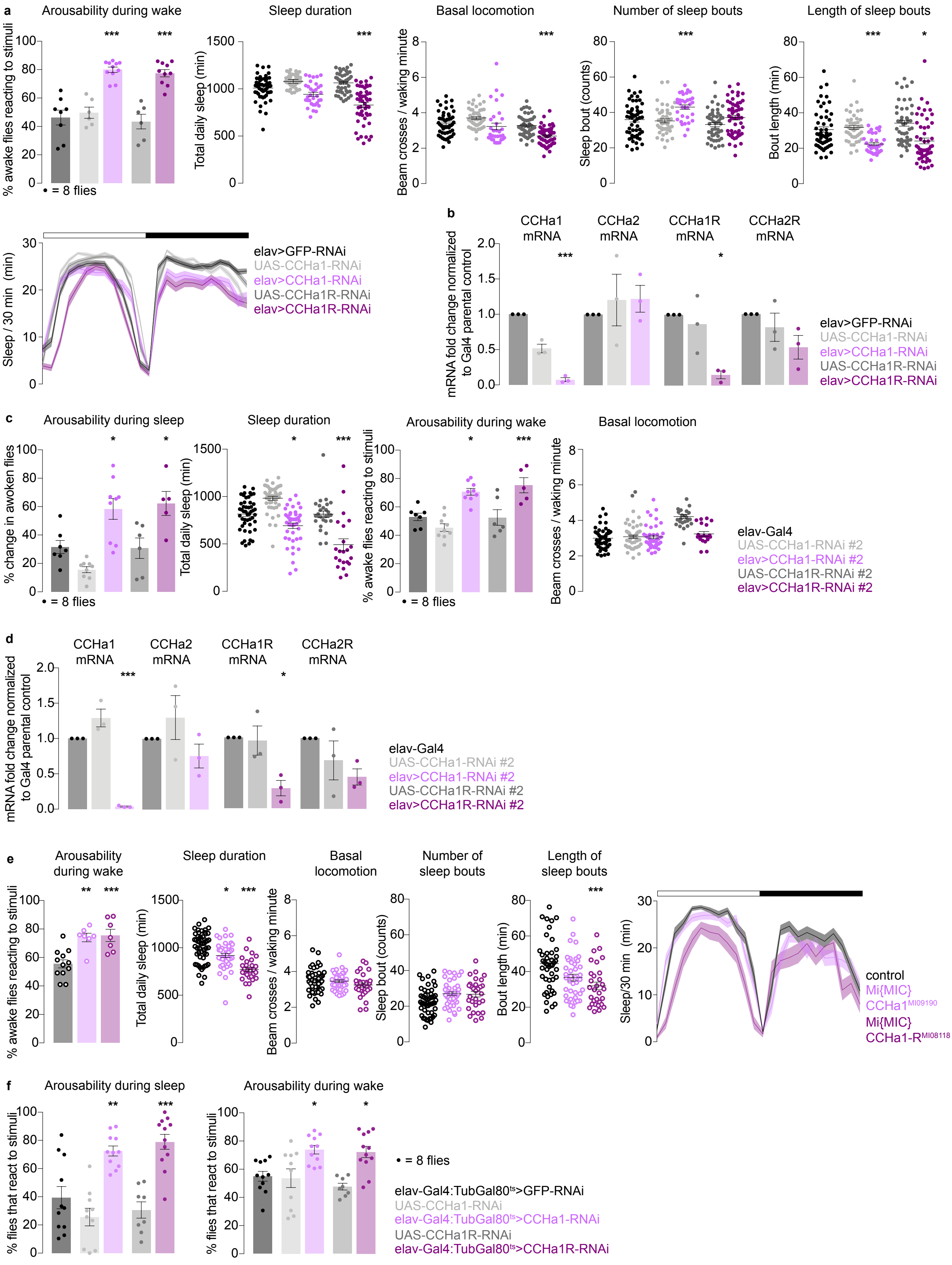

### Extended data figure 3

Extended Data Figure 3

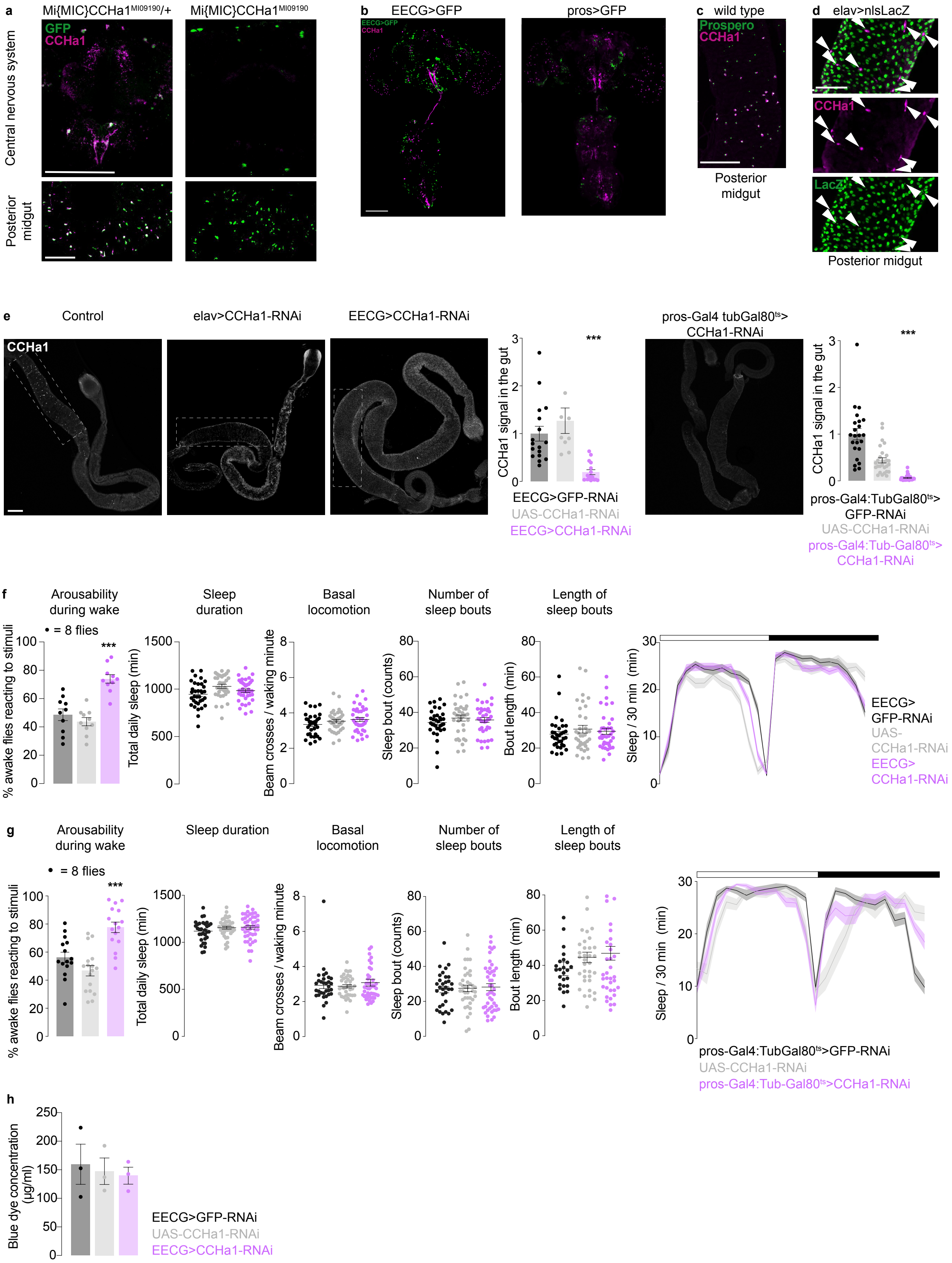

### Extended data figure 4

Extended Data Figure 4

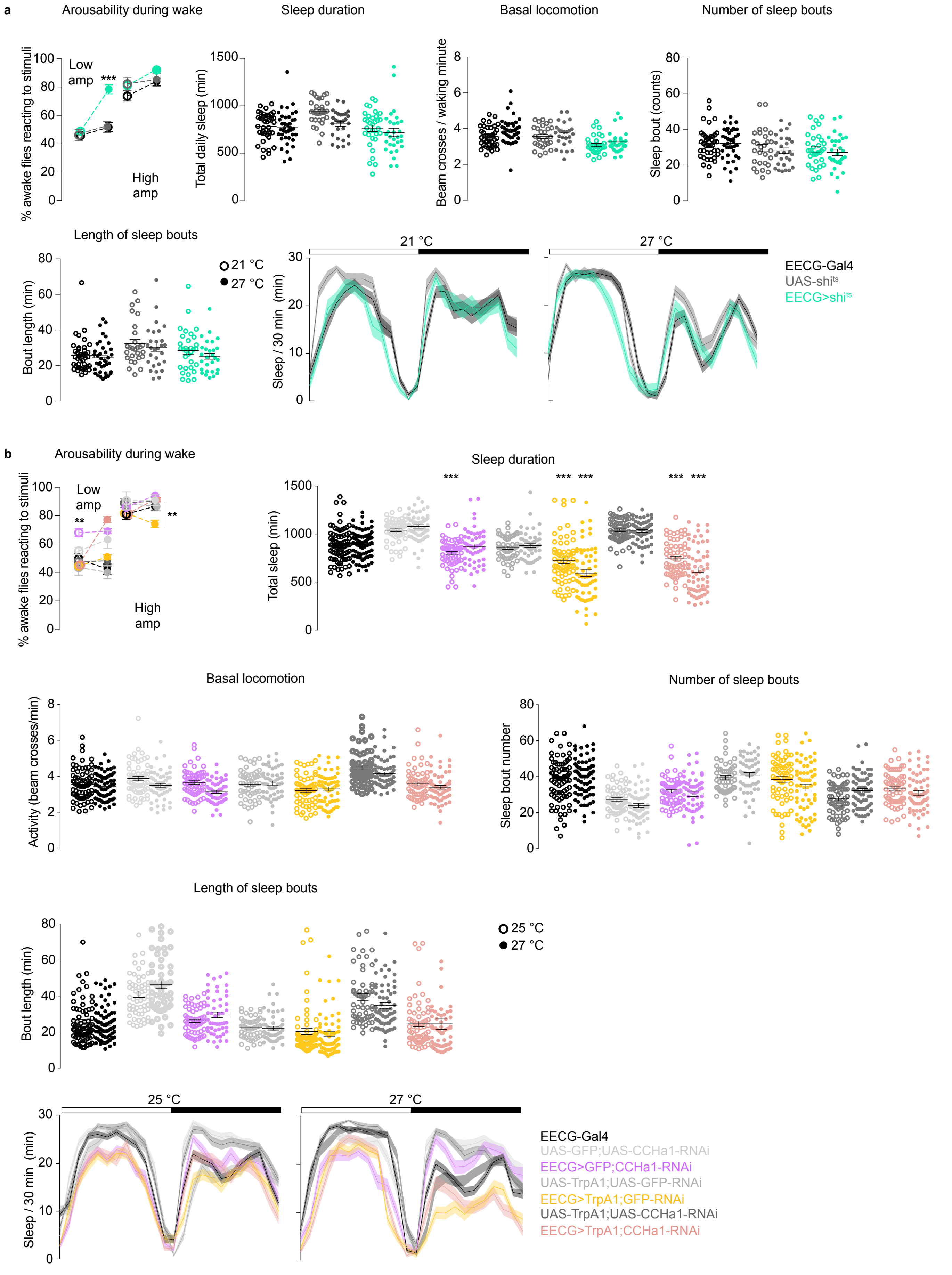

### Extended data figure 5

Extended Data Figure 5

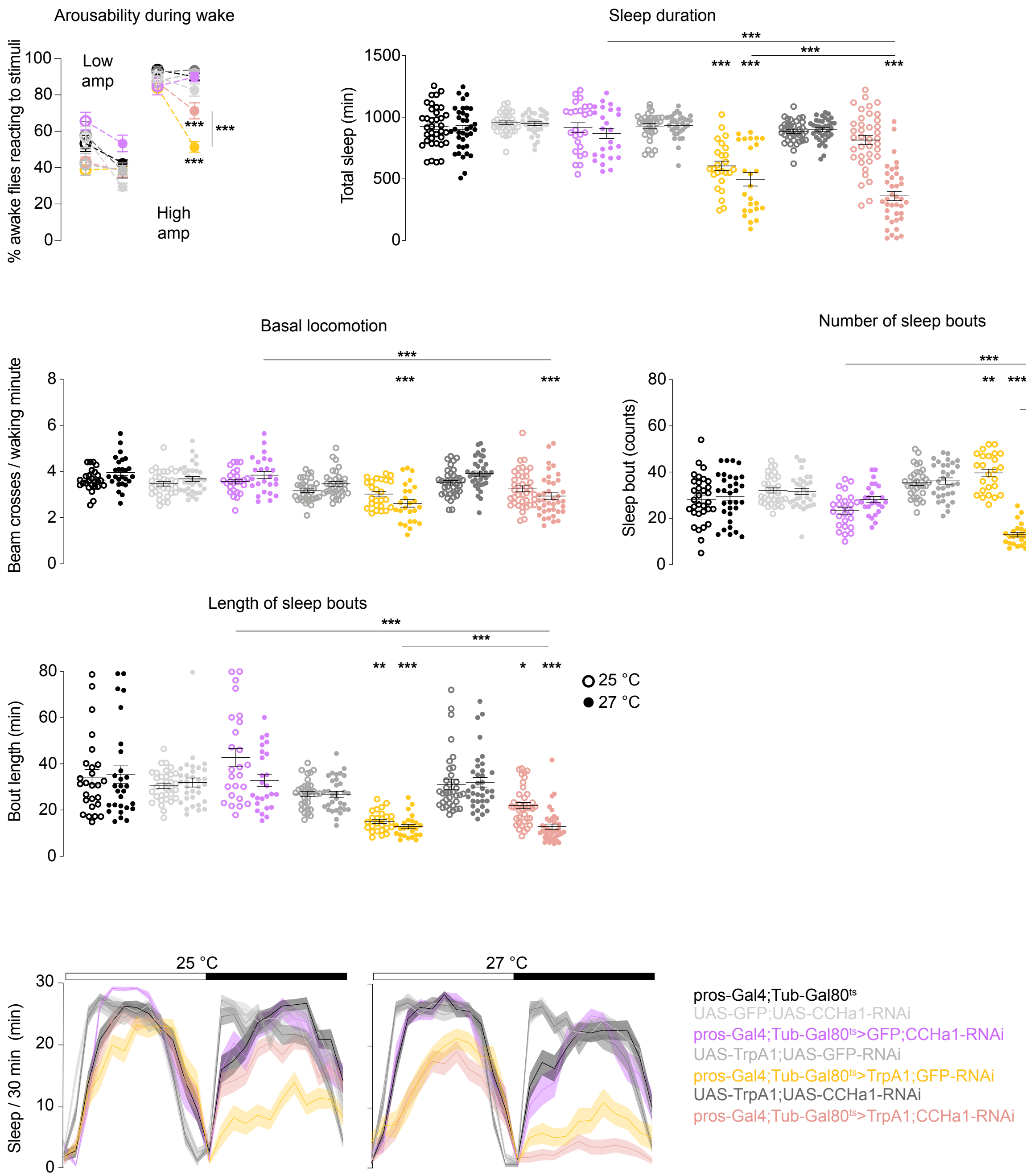

### Extended data figure 6

Extended Data Figure 6

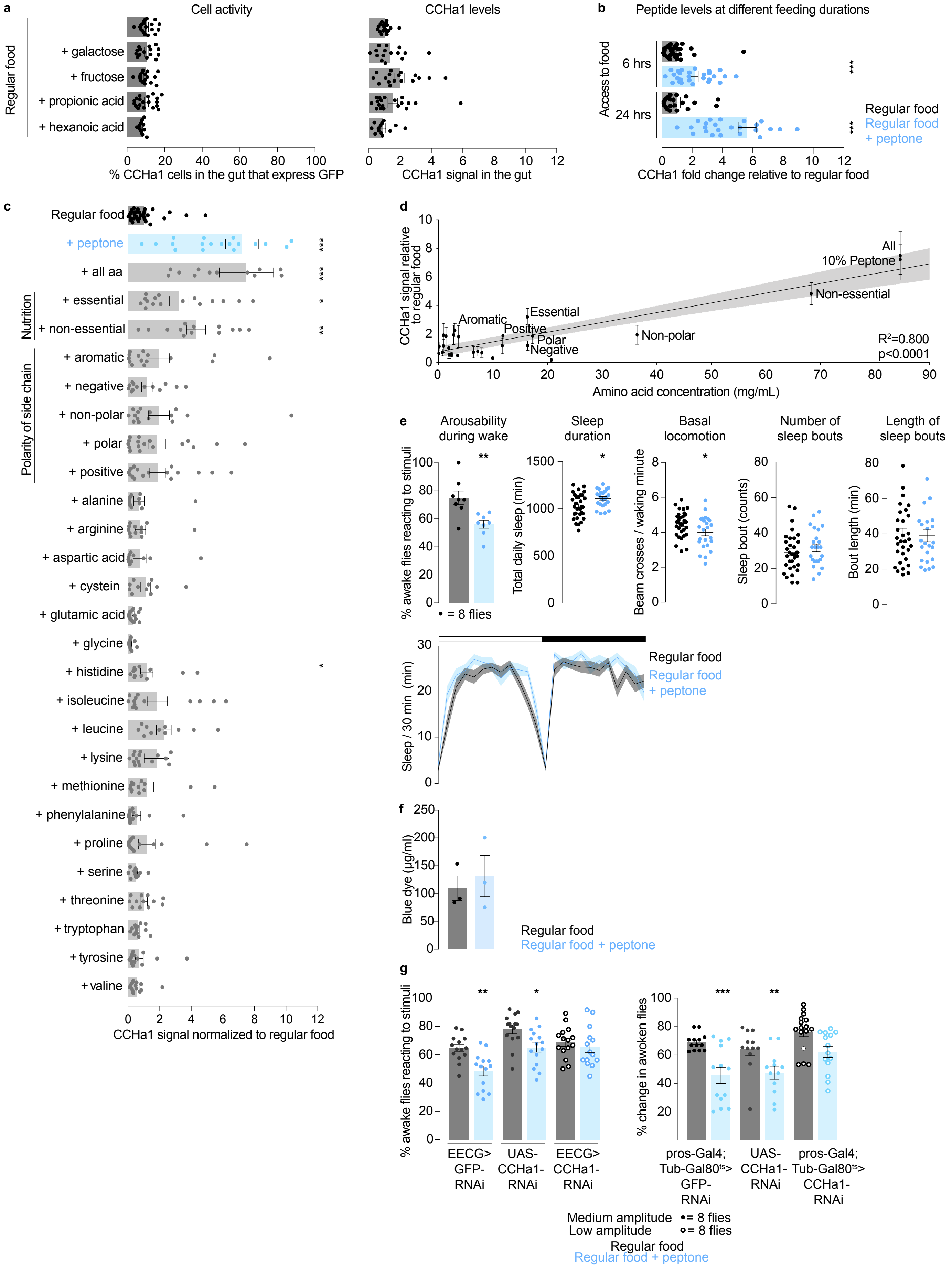

### Extended data figure 7

Extended Data Figure 7

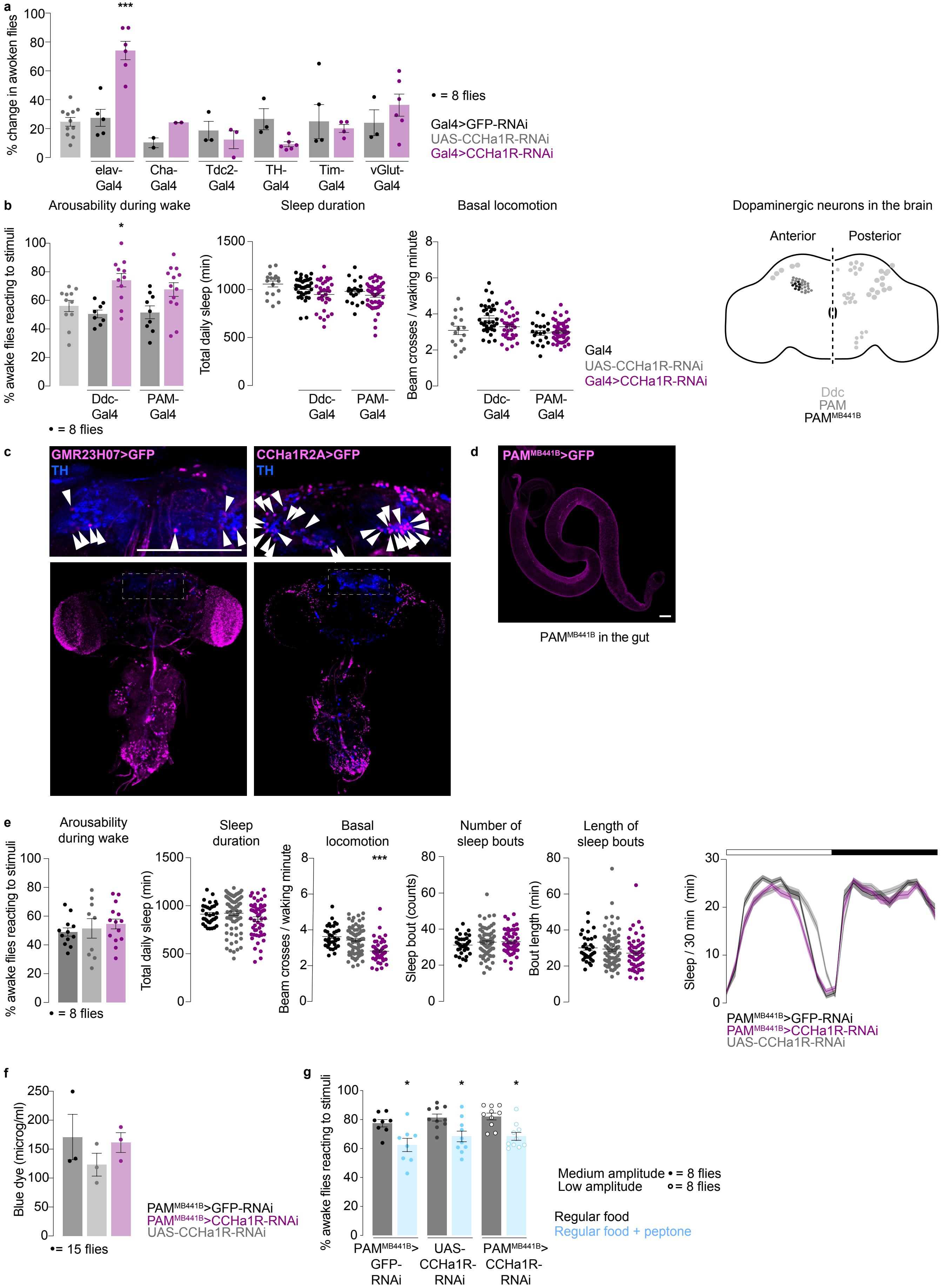

### Extended data figure 8

Extended Data Figure 8

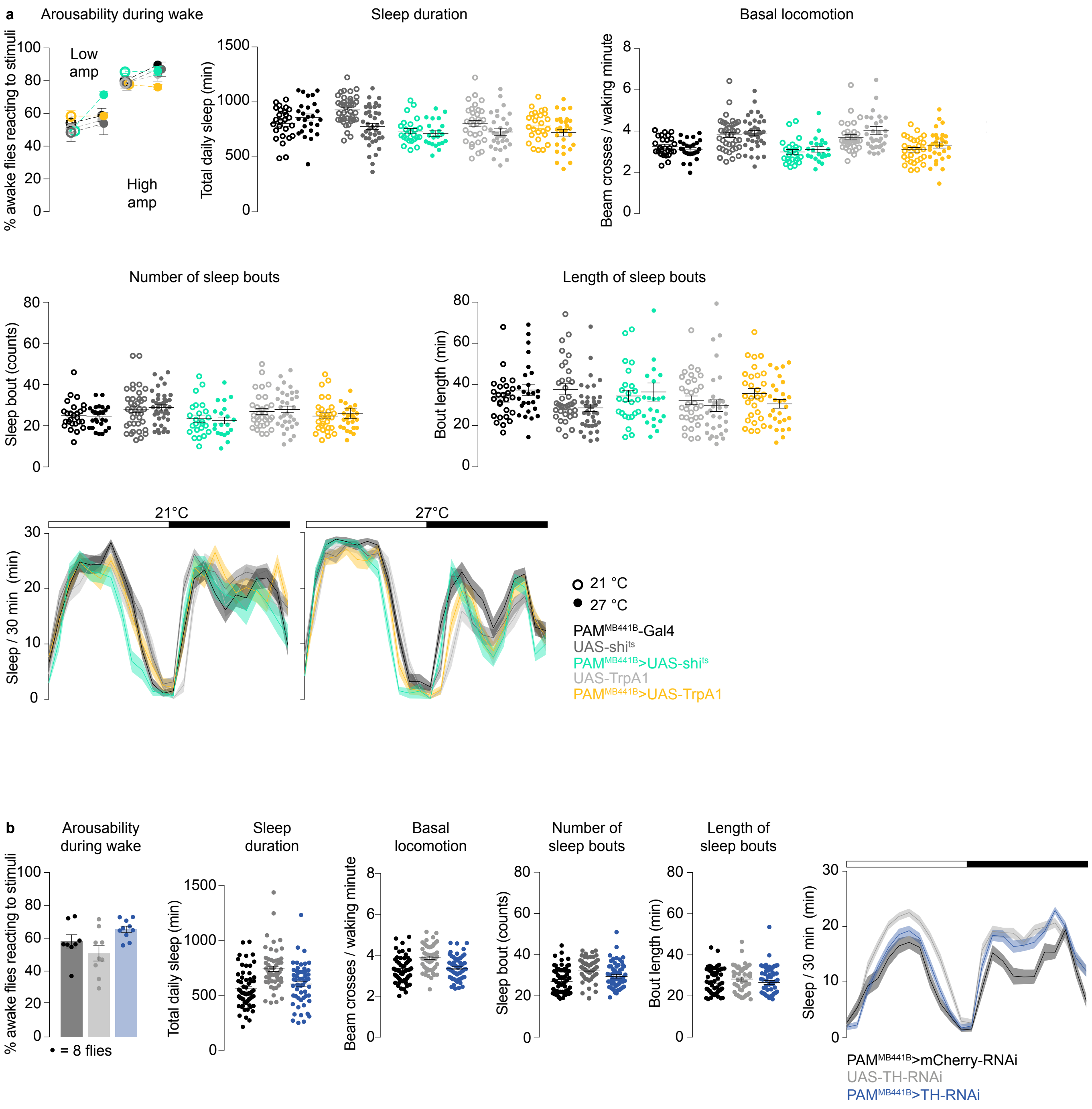

### Extended data figure 9

Extended Data Figure 9

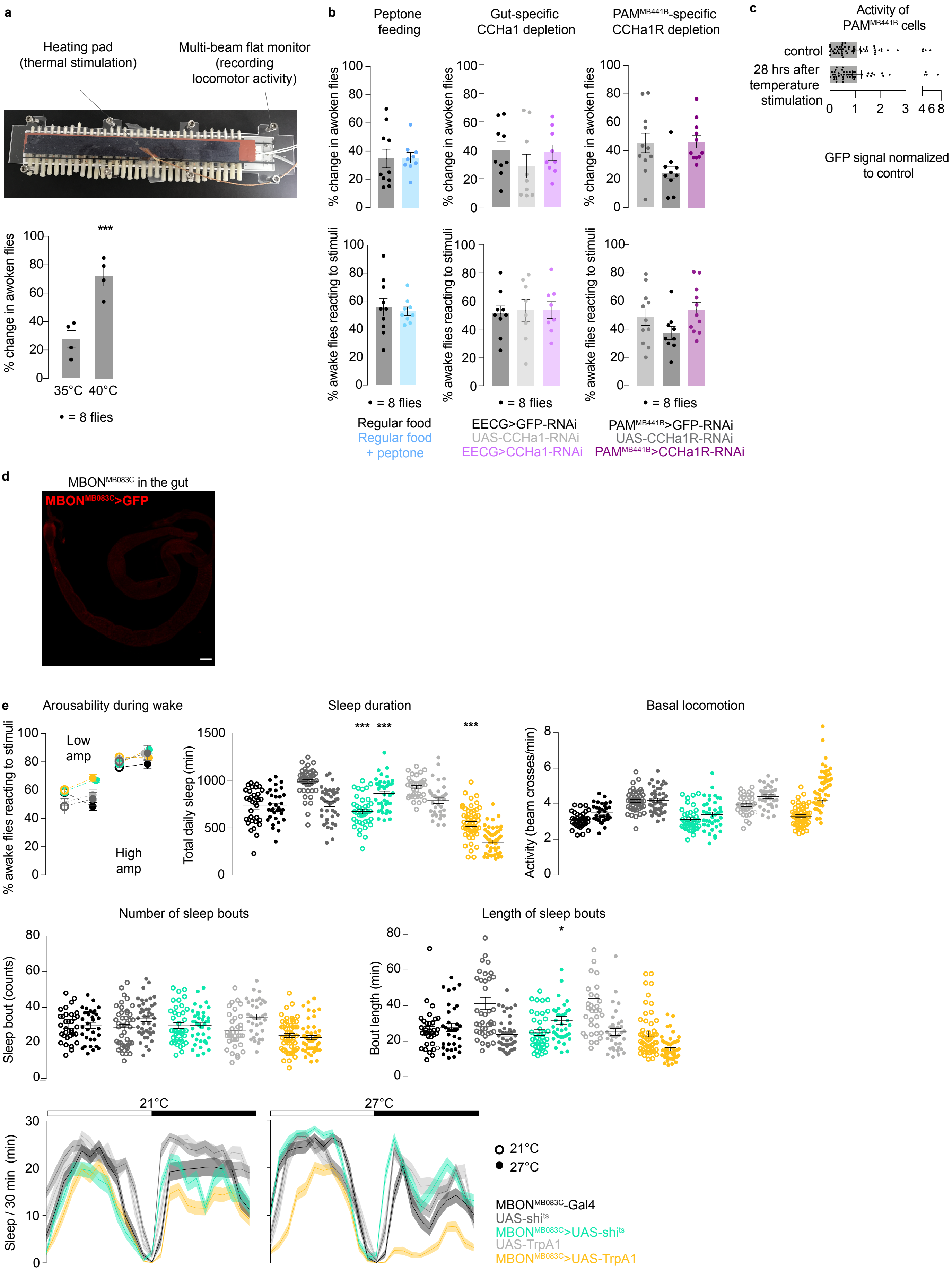
