## Extended data table 1 for "A gut-secreted peptide controls arousability through modulation of dopaminergic neurons in the brain"

**Extended Data Table 1. List of genes from the screen that when knocked down using elav-Gal4 show hypo or hyper-arousable phenotypes**

| Screen hits hyper-arousable |
|---|
| CG10001 |
| CG10002 |
| CG1004 |
| CG10268 |
| CG10377 |
| CG10449 |
| CG10899 |
| CG1098 |
| CG10990 |
| CG11177 |
| CG12069 |
| CG12367 |
| CG12819 |
| CG13575 |
| CG14026 |
| CG14230 |
| CG14358 |
| CG14472 |
| CG14686 |
| CG15497 |
| CG15522 |
| CG16725 |
| CG16901 |
| CG17161 |
| CG17471 |
| CG17686 |
| CG18402 |
| CG18572 |
| CG18660 |
| CG1887 |

| Screen hits hypo-arousable |
|---|
| CG10006 |
| CG1004 |
| CG10143 |
| CG10295 |
| CG10334 |
| CG10388 |
| CG10537 |
| CG10572 |
| CG10579 |
| CG10701 |
| CG10823 |
| CG10850 |
| CG10877 |
| CG10901 |
| CG10907 |
| CG10967 |
| CG10990 |
| CG11331 |
| CG11420 |
| CG11556 |
| CG11937 |
| CG12170 |
| CG12348 |
| CG12386 |
| CG12505 |
| CG12559 |
| CG13204 |
| CG13575 |
| CG13779 |
| CG13784 |

| |
|---|
| CG2087 |
| CG2615 |
| CG2845 |
| CG2846 |
| CG30106 |
| CG3143 |
| CG3171 |
| CG3178 |
| CG32110 |
| CG32139 |
| CG32281 |
| CG32445 |
| CG32498 |
| CG33276 |
| CG34381 |
| CG3593 |
| CG3705 |
| CG42250 |
| CG42341 |
| CG4313 |
| CG4322 |
| CG4385 |
| CG4637 |
| CG4927 |
| CG5216 |
| CG5517 |
| CG5671 |
| CG6054 |
| CG6315 |
| CG6355 |
| CG6438 |
| CG6496 |
| CG6951 |
| CG7437 |
| CG8167 |
| CG8173 |

| |
|---|
| CG14562 |
| CG1464 |
| CG15793 |
| CG1641 |
| CG16725 |
| CG16901 |
| CG17077 |
| CG17239 |
| CG17269 |
| CG1773 |
| CG18455 |
| CG1848 |
| CG2124 |
| CG2210 |
| CG2331 |
| CG2647 |
| CG31000 |
| CG31522 |
| CG3225 |
| CG32425 |
| CG33119 |
| CG3326 |
| CG33467 |
| CG33956 |
| CG3466 |
| CG3613 |
| CG3871 |
| CG3884 |
| CG4016 |
| CG40351 |
| CG42341 |
| CG4320 |
| CG4444 |
| CG4475 |
| CG4703 |
| CG4706 |

| |
|---|
| CG8318 |
| CG8726 |
| CG8950 |
| CG9696 |

| |
|---|
| CG5025 |
| CG5308 |
| CG5432 |
| CG5436 |
| CG5610 |
| CG5813 |
| CG5920 |
| CG5941 |
| CG5992 |
| CG6196 |
| CG6496 |
| CG6498 |
| CG6703 |
| CG6736 |
| CG6854 |
| CG6963 |
| CG7204 |
| CG7281 |
| CG7391 |
| CG7429 |
| CG7599 |
| CG7643 |
| CG7971 |
| CG8172 |
| CG8318 |
| CG8525 |
| CG8795 |
| CG8952 |
| CG8964 |
| CG9310 |
| CG9554 |
| CG9746 |
