## Extended data table 2 for "A gut-secreted peptide controls arousability through modulation of dopaminergic neurons in the brain"

**Extended Data Table 2. Sample sizes and statistical analysis**

| Figure (panel) | Genotype | N | Statistical Test | P value |
|---|---|---|---|---|
| **Figure 1** | | | | |
| 1 (b) | elav>GFP-RNAi vs<br>elav>CCHa1-RNAi | 10<br>12 | Kruskal-Wallis (5,46)=35.03 Dunn's multiple comparisons test | 0.0006 |
| | UAS-CCHa1 RNAi vs<br>elav>CCHa1-RNAi | 8<br>12 | | 0.0052 |
| | elav>GFP-RNAi vs<br>elav>CCHa1R-RNAi | 10<br>10 | | <0.0001 |
| | UAS-CCHA1R-RNAi vs<br>elav>CCHa1R-RNAi | 6<br>10 | | 0.0023 |
| 1 (c) | Control vs<br>Mi{MIC}CCHa1$^{MI09190}$ | 12<br>7 | One-way ANOVA F(2,23)=20.06 Tukey's multiple comparisons test | 0.0001 |
| | Control vs<br>Mi{MIC}CCHa1-R$^{MI08118}$ | 12<br>7 | | <0.0001 |
| **Figure 2** | | | | |
| 2 (b) Arousability from sleep | EECG>GFP-RNAi vs<br>EECG>CCHa1-RNAi | 10<br>10 | One-way ANOVA F(2,27)=51.47 Tukey's multiple comparisons test | <0.0001 |
| | UAS-CCHA1-RNAi vs<br>EECG>CCHa1-RNAi | 10<br>10 | | <0.0001 |
| 2 (b) Arousability from sleep | Pros-Gal4:Tub-Gal80>GFP-RNAi vs<br>Pros-Gal4:Tub-Gal80>CCHa1-RNAi | 15<br>16 | One-way ANOVA F(2,45)=27.90 Tukey's multiple comparisons test | <0.0001 |
| | UAS-CCHA1-RNAi vs<br>Pros-Gal4:Tub-Gal80>CCHa1-RNAi | 17<br>16 | | <0.0001 |
| 2 (c) Low amplitude | EECG-Gal4 21°C vs<br>EECG>shi$^{ts}$ 21°C vs | 10<br>10 | Two-way ANOVA F (2, 48) = 15.84 Tukey's multiple comparisons test | 0.9980 |
| | UAS- shi$^{ts}$ 21°C vs<br>EECG>shi$^{ts}$ 21°C vs | 7<br>10 | | 0.6925 |
| | EECG-Gal4 27°C vs<br>EECG>shi$^{ts}$ 27°C vs | 10<br>10 | | <0.0001 |
| | UAS- shi$^{ts}$ 27°C vs<br>EECG>shi$^{ts}$ 27°C vs | 7<br>10 | | <0.0001 |
| 2 (c) High amplitude | EECG-Gal4 21°C vs<br>EECG>shi$^{ts}$ 21°C vs | 10<br>10 | Two-way ANOVA F (2, 48) = 0.991 Tukey's multiple | 0.5956 |
| | UAS- shi$^{ts}$ 21°C vs<br>EECG>shi$^{ts}$ 21°C vs | 7<br>10 | | 0.9705 |
| | EECG-Gal4 27°C vs | 10<br>10 | | 0.9966 |

| | | | | |
|---|---|---|---|---|
| | EECG>shi$^{ts}$ 27°C vs | | comparisons test | |
| | UAS- shi$^{ts}$ 27°C vs<br>EECG>shi$^{ts}$ 27°C vs | 7<br>10 | | >0.9999 |
| 2 (d) EECG Low amplitude | EECG-Gal4 25°C vs<br>EECG>TrpA1;GFP-RNAi 25°C | 10<br>10 | Two-way ANOVA F (6,110) = 4.295 Tukey's multiple comparisons test | >0.9999 |
| | UAS-TrpA1;UAS-GFP-RNAi 25°C vs<br>EECG>TrpA1;GFP-RNAi 25°C | 10<br>10 | | >0.9999 |
| | EECG-Gal4 25°C vs<br>EECG>TrpA1;CCHa1-RNAi 25°C | 10<br>10 | | 0.5995 |
| | UAS-TrpA1;UAS-CCHa1-RNAi 25°C vs<br>EECG>TrpA1;CCHa1-RNAi 25°C | 10<br>10 | | >0.9999 |
| | EECG>TrpA1;GFP-RNAi 25°C vs<br>EECG>TrpA1;CCHa1-RNAi 25°C | 10<br>10 | | 0.0675 |
| | EECG-Gal4 25°C vs<br>EECG>CCHa1-RNAi;GFP 25°C | 6<br>6 | | 0.0011 |
| | UAS-CCHa1-RNAi;UAS-GFP 25°C vs<br>EECG>CCHa1-RNAi;GFP 25°C | 6<br>6 | | 0.9259 |
| | EECG>CCHa1-RNAi;GFP 25°C vs<br>EECG>TrpA1;CCHa1-RNAi 25°C | 6<br>10 | | 0.8784 |
| | EECG-Gal4 27°C vs<br>EECG>TrpA1;GFP-RNAi 27°C | 10<br>10 | | >0.9999 |
| | UAS-TrpA1;UAS-GFP-RNAi 27°C vs<br>EECG>TrpA1;GFP-RNAi 27°C | 10<br>10 | | >0.9999 |
| | EECG-Gal4 27°C vs<br>EECG>TrpA1;CCHa1-RNAi 27°C | 10<br>10 | | <0.0001 |
| | UAS-TrpA1;UAS-CCHa1-RNAi 27°C vs<br>EECG>TrpA1;CCHa1-RNAi 27°C | 10<br>10 | | <0.0001 |
| | EECG>TrpA1;GFP-RNAi 27°C vs<br>EECG>TrpA1;CCHa1-RNAi 27°C | 10<br>10 | | <0.0001 |
| | EECG-Gal4 27°C vs<br>EECG>CCHa1-RNAi;GFP 27°C | 6<br>6 | | >0.9999 |
| | UAS-CCHa1-RNAi;UAS-GFP 27°C vs<br>EECG>CCHa1-RNAi;GFP 27°C | 6<br>6 | | 0.0411 |
| | EECG>CCHa1-RNAi;GFP 27°C vs<br>EECG>TrpA1;CCHa1-RNAi 27°C | 6<br>10 | | >0.9999 |
| 2 (d) EECG High amplitude | EECG-Gal4 25°C vs<br>EECG>TrpA1;GFP-RNAi 25°C | 10<br>10 | Two-way ANOVA F (6,110) = 10.74 Tukey's multiple comparisons test | 0.0020 |
| | UAS-TrpA1;UAS-GFP-RNAi 25°C vs<br>EECG>TrpA1;GFP-RNAi 25°C | 10<br>10 | | 0.0011 |
| | EECG-Gal4 25°C vs<br>EECG>TrpA1;CCHa1-RNAi 25°C | 10<br>10 | | >0.9999 |
| | UAS-TrpA1;UAS-CCHa1-RNAi 25°C vs<br>EECG>TrpA1;CCHa1-RNAi 25°C | 10<br>10 | | >0.9999 |
| | EECG>TrpA1;GFP-RNAi 25°C vs<br>EECG>TrpA1;CCHa1-RNAi 25°C | 10<br>10 | | 0.0311 |
| | EECG-Gal4 25°C vs<br>EECG>CCHa1-RNAi;GFP 25°C | 6<br>6 | | >0.9999 |

| | | | | |
|---|---|---|---|---|
| | UAS-CCHa1-RNAi;UAS-GFP 25°C vs EECG>CCHa1-RNAi;GFP 25°C | 6<br>6 | | >0.9999 |
| | EECG>CCHa1-RNAi;GFP 25°C vs EECG>TrpA1;CCHa1-RNAi 25°C | 6<br>10 | | >0.9999 |
| | EECG-Gal4 27°C vs EECG>TrpA1;GFP-RNAi 27°C | 10<br>10 | | <0.0001 |
| | UAS-TrpA1;UAS-GFP-RNAi 27°C vs EECG>TrpA1;GFP-RNAi 27°C | 10<br>10 | | <0.0001 |
| | EECG-Gal4 27°C vs EECG>TrpA1;CCHa1-RNAi 27°C | 10<br>10 | | >0.9999 |
| | UAS-TrpA1;UAS-CCHa1-RNAi 27°C vs EECG>TrpA1;CCHa1-RNAi 27°C | 10<br>10 | | >0.9999 |
| | EECG>TrpA1;GFP-RNAi 27°C vs EECG>TrpA1;CCHa1-RNAi 27°C | 10<br>10 | | <0.0001 |
| | EECG-Gal4 27°C vs EECG>CCHa1-RNAi;GFP 27°C | 6<br>6 | | >0.9999 |
| | UAS-CCHa1-RNAi;UAS-GFP 27°C vs EECG>CCHa1-RNAi;GFP 27°C | 6<br>6 | | >0.9999 |
| | EECG>CCHa1-RNAi;GFP 27°C vs EECG>TrpA1;CCHa1-RNAi 27°C | 6<br>10 | | >0.9999 |
| 2 (d) Pros-Gal4:Tub-Gal80$^{ts}$ Low amplitude | Pros-Gal4:Tub-Gal80$^{ts}$ -Gal4 25°C vs Pros-Gal4:Tub-Gal80$^{ts}$ >TrpA1;GFP-RNAi 25°C | 6<br>6 | Two-way ANOVA F (6,81) = 1.025 Tukey's multiple comparisons test | >0.9999 |
| | UAS-TrpA1;UAS-GFP-RNAi 25°C vs Pros-Gal4:Tub-Gal80$^{ts}$ >TrpA1;GFP-RNAi 25°C | 5<br>6 | | >0.9999 |
| | Pros-Gal4:Tub-Gal80$^{ts}$ -Gal4 25°C vs Pros-Gal4:Tub-Gal80$^{ts}$ >TrpA1;CCHa1-RNAi 25°C | 6<br>6 | | >0.9999 |
| | UAS-TrpA1;UAS-CCHa1-RNAi 25°C vs Pros-Gal4:Tub-Gal80$^{ts}$ >TrpA1;CCHa1-RNAi 25°C | 5<br>6 | | 0.4552 |
| | Pros-Gal4:Tub-Gal80$^{ts}$ >TrpA1;GFP-RNAi 25°C vs Pros-Gal4:Tub-Gal80$^{ts}$ >TrpA1;CCHa1-RNAi 25°C | 6<br>6 | | >0.9999 |
| | Pros-Gal4:Tub-Gal80$^{ts}$ -Gal4 25°C vs Pros-Gal4:Tub-Gal80$^{ts}$ >CCHa1-RNAi;GFP 25°C | 6<br>6 | | 0.5854 |
| | UAS-CCHa1-RNAi;UAS-GFP 25°C vs Pros-Gal4:Tub-Gal80$^{ts}$ >CCHa1-RNAi;GFP 25°C | 6<br>6 | | >0.9999 |
| | Pros-Gal4:Tub-Gal80$^{ts}$ >CCHa1-RNAi;GFP 25°C vs Pros-Gal4:Tub-Gal80$^{ts}$ >TrpA1;CCHa1-RNAi 25°C | 6<br>6 | | 0.0053 |
| | Pros-Gal4:Tub-Gal80$^{ts}$ -Gal4 27°C vs | 8<br>9 | | >0.9999 |

| | | | | |
|---|---|---|---|---|
| | Pros-Gal4:Tub-Gal80$^{ts}$ >TrpA1;GFP-RNAi 27°C | | | |
| | UAS-TrpA1;UAS-GFP-RNAi 27°C vs Pros-Gal4:Tub-Gal80$^{ts}$ >TrpA1;GFP-RNAi 27°C | 8<br>9 | | >0.9999 |
| | Pros-Gal4:Tub-Gal80$^{ts}$ -Gal4 27°C vs Pros-Gal4:Tub-Gal80$^{ts}$ >TrpA1;CCHa1-RNAi 27°C | 8<br>8 | | >0.9999 |
| | UAS-TrpA1;UAS-CCHa1-RNAi 27°C vs Pros-Gal4:Tub-Gal80$^{ts}$ >TrpA1;CCHa1-RNAi 27°C | 6<br>8 | | 0.9995 |
| | Pros-Gal4:Tub-Gal80$^{ts}$ >TrpA1;GFP-RNAi 27°C vs Pros-Gal4:Tub-Gal80$^{ts}$ >TrpA1;CCHa1-RNAi 27°C | 9<br>8 | | >0.9999 |
| | Pros-Gal4:Tub-Gal80$^{ts}$ -Gal4 27°C vs Pros-Gal4:Tub-Gal80$^{ts}$ >CCHa1-RNAi;GFP 27°C | 8<br>10 | | <0.0001 |
| | UAS-CCHa1-RNAi;UAS-GFP 27°C vs Pros-Gal4:Tub-Gal80$^{ts}$ >CCHa1-RNAi;GFP 27°C | 6<br>10 | | 0.2883 |
| | Pros-Gal4:Tub-Gal80$^{ts}$ >CCHa1-RNAi;GFP 27°C vs Pros-Gal4:Tub-Gal80$^{ts}$ >TrpA1;CCHa1-RNAi 27°C | 10<br>8 | | <0.0001 |
| 2 (d) Pros-Gal4:Tub-Gal80$^{ts}$ High amplitude | Pros-Gal4:Tub-Gal80$^{ts}$ -Gal4 25°C vs Pros-Gal4:Tub-Gal80$^{ts}$ >TrpA1;GFP-RNAi 25°C | 6<br>6 | Two-way ANOVA $F(6,81) = 0.9661$ Tukey's multiple comparisons test | >0.9999 |
| | UAS-TrpA1;UAS-GFP-RNAi 25°C vs Pros-Gal4:Tub-Gal80$^{ts}$ >TrpA1;GFP-RNAi 25°C | 5<br>6 | | >0.9999 |
| | Pros-Gal4:Tub-Gal80$^{ts}$ -Gal4 25°C vs Pros-Gal4:Tub-Gal80$^{ts}$ >TrpA1;CCHa1-RNAi 25°C | 6<br>6 | | >0.9999 |
| | UAS-TrpA1;UAS-CCHa1-RNAi 25°C vs Pros-Gal4:Tub-Gal80$^{ts}$ >TrpA1;CCHa1-RNAi 25°C | 5<br>6 | | >0.9999 |
| | Pros-Gal4:Tub-Gal80$^{ts}$ >TrpA1;GFP-RNAi 25°C vs Pros-Gal4:Tub-Gal80$^{ts}$ >TrpA1;CCHa1-RNAi 25°C | 6<br>6 | | >0.9999 |
| | Pros-Gal4:Tub-Gal80$^{ts}$ -Gal4 25°C vs Pros-Gal4:Tub-Gal80$^{ts}$ >CCHa1-RNAi;GFP 25°C | 6<br>6 | | >0.9999 |
| | UAS-CCHa1-RNAi;UAS-GFP 25°C vs Pros-Gal4:Tub-Gal80$^{ts}$ >CCHa1-RNAi;GFP 25°C | 6<br>6 | | >0.9999 |

| | | | | |
|---|---|---|---|---|
| | Pros-Gal4:Tub-Gal80$^{ts}$ >CCHa1-RNAi;GFP 25°C vs<br>Pros-Gal4:Tub-Gal80$^{ts}$ >TrpA1;CCHa1-RNAi 25°C | 6<br>6 | | >0.9999 |
| | Pros-Gal4:Tub-Gal80$^{ts}$ -Gal4 27°C vs<br>Pros-Gal4:Tub-Gal80$^{ts}$ >TrpA1;GFP-RNAi 27°C | 8<br>9 | | <0.0001 |
| | UAS-TrpA1;UAS-GFP-RNAi 27°C vs<br>Pros-Gal4:Tub-Gal80$^{ts}$ >TrpA1;GFP-RNAi 27°C | 8<br>9 | | <0.0001 |
| | Pros-Gal4:Tub-Gal80$^{ts}$ -Gal4 27°C vs<br>Pros-Gal4:Tub-Gal80$^{ts}$ >TrpA1;CCHa1-RNAi 27°C | 8<br>8 | | 0.0569 |
| | UAS-TrpA1;UAS-CCHa1-RNAi 27°C vs<br>Pros-Gal4:Tub-Gal80$^{ts}$ >TrpA1;CCHa1-RNAi 27°C | 6<br>8 | | 0.0200 |
| | Pros-Gal4:Tub-Gal80$^{ts}$ >TrpA1;GFP-RNAi 27°C vs<br>Pros-Gal4:Tub-Gal80$^{ts}$ >TrpA1;CCHa1-RNAi 27°C | 9<br>8 | | 0.0002 |
| | Pros-Gal4:Tub-Gal80$^{ts}$ -Gal4 27°C vs<br>Pros-Gal4:Tub-Gal80$^{ts}$ >CCHa1-RNAi;GFP 27°C | 8<br>10 | | >0.9999 |
| | UAS-CCHa1-RNAi;UAS-GFP 27°C vs<br>Pros-Gal4:Tub-Gal80$^{ts}$ >CCHa1-RNAi;GFP 27°C | 6<br>10 | | >0.9999 |
| | Pros-Gal4:Tub-Gal80$^{ts}$ >CCHa1-RNAi;GFP 27°C vs<br>Pros-Gal4:Tub-Gal80$^{ts}$ >TrpA1;CCHa1-RNAi 27°C | 10<br>8 | | 0.1367 |
| **Figure 3** | | | | |
| 3 (c) Cell activity | Regular vs<br>Regular + glucose | 21<br>13 | One-way ANOVA<br>F (3, 70) = 113.1<br>Tukey's multiple comparisons test | 0.9347 |
| | Regular vs<br>Regular + coconut oil | 21<br>17 | | 0.6456 |
| | Regular vs<br>Regular + peptone | 21<br>23 | | <0.0001 |
| 3 (c) CCHa1 levels | Regular vs<br>Regular + glucose | 21<br>13 | One-way ANOVA<br>F (3, 70) = 25.87<br>Tukey's multiple comparisons test | 0.1636 |
| | Regular vs<br>Regular + coconut oil | 21<br>17 | | 0.9584 |
| | Regular vs<br>Regular + peptone | 21<br>23 | | <0.0001 |
| 3 (c) mRNA levels | Regular vs<br>Regular + glucose | 3<br>3 | One-way ANOVA | 0.9972 |

| | | | | |
|---|---|---|---|---|
| | Regular vs<br>Regular + coconut oil | 3<br>3 | F (3,8) = 7.66<br>Tukey's multiple comparisons test | 0.8024 |
| | Regular vs<br>Regular + peptone | 3<br>3 | | 0.0344 |
| 3 (d) | UAS-CCHa1-RNAi regular vs<br>UAS-CCHa1-RNAi peptone | 14<br>14 | T-test<br>t=5.76 df=14 | <0.0001 |
| 3 (e) EECG-Gal4 | EECG>GFP-RNAi regular vs<br>EECG>GFP-RNAi peptone | 14<br>14 | Two-way ANOVA<br>F (2,79) = 6.041<br>Tukey's multiple comparisons test | 0.0001 |
| | UAS-CCHa1-RNAi regular vs<br>UAS-CCHa1-RNAi peptone | 14<br>14 | | <0.0001 |
| | EECG>CCHa1-RNAi regular vs<br>EECG>CCHa1-RNAi peptone | 15<br>14 | | 0.7444 |
| 3 (e) pros-Gal4:TubGal80$^{ts}$ | Pros-Gal4:Tub-Gal80$^{ts}$>GFP-RNAi regular vs<br>Pros-Gal4:Tub-Gal80$^{ts}$>GFP-RNAi peptone | 13<br>13 | Two-way ANOVA<br>F (2,74) = 5.953<br>Tukey's multiple comparisons test | <0.0001 |
| | UAS-CCHa1-RNAi regular vs<br>UAS-CCHa1-RNAi peptone | 12<br>12 | | 0.0234 |
| | Pros-Gal4:Tub-Gal80$^{ts}$ >CCHa1-RNAi regular vs<br>Pros-Gal4:Tub-Gal80$^{ts}$ >CCHa1-RNAi peptone | 16<br>14 | | 0.4720 |
| **Figure 4** | | | | |
| 4 (a) | UAS-CCHa1R-RNAi vs<br>Ddc>CCHa1R-RNAi | 11<br>11 | One-way ANOVA<br>F (4,47) = 16.31<br>Tukey's multiple comparisons test | <0.0001 |
| | Ddc>GFP-RNAi vs<br>Ddc>CCHa1R-RNAi | 8<br>11 | | <0.0001 |
| | UAS-CCHa1R-RNAi vs<br>PAM>CCHa1R-RNAi | 11<br>13 | | 0.0064 |
| | PAM>GFP-RNAi vs<br>PAM>CCHa1R-RNAi | 9<br>13 | | 0.0004 |
| 4 (b) | PAM$^{MB441B}$>GFP-RNAi vs<br>PAM$^{MB441B}$>CCHa1R-RNAi | 12<br>14 | One-way ANOVA<br>F (4,32) = 14.28<br>Tukey's multiple comparisons test | <0.0001 |
| | UAS-CCHa1R-RNAi vs<br>PAM$^{MB441B}$>CCHa1R-RNAi | 10<br>14 | | <0.0001 |
| 4 (c) | PAM$^{MB441B}$>CalexA regular vs<br>PAM$^{MB441B}$>CalexA peptone | 58<br>57 | Two-way ANOVA<br>F (1,198) = 6.489<br>Tukey's multiple comparisons test | 0.0048 |
| | PAM$^{MB441B}$>CalexA;CCHa1R-RNAi regular vs<br>PAM$^{MB441B}$>CalexA;CCHa1R-RNAi peptone | 46<br>47 | | 0.9436 |

| | | | | |
|---|---|---|---|---|
| 4 (d) | $PAM^{MB441B}$>GFP-RNAi regular vs $PAM^{MB441B}$>GFP-RNAi peptone | 8<br>8 | Two-way ANOVA F (2,50) = 8.037 Tukey's multiple comparisons test | <0.0001 |
| | UAS-CCHa1R-RNAi regular vs UAS-CCHa1R-RNAi peptone | 10<br>10 | | 0.0008 |
| | $PAM^{MB441B}$>CCHa1R-RNAi regular vs $PAM^{MB441B}$>CCHa1R-RNAi peptone | 10<br>10 | | 0.9913 |
| 4 (e) Low amplitude | $PAM^{MB441B}$-Gal4 21°C vs $PAM^{MB441B}$>TrpA1 21°C | 10<br>10 | Two-way ANOVA F (4,78) = 29.06 Tukey's multiple comparisons test | 0.9810 |
| | UAS-TrpA1 21°C vs PAMMB441B>TrpA1 21°C | 8<br>10 | | 0.7663 |
| | $PAM^{MB441B}$-Gal4 21°C vs $PAM^{MB441B}$>$shi^{ts}$ 21°C | 10<br>10 | | >0.9999 |
| | UAS- $shi^{ts}$ 21°C vs $PAM^{MB441B}$>$shi^{ts}$ 21°C | 6<br>10 | | 0.4820 |
| | $PAM^{MB441B}$-Gal4 27°C vs $PAM^{MB441B}$>TrpA1 27°C | 10<br>10 | | 0.4339 |
| | UAS-TrpA1 27°C vs $PAM^{MB441B}$>TrpA1 27°C | 8<br>10 | | 0.2073 |
| | $PAM^{MB441B}$-Gal4 27°C vs $PAM^{MB441B}$>$shi^{ts}$ 27°C | 10<br>10 | | <0.0001 |
| | UAS- $shi^{ts}$ 27°C vs $PAM^{MB441B}$>$shi^{ts}$ 27°C | 6<br>10 | | <0.0001 |
| 4 (e) High amplitude | $PAM^{MB441B}$-Gal4 21°C vs $PAM^{MB441B}$>TrpA1 21°C | 10<br>10 | Two-way ANOVA F (4,78) = 2.457 Tukey's multiple comparisons test | 0.9935 |
| | UAS-TrpA1 21°C vs $PAM^{MB441B}$>TrpA1 21°C | 8<br>10 | | >0.9999 |
| | $PAM^{MB441B}$-Gal4 21°C vs $PAM^{MB441B}$>$shi^{ts}$ 21°C | 10<br>10 | | 0.9554 |
| | UAS- $shi^{ts}$ 21°C vs $PAM^{MB441B}$>$shi^{ts}$ 21°C | 6<br>10 | | >0.9999 |
| | $PAM^{MB441B}$-Gal4 27°C vs $PAM^{MB441B}$>TrpA1 27°C | 10<br>10 | | 0.0003 |
| | UAS-TrpA1 27°C vs $PAM^{MB441B}$>TrpA1 27°C | 8<br>10 | | 0.0341 |
| | $PAM^{MB441B}$-Gal4 27°C vs $PAM^{MB441B}$>$shi^{ts}$ 27°C | 10<br>10 | | 0.9916 |
| | UAS- $shi^{ts}$ 27°C vs $PAM^{MB441B}$>$shi^{ts}$ 27°C | 6<br>10 | | 0.9376 |
| 4 (f) | $PAM^{MB441B}$>mCherry-RNAi;dcr2 vs $PAM^{MB441B}$>TH-RNAi;dcr2 | 8<br>10 | One-way ANOVA F (4,24) = 12.39 Tukey's multiple comparisons test | 0.0023 |
| | UAS-TH-RNAi;UAS-dcr2 vs $PAM^{MB441B}$>TH-RNAi;dcr | 9<br>10 | | 0.0003 |

| | | | | |
|---|---|---|---|---|
| 4 (g) | $PAM^{MB441B}$>CaLexA no vibrations vs $PAM^{MB441B}$>CaLexA 4h after vibrations | 68<br>41 | Kruskal-Wallis F (3,143) = 29.95 Dunn's multiple comparisons test | >0.9999 |
| | $PAM^{MB441B}$>CaLexA no vibrations vs $PAM^{MB441B}$>CaLexA 28h after vibrations | 68<br>34 | | <0.0001 |
| 4 (h) Low amplitude | $MBON^{MB083C}$-Gal4 21°C vs $MBON^{MB083C}$ >TrpA1 21°C | 10<br>10 | Two-way ANOVA F (4,78) = 1.901 Tukey's multiple comparisons test | >0.9999 |
| | UAS-TrpA1 21°C vs $MBON^{MB083C}$ >TrpA1 21°C | 8<br>10 | | >0.9999 |
| | $MBON^{MB083C}$ -Gal4 21°C vs $MBON^{MB083C}$ >$shi^{ts}$ 21°C | 10<br>10 | | 0.8372 |
| | UAS- $shi^{ts}$ 21°C vs $MBON^{MB083C}$ >$shi^{ts}$ 21°C | 6<br>10 | | 0.8798 |
| | $MBON^{MB083C}$ -Gal4 27°C vs $MBON^{MB083C}$ >TrpA1 27°C | 10<br>10 | | 0.9992 |
| | UAS-TrpA1 27°C vs $MBON^{MB083C}$ >TrpA1 27°C | 8<br>10 | | >0.9999 |
| | $MBON^{MB083C}$ -Gal4 27°C vs $MBON^{MB083C}$ >$shi^{ts}$ 27°C | 10<br>10 | | <0.0001 |
| | UAS- $shi^{ts}$ 27°C vs $MBON^{MB083C}$ >$shi^{ts}$ 27°C | 6<br>10 | | 0.0113 |
| 4 (h) High amplitude | $MBON^{MB083C}$ -Gal4 21°C vs $MBON^{MB083C}$ >TrpA1 21°C | 10<br>10 | Two-way ANOVA F (4,78) = 6.841 Tukey's multiple comparisons test | 0.9770 |
| | UAS-TrpA1 21°C vs $MBON^{MB083C}$ >TrpA1 21°C | 8<br>10 | | 0.9946 |
| | $MBON^{MB083C}$ -Gal4 21°C vs $MBON^{MB083C}$ >$shi^{ts}$ 21°C | 10<br>10 | | >0.9999 |
| | UAS- $shi^{ts}$ 21°C vs $MBON^{MB083C}$ >$shi^{ts}$ 21°C | 6<br>10 | | 0.9952 |
| | $MBON^{MB083C}$ -Gal4 27°C vs $MBON^{MB083C}$ >TrpA1 27°C | 10<br>10 | | <0.0001 |
| | UAS-TrpA1 27°C vs $MBON^{MB083C}$ >TrpA1 27°C | 8<br>10 | | 0.0002 |
| | $MBON^{MB083C}$ -Gal4 27°C vs $MBON^{MB083C}$ >$shi^{ts}$ 27°C | 10<br>10 | | >0.9999 |
| | UAS- $shi^{ts}$ 27°C vs $MBON^{MB083C}$ >$shi^{ts}$ 27°C | 6<br>10 | | >0.9999 |
| **Extended Data Figure 2** | | | | |
| 2 (a) Arousability during wake | elav>GFP-RNAi vs elav>CCHa1-RNAi | 10<br>12 | One-way ANOVA F(4,41)=30.27 Tukey's multiple | <0.0001 |
| | UAS-CCHa1 RNAi vs elav>CCHa1-RNAi | 8<br>12 | | <0.0001 |
| | elav>GFP-RNAi vs elav>CCHa1R-RNAi | 10<br>10 | | <0.0001 |
| | UAS-CCHA1R-RNAi vs | 6 | | <0.0001 |

| | | | | |
|---|---|---|---|---|
| | elav>CCHa1R-RNAi | 10 | comparisons test | |
| 2 (a)<br>Sleep duration | elav>GFP-RNAi vs<br>elav>CCHa1-RNAi | 53<br>32 | One-way ANOVA F(4,226)=31 Tukey's multiple comparisons test | 0.1387 |
| | UAS-CCHa1 RNAi vs<br>elav>CCHa1-RNAi | 47<br>32 | | <0.0001 |
| | elav>GFP-RNAi vs<br>elav>CCHa1R-RNAi | 53<br>52 | | <0.0001 |
| | UAS-CCHA1R-RNAi vs<br>elav>CCHa1R-RNAi | 47<br>52 | | <0.0001 |
| 2 (a)<br>Basal locomotion | elav>GFP-RNAi vs<br>elav>CCHa1-RNAi | 53<br>32 | One-way ANOVA F(4,226)=15.8 Tukey's multiple comparisons test | 0.8440 |
| | UAS-CCHa1 RNAi vs<br>elav>CCHa1-RNAi | 47<br>32 | | 0.0152 |
| | elav>GFP-RNAi vs<br>elav>CCHa1R-RNAi | 53<br>52 | | <0.0001 |
| | UAS-CCHa1R-RNAi vs<br>elav>CCHa1R-RNAi | 47<br>52 | | 0.0001 |
| 2 (a) Number of sleep bouts | elav>GFP-RNAi vs<br>elav>CCHa1-RNAi | 53<br>32 | One-way ANOVA F(4,229)=6.82 Tukey's multiple comparisons test | 0.0024 |
| | UAS-CCHa1 RNAi vs<br>elav>CCHa1-RNAi | 47<br>32 | | 0.0009 |
| | elav>GFP-RNAi vs<br>elav>CCHa1R-RNAi | 53<br>52 | | 0.9563 |
| | UAS-CCHA1R-RNAi vs<br>elav>CCHa1R-RNAi | 47<br>52 | | 0.1586 |
| 2 (a) Length of sleep bout | elav>GFP-RNAi vs<br>elav>CCHa1-RNAi | 53<br>32 | One-way ANOVA F(4,229)=10.65 Tukey's multiple comparisons test | 0.0037 |
| | UAS-CCHa1 RNAi vs<br>elav>CCHa1-RNAi | 47<br>32 | | 0.0007 |
| | elav>GFP-RNAi vs<br>elav>CCHa1R-RNAi | 53<br>52 | | 0.0145 |
| | UAS-CCHA1R-RNAi vs<br>elav>CCHa1R-RNAi | 47<br>52 | | <0.0001 |
| 2 (b) CCHa1 mRNA | elav>GFP-RNAi vs<br>elav>CCHa1-RNAi | 3<br>3 | One-way ANOVA F(2,6)=139.6 Tukey's multiple comparisons test | <0.0001 |
| | UAS-CCHa1 RNAi vs<br>elav>CCHa1-RNAi | 3<br>3 | | 0.0005 |
| 2 (b) CCHa2 mRNA | elav>GFP-RNAi vs<br>elav>CCHa1-RNAi | 3<br>3 | One-way ANOVA F(2,6)=0.274 Tukey's multiple comparisons test | 0.7915 |
| | UAS-CCHa1 RNAi vs<br>elav>CCHa1-RNAi | 3<br>3 | | 0.9987 |
| 2 (b) CCHa1R mRNA | elav>GFP-RNAi vs<br>elav>CCHa1R-RNAi | 3<br>3 | One-way ANOVA | 0.0255 |

| | | | | |
|---|---|---|---|---|
| | UAS-CCHa1R RNAi vs<br>elav>CCHa1R-RNAi | 3<br>3 | F(2,6)=10.75<br>Tukey's multiple comparisons test | 0.0119 |
| 2 (b) CCHa2R mRNA | elav>GFP-RNAi vs<br>elav>CCHa1R-RNAi | 3<br>3 | One-way ANOVA<br>F(2,6)=2.448<br>Tukey's multiple comparisons test | 0.1504 |
| | UAS-CCHa1R-RNAi vs<br>elav>CCHa1R-RNAi | 3<br>3 | | 0.4308 |
| 2 (c) Arousability during sleep | elav-Gal4 vs<br>elav>CCHa1-RNAi #2 | 7<br>9 | One-way ANOVA<br>F(4,31)=11.35<br>Tukey's multiple comparisons test | 0.0185 |
| | UAS-CCHa1 RNAi vs<br>elav>CCHa1-RNAi #2 | 9<br>9 | | <0.0001 |
| | elav-Gal4 vs<br>elav>CCHa1R-RNAi #2 | 7<br>6 | | 0.0213 |
| | UAS-CCHA1R-RNAi vs<br>elav>CCHa1R-RNAi #2 | 6<br>5 | | 0.0227 |
| 2 (c) Sleep duration | elav-Gal4 vs<br>elav>CCHa1-RNAi #2 | 47<br>45 | One-way ANOVA<br>F(4,175)=29.82<br>Tukey's multiple comparisons test | 0.0146 |
| | UAS-CCHa1 RNAi vs<br>elav>CCHa1-RNAi #2 | 44<br>45 | | <0.0001 |
| | elav-Gal4 vs<br>elav>CCHa1R-RNAi #2 | 47<br>19 | | <0.0001 |
| | UAS-CCHA1R-RNAi vs<br>elav>CCHa1R-RNAi #2 | 25<br>19 | | <0.0001 |
| 2 (c) Arousability during wake | elav-Gal4 vs<br>elav>CCHa1-RNAi #2 | 7<br>9 | One-way ANOVA<br>F(4,31)=13.49<br>Tukey's multiple comparisons test | 0.0061 |
| | UAS-CCHa1 RNAi vs<br>elav>CCHa1-RNAi #2 | 9<br>9 | | <0.0001 |
| | elav-Gal4 vs<br>elav>CCHa1R-RNAi #2 | 7<br>6 | | 0.0026 |
| | UAS-CCHA1R-RNAi vs<br>elav>CCHa1R-RNAi #2 | 6<br>5 | | 0.0029 |
| 2 (c) Basal locomotion | elav-Gal4 vs<br>elav>CCHa1-RNAi #2 | 47<br>45 | Kruskal-Wallis (5,5)=44.21<br>Dunn's multiple comparisons test | >0.9999 |
| | UAS-CCHa1 RNAi vs<br>elav>CCHa1-RNAi #2 | 44<br>45 | | >0.9999 |
| | elav-Gal4 vs<br>elav>CCHa1R-RNAi #2 | 47<br>19 | | >0.9999 |
| | UAS-CCHA1R-RNAi vs<br>elav>CCHa1R-RNAi #2 | 25<br>19 | | 0.0033 |
| 2 (d) CCHa1 mRNA | elav-Gal4 vs<br>elav>CCHa1-RNAi #2 | 3<br>3 | One-way ANOVA<br>F(2,6)=81.92<br>Tukey's multiple | 0.0002 |
| | UAS-CCHa1 RNAi vs<br>elav>CCHa1-RNAi #2 | 3<br>3 | | <0.0001 |

| | | | | |
|---|---|---|---|---|
| | | | comparisons test | |
| 2 (d) CCHa2 mRNA | elav-Gal4 vs elav>CCHa1-RNAi #2 | 3<br>3 | One-way ANOVA F(2,6)=1.786 Tukey's multiple comparisons test | 0.6922 |
| | UAS-CCHa1 RNAi vs elav>CCHa1-RNAi #2 | 3<br>3 | | 0.2225 |
| 2 (d) CCHa1R mRNA | elav-Gal4 vs elav>CCHa1-RNAi #2 | 3<br>3 | One-way ANOVA F(2,6)=10.75 Tukey's multiple comparisons test | 0.0222 |
| | UAS-CCHa1 RNAi vs elav>CCHa1-RNAi #2 | 3<br>3 | | 0.0288 |
| 2 (d) CCHa2R mRNA | elav-Gal4 vs elav>CCHa1-RNAi #2 | 3<br>3 | One-way ANOVA F(2,6)=2.523 Tukey's multiple comparisons test | 0.1425 |
| | UAS-CCHa1 RNAi vs elav>CCHa1-RNAi #2 | 3<br>3 | | 0.6254 |
| 2 (e) Arousability during wake | Control vs Mi{MIC}CCHa1$^{MI09190}$ | 12<br>7 | One-way ANOVA F(2,23)=12.33 Tukey's multiple comparisons test | 0.0018 |
| | Control vs Mi{MIC}CCHa1-R$^{MI08118}$ | 12<br>7 | | 0.0008 |
| 2 (e) Sleep duration | Control vs Mi{MIC}CCHa1$^{MI09190}$ | 52<br>41 | One-way ANOVA F(2,120)=24.79 Tukey's multiple comparisons test | 0.0144 |
| | Control vs Mi{MIC}CCHa1-R$^{MI08118}$ | 52<br>30 | | <0.0001 |
| 2 (e) Basal locomotion | Control vs Mi{MIC}CCHa1$^{MI09190}$ | 52<br>41 | One-way ANOVA F(2,120)=1.731 Tukey's multiple comparisons test | 0.9858 |
| | Control vs Mi{MIC}CCHa1-R$^{MI08118}$ | 52<br>30 | | 0.1878 |
| 2 (e) Number of sleep bouts | Control vs Mi{MIC}CCHa1$^{MI09190}$ | 52<br>41 | One-way ANOVA | 0.0150 |

| | | | | |
|---|---|---|---|---|
| | Control vs<br>Mi{MIC}CCHa1-R$^{MI08118}$ | 52<br>30 | F(2,120)=5.005<br>Tukey's multiple comparisons test | 0.0431 |
| 2 (e) Length of sleep bouts | Control vs<br>Mi{MIC}CCHa1$^{MI09190}$ | 52<br>41 | Kruskal-Wallis (3,120)=13.82<br>Dunn's multiple comparisons test | 0.0703 |
| | Control vs<br>Mi{MIC}CCHa1-R$^{MI08118}$ | 52<br>30 | | 0.0008 |
| 2 (f) Arousability during sleep | elav-Gal4 tubGal80$^{ts}$> GFP-RNAi vs<br>elav-Gal4 tubGal80$^{ts}$>CCHa1-RNAi | 11<br>11 | One-way ANOVA<br>F(4,47)=17.45<br>Tukey's multiple comparisons test | 0.0017 |
| | UAS-CCHa1-RNAi vs<br>Elav-Gal4 tubGal80$^{ts}$>CCHa1-RNAi | 10<br>11 | | <0.0001 |
| | elav-Gal4 tubGal80$^{ts}$> GFP-RNAi vs<br>elav-Gal4 tubGal80$^{ts}$>CCHa1R-RNAi | 11<br>12 | | <0.0001 |
| | UAS-CCHa1R-RNAi vs<br>Elav-Gal4 tubGal80$^{ts}$>CCHa1R-RNAi | 8<br>12 | | <0.0001 |
| 2 (f) Arousability during wakefulness | elav-Gal4 tubGal80$^{ts}$> GFP-RNAi vs<br>elav-Gal4 tubGal80$^{ts}$>CCHa1-RNAi | 11<br>11 | One-way ANOVA<br>F(4,47)=7.7<br>Tukey's multiple comparisons test | 0.0170 |
| | UAS-CCHa1-RNAi vs<br>Elav-Gal4 tubGal80$^{ts}$>CCHa1-RNAi | 10<br>11 | | <0.0001 |
| | elav-Gal4 tubGal80$^{ts}$> GFP-RNAi vs<br>elav-Gal4 tubGal80$^{ts}$>CCHa1R-RNAi | 11<br>12 | | 0.0116 |
| | UAS-CCHa1R-RNAi vs<br>Elav-Gal4 tubGal80$^{ts}$>CCHa1R-RNAi | 8<br>12 | | 0.0022 |
| **Extended Data Figure 3** | | | | |
| 3 (e) | EECG>GFP-RNAi vs<br>EECG>CCHa1-RNAi | 17<br>16 | Kruskal-Wallis<br>F(3,42)=25.56<br>Dunn's multiple comparisons test | <0.0001 |
| | UAS-CCHA1-RNAi vs<br>EECG>CCHa1-RNAi | 9<br>16 | | <0.0001 |
| 3 (e) | Pros-Gal4:Tub-Gal80$^{ts}$>GFP-RNAi vs<br>Pros-Gal4:Tub-Gal80$^{ts}$ >CCHa1-RNAi | 23<br>32 | Kruskal-Wallis<br>F(3,83)=61.67<br>Dunn's multiple comparisons test | <0.0001 |
| | UAS-CCHA1-RNAi vs<br>Pros-Gal4:Tub-Gal80$^{ts}$ >CCHa1-RNAi | 28<br>32 | | <0.0001 |
| 3 (f) Arousability during wake | EECG>GFP-RNAi vs<br>EECG>CCHa1-RNAi | 10<br>10 | One-way ANOVA | <0.0001 |
| | UAS-CCHA1-RNAi vs | 10 | | <0.0001 |

| | | | | |
|---|---|---|---|---|
| | EECG>CCHa1-RNAi | 10 | F(2,27)=23.17 Tukey's multiple comparisons test | |
| 3 (f) Sleep duration | EECG>GFP-RNAi vs EECG>CCHa1-RNAi | 33 38 | One-way ANOVA F (2,103) = 4.068 Tukey's multiple comparisons test | 0.3739 |
| | UAS-CCHA1-RNAi vs EECG>CCHa1-RNAi | 35 38 | | 0.2620 |
| 3 (f) Basal locomotion | EECG>GFP-RNAi vs EECG>CCHa1-RNAi | 33 38 | One-way ANOVA F (2,103) = 1.742 Tukey's multiple comparisons test | 0.1597 |
| | UAS-CCHA1-RNAi vs EECG>CCHa1-RNAi | 35 38 | | 0.8027 |
| 3 (f) Number of sleep bouts | EECG>GFP-RNAi vs EECG>CCHa1-RNAi | 33 38 | One-way ANOVA F (2,103) = 0.693 Tukey's multiple comparisons test | 0.7667 |
| | UAS-CCHA1-RNAi vs EECG>CCHa1-RNAi | 35 38 | | 0.8692 |
| 3 (f) Length of sleep bouts | EECG>GFP-RNAi vs EECG>CCHa1-RNAi | 33 38 | One-way ANOVA F (2,103) = 0.564 Tukey's multiple comparisons test | 0.8142 |
| | UAS-CCHA1-RNAi vs EECG>CCHa1-RNAi | 35 38 | | 0.8822 |
| 3 (g) Arousability during wake | Pros-Gal4:Tub-Gal80$^{ts}$ >GFP-RNAi vs Pros-Gal4:Tub-Gal80$^{ts}$ >CCHa1-RNAi | 15 16 | One-way ANOVA F(2,45)=18.37 Tukey's multiple comparisons test | 0.0007 |
| | UAS-CCHA1-RNAi vs Pros-Gal4:Tub-Gal80$^{ts}$ >CCHa1-RNAi | 17 16 | | <0.0001 |
| 3 (g) Sleep duration | Pros-Gal4:Tub-Gal80$^{ts}$ >GFP-RNAi vs Pros-Gal4:Tub-Gal80$^{ts}$ >CCHa1-RNAi | 32 45 | One-way ANOVA F (2,113) = 1.404 | 0.2595 |
| | UAS-CCHA1-RNAi vs Pros-Gal4:Tub-Gal80$^{ts}$ >CCHa1-RNAi | 39 45 | | 0.9812 |

| | | | | |
|---|---|---|---|---|
| | | | Tukey's multiple comparisons test | |
| 3 (g) Basal locomotion | Pros-Gal4:Tub-Gal80$^{ts}$ >GFP-RNAi vs Pros-Gal4:Tub-Gal80$^{ts}$ >CCHa1-RNAi | 32 45 | One-way ANOVA F (2,113) = 0.4353 Tukey's multiple comparisons test | 0.7999 |
| | UAS-CCHA1-RNAi vs Pros-Gal4:Tub-Gal80$^{ts}$ >CCHa1-RNAi | 39 45 | | 0.6476 |
| 3 (g) Number of sleep bouts | Pros-Gal4:Tub-Gal80$^{ts}$ >GFP-RNAi vs Pros-Gal4:Tub-Gal80$^{ts}$ >CCHa1-RNAi | 32 45 | One-way ANOVA F (2,113) = 0.139 Tukey's multiple comparisons test | 0.8599 |
| | UAS-CCHA1-RNAi vs Pros-Gal4:Tub-Gal80$^{ts}$ >CCHa1-RNAi | 39 45 | | 0.9571 |
| 3 (g) Length of sleep bouts | Pros-Gal4:Tub-Gal80$^{ts}$ >GFP-RNAi vs Pros-Gal4:Tub-Gal80$^{ts}$ >CCHa1-RNAi | 32 45 | One-way ANOVA F (2,113) = 0.116 Tukey's multiple comparisons test | 0.9577 |
| | UAS-CCHA1-RNAi vs Pros-Gal4:Tub-Gal80$^{ts}$ >CCHa1-RNAi | 39 45 | | 0.8831 |
| 3 (h) | EECG>GFP-RNAi vs EECG>CCHa1-RNAi | 3 3 | One-way ANOVA F (2,6) = 0.152 Tukey's multiple comparisons test | 0.9418 |
| | UAS-CCHA1-RNAi vs EECG>CCHa1-RNAi | 3 3 | | 0.9752 |
| **Extended Data Figure 4** | | | | |
| 4 (a) Arousability during wake, low amplitude | EECG-Gal4 21°C vs EECG>shi$^{ts}$ 21°C vs | 10 10 | Two-way ANOVA F (2, 48) = 10.82 Tukey's multiple comparisons test | 0.9752 |
| | UAS- shi$^{ts}$ 21°C vs EECG>shi$^{ts}$ 21°C vs | 7 10 | | 0.9893 |
| | EECG-Gal4 27°C vs EECG>shi$^{ts}$ 27°C vs | 10 10 | | <0.0001 |
| | UAS- shi$^{ts}$ 27°C vs EECG>shi$^{ts}$ 27°C vs | 7 10 | | <0.0001 |
| 4 (a) Arousability during wake, high amplitude | EECG-Gal4 21°C vs EECG>shi$^{ts}$ 21°C vs | 10 10 | Two-way ANOVA F (2, 48) = 1.027 | 0.3981 |
| | UAS- shi$^{ts}$ 21°C vs EECG>shi$^{ts}$ 21°C vs | 7 10 | | 0.4140 |

| | | | | |
|---|---|---|---|---|
| | EECG-Gal4 27°C vs<br>EECG>shi$^{ts}$ 27°C vs | 10<br>10 | Tukey's multiple comparisons test | 0.2871 |
| | UAS- shi$^{ts}$ 27°C vs<br>EECG>shi$^{ts}$ 27°C vs | 7<br>10 | | 0.6460 |
| 4 (a) Sleep duration | EECG-Gal4 21°C vs<br>EECG>shi$^{ts}$ 21°C vs | 38<br>28 | Two-way ANOVA<br>$F(2,186) = 1.934$<br>Tukey's multiple comparisons test | 0.9955 |
| | UAS- shi$^{ts}$ 21°C vs<br>EECG>shi$^{ts}$ 21°C vs | 28<br>28 | | 0.0010 |
| | EECG-Gal4 27°C vs<br>EECG>shi$^{ts}$ 27°C vs | 38<br>28 | | 0.8145 |
| | UAS- shi$^{ts}$ 27°C vs<br>EECG>shi$^{ts}$ 27°C vs | 28<br>28 | | 0.3246 |
| 4 (a) Basal locomotion | EECG-Gal4 21°C vs<br>EECG>shi$^{ts}$ 21°C vs | 38<br>28 | Two-way ANOVA<br>$F(2,186) = 0.129$<br>Tukey's multiple comparisons test | 0.8903 |
| | UAS- shi$^{ts}$ 21°C vs<br>EECG>shi$^{ts}$ 21°C vs | 28<br>28 | | 0.7140 |
| | EECG-Gal4 27°C vs<br>EECG>shi$^{ts}$ 27°C vs | 38<br>28 | | 0.9997 |
| | UAS- shi$^{ts}$ 27°C vs<br>EECG>shi$^{ts}$ 27°C vs | 28<br>28 | | 0.4645 |
| 4 (a) Number of sleep bouts | EECG-Gal4 21°C vs<br>EECG>shi$^{ts}$ 21°C vs | 38<br>28 | One-way ANOVA<br>$F(2,186) = 0.198$<br>Tukey's multiple comparisons test | 0.7026 |
| | UAS- shi$^{ts}$ 21°C vs<br>EECG>shi$^{ts}$ 21°C vs | 28<br>28 | | 0.9998 |
| | EECG-Gal4 27°C vs<br>EECG>shi$^{ts}$ 27°C vs | 38<br>28 | | 0.2152 |
| | UAS- shi$^{ts}$ 27°C vs<br>EECG>shi$^{ts}$ 27°C vs | 28<br>28 | | 0.9987 |
| 4 (a) Length of sleep bouts | EECG-Gal4 21°C vs<br>EECG>shi$^{ts}$ 21°C vs | 38<br>28 | One-way ANOVA<br>$F(2,186) = 0.129$<br>Tukey's multiple comparisons test | 0.8903 |
| | UAS- shi$^{ts}$ 21°C vs<br>EECG>shi$^{ts}$ 21°C vs | 28<br>28 | | 0.7140 |
| | EECG-Gal4 27°C vs<br>EECG>shi$^{ts}$ 27°C vs | 38<br>28 | | 0.9997 |
| | UAS- shi$^{ts}$ 27°C vs<br>EECG>shi$^{ts}$ 27°C vs | 28<br>28 | | 0.4645 |
| 4 (b) Arousablity during wake, low amplitude | EECG-Gal4 25°C vs<br>EECG>TrpA1;GFP-RNAi 25°C | 10<br>10 | Two-way ANOVA<br>$F(6,110) = 7.310$<br>Tukey's multiple comparisons test | >0.9999 |
| | UAS-TrpA1;UAS-GFP-RNAi 25°C vs<br>EECG>TrpA1;GFP-RNAi 25°C | 10<br>10 | | >0.9999 |
| | EECG-Gal4 25°C vs<br>EECG>TrpA1;CCHa1-RNAi 25°C | 10<br>10 | | >0.9999 |
| | UAS-TrpA1;UAS-CCHa1-RNAi 25°C vs<br>EECG>TrpA1;CCHa1-RNAi 25°C | 10<br>10 | | >0.9999 |
| | EECG>TrpA1;GFP-RNAi 25°C vs<br>EECG>TrpA1;CCHa1-RNAi 25°C | 10<br>10 | | >0.9999 |
| | EECG-Gal4 25°C vs | 6 | | 0.0946 |

| | | | | |
|---|---|---|---|---|
| | EECG>CCHa1-RNAi;GFP 25°C | 6 | | |
| | UAS-CCHa1-RNAi;UAS-GFP 25°C vs<br>EECG>CCHa1-RNAi;GFP 25°C | 6<br>6 | | 0.9716 |
| | EECG>CCHa1-RNAi;GFP 25°C vs<br>EECG>TrpA1;CCHa1-RNAi 25°C | 6<br>10 | | 0.0066 |
| | EECG-Gal4 27°C vs<br>EECG>TrpA1;GFP-RNAi 27°C | 10<br>10 | | <0.0001 |
| | UAS-TrpA1;UAS-GFP-RNAi 27°C vs<br>EECG>TrpA1;GFP-RNAi 27°C | 10<br>10 | | 0.9022 |
| | EECG-Gal4 27°C vs<br>EECG>TrpA1;CCHa1-RNAi 27°C | 10<br>10 | | <0.0001 |
| | UAS-TrpA1;UAS-CCHa1-RNAi 27°C vs<br>EECG>TrpA1;CCHa1-RNAi 27°C | 10<br>10 | | <0.0001 |
| | EECG>TrpA1;GFP-RNAi 27°C vs<br>EECG>TrpA1;CCHa1-RNAi 27°C | 10<br>10 | | <0.0001 |
| | EECG-Gal4 27°C vs<br>EECG>CCHa1-RNAi;GFP 27°C | 6<br>6 | | 0.0003 |
| | UAS-CCHa1-RNAi;UAS-GFP 27°C vs<br>EECG>CCHa1-RNAi;GFP 27°C | 6<br>6 | | >0.9999 |
| | EECG>CCHa1-RNAi;GFP 27°C vs<br>EECG>TrpA1;CCHa1-RNAi 27°C | 6<br>10 | | >0.9999 |
| 4 (b)<br>Arousablity during wake, high amplitude | EECG-Gal4 25°C vs<br>EECG>TrpA1;GFP-RNAi 25°C | 10<br>10 | Two-way ANOVA<br>F (6,110) = 2.201<br>Tukey's multiple comparisons test | >0.9999 |
| | UAS-TrpA1;UAS-GFP-RNAi 25°C vs<br>EECG>TrpA1;GFP-RNAi 25°C | 10<br>10 | | 0.9999 |
| | EECG-Gal4 25°C vs<br>EECG>TrpA1;CCHa1-RNAi 25°C | 10<br>10 | | >0.9999 |
| | UAS-TrpA1;UAS-CCHa1-RNAi 25°C vs<br>EECG>TrpA1;CCHa1-RNAi 25°C | 10<br>10 | | 0.9884 |
| | EECG>TrpA1;GFP-RNAi 25°C vs<br>EECG>TrpA1;CCHa1-RNAi 25°C | 10<br>10 | | >0.9999 |
| | EECG-Gal4 25°C vs<br>EECG>CCHa1-RNAi;GFP 25°C | 6<br>6 | | >0.9999 |
| | UAS-CCHa1-RNAi;UAS-GFP 25°C vs<br>EECG>CCHa1-RNAi;GFP 25°C | 6<br>6 | | >0.9999 |
| | EECG>CCHa1-RNAi;GFP 25°C vs<br>EECG>TrpA1;CCHa1-RNAi 25°C | 6<br>10 | | >0.9999 |
| | EECG-Gal4 27°C vs<br>EECG>TrpA1;GFP-RNAi 27°C | 10<br>10 | | 0.2224 |
| | UAS-TrpA1;UAS-GFP-RNAi 27°C vs<br>EECG>TrpA1;GFP-RNAi 27°C | 10<br>10 | | 0.2224 |
| | EECG-Gal4 27°C vs<br>EECG>TrpA1;CCHa1-RNAi 27°C | 10<br>10 | | >0.9999 |
| | UAS-TrpA1;UAS-CCHa1-RNAi 27°C vs<br>EECG>TrpA1;CCHa1-RNAi 27°C | 10<br>10 | | >0.9999 |
| | EECG>TrpA1;GFP-RNAi 27°C vs<br>EECG>TrpA1;CCHa1-RNAi 27°C | 10<br>10 | | 0.0039 |

| | | | | |
|---|---|---|---|---|
| | EECG-Gal4 27°C vs<br>EECG>CCHa1-RNAi;GFP 27°C | 6<br>6 | | >0.9999 |
| | UAS-CCHa1-RNAi;UAS-GFP 27°C vs<br>EECG>CCHa1-RNAi;GFP 27°C | 6<br>6 | | >0.9999 |
| | EECG>CCHa1-RNAi;GFP 27°C vs<br>EECG>TrpA1;CCHa1-RNAi 27°C | 6<br>10 | | >0.9999 |
| 4 (b) Sleep duration | EECG-Gal4 25°C vs<br>EECG>TrpA1;GFP-RNAi 25°C | 75<br>61 | Two-way ANOVA F (6,840) = 6.079 Tukey's multiple comparisons test | <0.0001 |
| | UAS-TrpA1;UAS-GFP-RNAi 25°C vs<br>EECG>TrpA1;GFP-RNAi 25°C | 63<br>61 | | 0.0034 |
| | EECG-Gal4 25°C vs<br>EECG>TrpA1;CCHa1-RNAi 25°C | 75<br>62 | | 0.0004 |
| | UAS-TrpA1;UAS-CCHa1-RNAi 25°C vs<br>EECG>TrpA1;CCHa1-RNAi 25°C | 63<br>62 | | <0.0001 |
| | EECG>TrpA1;GFP-RNAi 25°C vs<br>EECG>TrpA1;CCHa1-RNAi 25°C | 61<br>62 | | >0.9999 |
| | EECG-Gal4 25°C vs<br>EECG>CCHa1-RNAi;GFP 25°C | 75<br>59 | | 0.5770 |
| | UAS-CCHa1-RNAi;UAS-GFP 25°C vs<br>EECG>CCHa1-RNAi;GFP 25°C | 57<br>59 | | <0.0001 |
| | EECG>CCHa1-RNAi;GFP 25°C vs<br>EECG>TrpA1;CCHa1-RNAi 25°C | 59<br>57 | | 0.9977 |
| | EECG-Gal4 27°C vs<br>EECG>TrpA1;GFP-RNAi 27°C | 75<br>61 | | <0.0001 |
| | UAS-TrpA1;UAS-GFP-RNAi 27°C vs<br>EECG>TrpA1;GFP-RNAi 27°C | 63<br>61 | | <0.0001 |
| | EECG-Gal4 27°C vs<br>EECG>TrpA1;CCHa1-RNAi 27°C | 75<br>62 | | <0.0001 |
| | UAS-TrpA1;UAS-CCHa1-RNAi 27°C vs<br>EECG>TrpA1;CCHa1-RNAi 27°C | 63<br>62 | | <0.0001 |
| | EECG>TrpA1;GFP-RNAi 27°C vs<br>EECG>TrpA1;CCHa1-RNAi 27°C | 61<br>62 | | >0.9999 |
| | EECG-Gal4 27°C vs<br>EECG>CCHa1-RNAi;GFP 27°C | 75<br>59 | | >0.9999 |
| | UAS-CCHa1-RNAi;UAS-GFP 27°C vs<br>EECG>CCHa1-RNAi;GFP 27°C | 57<br>59 | | <0.0001 |
| | EECG>CCHa1-RNAi;GFP 27°C vs<br>EECG>TrpA1;CCHa1-RNAi 27°C | 59<br>57 | | <0.0001 |
| 4 (b) Basal locomotion | EECG-Gal4 25°C vs<br>EECG>TrpA1;GFP-RNAi 25°C | 75<br>61 | Two-way ANOVA F (6,840) = 2.284 Tukey's multiple comparisons test | 0.6575 |
| | UAS-TrpA1;UAS-GFP-RNAi 25°C vs<br>EECG>TrpA1;GFP-RNAi 25°C | 63<br>61 | | 0.8041 |
| | EECG-Gal4 25°C vs<br>EECG>TrpA1;CCHa1-RNAi 25°C | 75<br>62 | | >0.9999 |
| | UAS-TrpA1;UAS-CCHa1-RNAi 25°C vs<br>EECG>TrpA1;CCHa1-RNAi 25°C | 63<br>62 | | <0.0001 |
| | EECG>TrpA1;GFP-RNAi 25°C vs | 61 | | 0.6368 |

| | | | | |
|---|---|---|---|---|
| | EECG>TrpA1;CCHa1-RNAi 25°C | 62 | | |
| | EECG-Gal4 25°C vs<br>EECG>CCHa1-RNAi;GFP 25°C | 75<br>59 | | 0.3221 |
| | UAS-CCHa1-RNAi;UAS-GFP 25°C vs<br>EECG>CCHa1-RNAi;GFP 25°C | 57<br>59 | | >0.9999 |
| | EECG>CCHa1-RNAi;GFP 25°C vs<br>EECG>TrpA1;CCHa1-RNAi 25°C | 59<br>57 | | >0.9999 |
| | EECG-Gal4 27°C vs<br>EECG>TrpA1;GFP-RNAi 27°C | 75<br>61 | | >0.9999 |
| | UAS-TrpA1;UAS-GFP-RNAi 27°C vs<br>EECG>TrpA1;GFP-RNAi 27°C | 63<br>61 | | 0.9511 |
| | EECG-Gal4 27°C vs<br>EECG>TrpA1;CCHa1-RNAi 27°C | 75<br>62 | | >0.9999 |
| | UAS-TrpA1;UAS-CCHa1-RNAi 27°C vs<br>EECG>TrpA1;CCHa1-RNAi 27°C | 63<br>62 | | <0.0001 |
| | EECG>TrpA1;GFP-RNAi 27°C vs<br>EECG>TrpA1;CCHa1-RNAi 27°C | 61<br>62 | | >0.9999 |
| | EECG-Gal4 27°C vs<br>EECG>CCHa1-RNAi;GFP 27°C | 75<br>59 | | 0.9554 |
| | UAS-CCHa1-RNAi;UAS-GFP 27°C vs<br>EECG>CCHa1-RNAi;GFP 27°C | 57<br>59 | | 0.8578 |
| | EECG>CCHa1-RNAi;GFP 27°C vs<br>EECG>TrpA1;CCHa1-RNAi 27°C | 59<br>57 | | 0.9999 |
| 4 (b) Number of sleep bouts | EECG-Gal4 25°C vs<br>EECG>TrpA1;GFP-RNAi 25°C | 75<br>61 | Two-way ANOVA $F(6,840) = 3.262$ Tukey's multiple comparisons test | >0.9999 |
| | UAS-TrpA1;UAS-GFP-RNAi 25°C vs<br>EECG>TrpA1;GFP-RNAi 25°C | 63<br>61 | | >0.9999 |
| | EECG-Gal4 25°C vs<br>EECG>TrpA1;CCHa1-RNAi 25°C | 75<br>62 | | 0.8472 |
| | UAS-TrpA1;UAS-CCHa1-RNAi 25°C vs<br>EECG>TrpA1;CCHa1-RNAi 25°C | 63<br>62 | | 0.0968 |
| | EECG>TrpA1;GFP-RNAi 25°C vs<br>EECG>TrpA1;CCHa1-RNAi 25°C | 61<br>62 | | 0.5172 |
| | EECG-Gal4 25°C vs<br>EECG>CCHa1-RNAi;GFP 25°C | 75<br>59 | | 0.1192 |
| | UAS-CCHa1-RNAi;UAS-GFP 25°C vs<br>EECG>CCHa1-RNAi;GFP 25°C | 57<br>59 | | 0.7293 |
| | EECG>CCHa1-RNAi;GFP 25°C vs<br>EECG>TrpA1;CCHa1-RNAi 25°C | 59<br>57 | | >0.9999 |
| | EECG-Gal4 27°C vs<br>EECG>TrpA1;GFP-RNAi 27°C | 75<br>61 | | 0.1431 |
| | UAS-TrpA1;UAS-GFP-RNAi 27°C vs<br>EECG>TrpA1;GFP-RNAi 27°C | 63<br>61 | | 0.0227 |
| | EECG-Gal4 27°C vs<br>EECG>TrpA1;CCHa1-RNAi 27°C | 75<br>62 | | 0.0002 |
| | UAS-TrpA1;UAS-CCHa1-RNAi 27°C vs<br>EECG>TrpA1;CCHa1-RNAi 27°C | 63<br>62 | | >0.9999 |

| | | | | |
|---|---|---|---|---|
| | EECG>TrpA1;GFP-RNAi 27°C vs EECG>TrpA1;CCHa1-RNAi 27°C | 61 62 | | >0.9999 |
| | EECG-Gal4 27°C vs EECG>CCHa1-RNAi;GFP 27°C | 75 59 | | 0.0001 |
| | UAS-CCHa1-RNAi;UAS-GFP 27°C vs EECG>CCHa1-RNAi;GFP 27°C | 57 59 | | 0.0651 |
| | EECG>CCHa1-RNAi;GFP 27°C vs EECG>TrpA1;CCHa1-RNAi 27°C | 59 57 | | >0.9999 |
| 4 (b) Length of sleep bouts | EECG-Gal4 25°C vs EECG>TrpA1;GFP-RNAi 25°C | 75 61 | Two-way ANOVA F (6,840) = 1.776 Tukey's multiple comparisons test | 0.8764 |
| | UAS-TrpA1;UAS-GFP-RNAi 25°C vs EECG>TrpA1;GFP-RNAi 25°C | 63 61 | | >0.9999 |
| | EECG-Gal4 25°C vs EECG>TrpA1;CCHa1-RNAi 25°C | 75 62 | | >0.9999 |
| | UAS-TrpA1;UAS-CCHa1-RNAi 25°C vs EECG>TrpA1;CCHa1-RNAi 25°C | 63 62 | | <0.0001 |
| | EECG>TrpA1;GFP-RNAi 25°C vs EECG>TrpA1;CCHa1-RNAi 25°C | 61 62 | | 0.9991 |
| | EECG-Gal4 25°C vs EECG>CCHa1-RNAi;GFP 25°C | 75 59 | | >0.9999 |
| | UAS-CCHa1-RNAi;UAS-GFP 25°C vs EECG>CCHa1-RNAi;GFP 25°C | 57 59 | | <0.0001 |
| | EECG>CCHa1-RNAi;GFP 25°C vs EECG>TrpA1;CCHa1-RNAi 25°C | 59 57 | | >0.9999 |
| | EECG-Gal4 27°C vs EECG>TrpA1;GFP-RNAi 27°C | 75 61 | | 0.7216 |
| | UAS-TrpA1;UAS-GFP-RNAi 27°C vs EECG>TrpA1;GFP-RNAi 27°C | 63 61 | | >0.9999 |
| | EECG-Gal4 27°C vs EECG>TrpA1;CCHa1-RNAi 27°C | 75 62 | | >0.9999 |
| | UAS-TrpA1;UAS-CCHa1-RNAi 27°C vs EECG>TrpA1;CCHa1-RNAi 27°C | 63 62 | | 0.0016 |
| | EECG>TrpA1;GFP-RNAi 27°C vs EECG>TrpA1;CCHa1-RNAi 27°C | 61 62 | | 0.8115 |
| | EECG-Gal4 27°C vs EECG>CCHa1-RNAi;GFP 27°C | 75 59 | | 0.9683 |
| | UAS-CCHa1-RNAi;UAS-GFP 27°C vs EECG>CCHa1-RNAi;GFP 27°C | 57 59 | | <0.0001 |
| | EECG>CCHa1-RNAi;GFP 27°C vs EECG>TrpA1;CCHa1-RNAi 27°C | 59 57 | | 0.9870 |
| **Extended Data Figure 5** | | | | |
| 5 (a) Pros-Gal4:Tub-Gal80$^{ts}$ Low amplitude | Pros-Gal4:Tub-Gal80$^{ts}$ -Gal4 25°C vs Pros-Gal4:Tub-Gal80$^{ts}$ >TrpA1;GFP-RNAi 25°C | 6 6 | Two-way ANOVA F (6,80) = 2.680 Tukey's multiple | 0.2159 |
| | UAS-TrpA1;UAS-GFP-RNAi 25°C vs Pros-Gal4:Tub-Gal80$^{ts}$ >TrpA1;GFP-RNAi 25°C | 5 6 | | 0.9991 |

| | | | | |
|---|---|---|---|---|
| | Pros-Gal4:Tub-Gal80$^{ts}$ -Gal4 25°C vs Pros-Gal4:Tub-Gal80$^{ts}$ >TrpA1;CCHa1-RNAi 25°C | 6<br>6 | comparisons test | 0.6388 |
| | UAS-TrpA1;UAS-CCHa1-RNAi 25°C vs Pros-Gal4:Tub-Gal80$^{ts}$ >TrpA1;CCHa1-RNAi 25°C | 5<br>6 | | 0.2540 |
| | Pros-Gal4:Tub-Gal80$^{ts}$ >TrpA1;GFP-RNAi 25°C vs Pros-Gal4:Tub-Gal80$^{ts}$ >TrpA1;CCHa1-RNAi 25°C | 6<br>6 | | 0.9908 |
| | Pros-Gal4:Tub-Gal80$^{ts}$ -Gal4 25°C vs Pros-Gal4:Tub-Gal80$^{ts}$ >CCHa1-RNAi;GFP 25°C | 6<br>6 | | 0.4143 |
| | UAS-CCHa1-RNAi;UAS-GFP 25°C vs Pros-Gal4:Tub-Gal80$^{ts}$ >CCHa1-RNAi;GFP 25°C | 6<br>6 | | 0.7900 |
| | Pros-Gal4:Tub-Gal80$^{ts}$ >CCHa1-RNAi;GFP 25°C vs Pros-Gal4:Tub-Gal80$^{ts}$ >TrpA1;CCHa1-RNAi 25°C | 6<br>6 | | 0.0074 |
| | Pros-Gal4:Tub-Gal80$^{ts}$ -Gal4 27°C vs Pros-Gal4:Tub-Gal80$^{ts}$ >TrpA1;GFP-RNAi 27°C | 8<br>9 | | 0.9976 |
| | UAS-TrpA1;UAS-GFP-RNAi 27°C vs Pros-Gal4:Tub-Gal80$^{ts}$ >TrpA1;GFP-RNAi 27°C | 8<br>9 | | >0.9999 |
| | Pros-Gal4:Tub-Gal80$^{ts}$ -Gal4 27°C vs Pros-Gal4:Tub-Gal80$^{ts}$ >TrpA1;CCHa1-RNAi 27°C | 8<br>8 | | 0.9771 |
| | UAS-TrpA1;UAS-CCHa1-RNAi 27°C vs Pros-Gal4:Tub-Gal80$^{ts}$ >TrpA1;CCHa1-RNAi 27°C | 6<br>8 | | >0.9999 |
| | Pros-Gal4:Tub-Gal80$^{ts}$ >TrpA1;GFP-RNAi 27°C vs Pros-Gal4:Tub-Gal80$^{ts}$ >TrpA1;CCHa1-RNAi 27°C | 9<br>8 | | >0.9999 |
| | Pros-Gal4:Tub-Gal80$^{ts}$ -Gal4 27°C vs Pros-Gal4:Tub-Gal80$^{ts}$ >CCHa1-RNAi;GFP 27°C | 8<br>10 | | 0.3341 |
| | UAS-CCHa1-RNAi;UAS-GFP 27°C vs Pros-Gal4:Tub-Gal80$^{ts}$ >CCHa1-RNAi;GFP 27°C | 6<br>10 | | 0.0006 |
| | Pros-Gal4:Tub-Gal80$^{ts}$ >CCHa1-RNAi;GFP 27°C vs Pros-Gal4:Tub-Gal80$^{ts}$ >TrpA1;CCHa1-RNAi 27°C | 10<br>8 | | 0.0375 |
| 5 (a) Pros-Gal4:Tub- | Pros-Gal4:Tub-Gal80$^{ts}$ -Gal4 25°C vs | 6<br>6 | Two-way ANOVA | 0.4617 |

| | | | | |
|---|---|---|---|---|
| Gal80$^{ts}$ Low amplitude | Pros-Gal4:Tub-Gal80$^{ts}$ >TrpA1;GFP-RNAi 25°C | | F (6,80) = 9.468 Tukey's multiple comparisons test | |
| | UAS-TrpA1;UAS-GFP-RNAi 25°C vs Pros-Gal4:Tub-Gal80$^{ts}$ >TrpA1;GFP-RNAi 25°C | 5 6 | | 0.9973 |
| | Pros-Gal4:Tub-Gal80$^{ts}$ -Gal4 25°C vs Pros-Gal4:Tub-Gal80$^{ts}$ >TrpA1;CCHa1-RNAi 25°C | 6 6 | | 0.8241 |
| | UAS-TrpA1;UAS-CCHa1-RNAi 25°C vs Pros-Gal4:Tub-Gal80$^{ts}$ >TrpA1;CCHa1-RNAi 25°C | 5 6 | | 0.8829 |
| | Pros-Gal4:Tub-Gal80$^{ts}$ >TrpA1;GFP-RNAi 25°C vs Pros-Gal4:Tub-Gal80$^{ts}$ >TrpA1;CCHa1-RNAi 25°C | 6 6 | | 0.9946 |
| | Pros-Gal4:Tub-Gal80$^{ts}$ -Gal4 25°C vs Pros-Gal4:Tub-Gal80$^{ts}$ >CCHa1-RNAi;GFP 25°C | 6 6 | | 0.5560 |
| | UAS-CCHa1-RNAi;UAS-GFP 25°C vs Pros-Gal4:Tub-Gal80$^{ts}$ >CCHa1-RNAi;GFP 25°C | 6 6 | | 0.9168 |
| | Pros-Gal4:Tub-Gal80$^{ts}$ >CCHa1-RNAi;GFP 25°C vs Pros-Gal4:Tub-Gal80$^{ts}$ >TrpA1;CCHa1-RNAi 25°C | 6 6 | | 0.9989 |
| | Pros-Gal4:Tub-Gal80$^{ts}$ -Gal4 27°C vs Pros-Gal4:Tub-Gal80$^{ts}$ >TrpA1;GFP-RNAi 27°C | 8 9 | | <0.0001 |
| | UAS-TrpA1;UAS-GFP-RNAi 27°C vs Pros-Gal4:Tub-Gal80$^{ts}$ >TrpA1;GFP-RNAi 27°C | 8 9 | | <0.0001 |
| | Pros-Gal4:Tub-Gal80$^{ts}$ -Gal4 27°C vs Pros-Gal4:Tub-Gal80$^{ts}$ >TrpA1;CCHa1-RNAi 27°C | 8 8 | | <0.0001 |
| | UAS-TrpA1;UAS-CCHa1-RNAi 27°C vs Pros-Gal4:Tub-Gal80$^{ts}$ >TrpA1;CCHa1-RNAi 27°C | 6 8 | | <0.0001 |
| | Pros-Gal4:Tub-Gal80$^{ts}$ >TrpA1;GFP-RNAi 27°C vs Pros-Gal4:Tub-Gal80$^{ts}$ >TrpA1;CCHa1-RNAi 27°C | 9 8 | | 0.0001 |
| | Pros-Gal4:Tub-Gal80$^{ts}$ -Gal4 27°C vs Pros-Gal4:Tub-Gal80$^{ts}$ >CCHa1-RNAi;GFP 27°C | 8 10 | | >0.9999 |
| | UAS-CCHa1-RNAi;UAS-GFP 27°C vs Pros-Gal4:Tub-Gal80$^{ts}$ >CCHa1-RNAi;GFP 27°C | 6 10 | | 0.5664 |

| | | | | |
|---|---|---|---|---|
| | Pros-Gal4:Tub-Gal80$^{ts}$ >CCHa1-RNAi;GFP 27°C vs Pros-Gal4:Tub-Gal80$^{ts}$ >TrpA1;CCHa1-RNAi 27°C | 10 8 | | <0.0001 |
| 5 (a) Sleep duration | Pros-Gal4:Tub-Gal80$^{ts}$ -Gal4 25°C vs Pros-Gal4:Tub-Gal80$^{ts}$ >TrpA1;GFP-RNAi 25°C | 36 25 | Two-way ANOVA F (6,434) = 18.11 Tukey's multiple comparisons test | <0.0001 |
| | UAS-TrpA1;UAS-GFP-RNAi 25°C vs Pros-Gal4:Tub-Gal80$^{ts}$ >TrpA1;GFP-RNAi 25°C | 32 25 | | <0.0001 |
| | Pros-Gal4:Tub-Gal80$^{ts}$ -Gal4 25°C vs Pros-Gal4:Tub-Gal80$^{ts}$ >TrpA1;CCHa1-RNAi 25°C | 36 38 | | 0.0656 |
| | UAS-TrpA1;UAS-CCHa1-RNAi 25°C vs Pros-Gal4:Tub-Gal80$^{ts}$ >TrpA1;CCHa1-RNAi 25°C | 36 38 | | 0.5842 |
| | Pros-Gal4:Tub-Gal80$^{ts}$ >TrpA1;GFP-RNAi 25°C vs Pros-Gal4:Tub-Gal80$^{ts}$ >TrpA1;CCHa1-RNAi 25°C | 25 38 | | <0.0001 |
| | Pros-Gal4:Tub-Gal80$^{ts}$ -Gal4 25°C vs Pros-Gal4:Tub-Gal80$^{ts}$ >CCHa1-RNAi;GFP 25°C | 36 25 | | >0.9999 |
| | UAS-CCHa1-RNAi;UAS-GFP 25°C vs Pros-Gal4:Tub-Gal80$^{ts}$ >CCHa1-RNAi;GFP 25°C | 32 25 | | 0.9770 |
| | Pros-Gal4:Tub-Gal80$^{ts}$ >CCHa1-RNAi;GFP 25°C vs Pros-Gal4:Tub-Gal80$^{ts}$ >TrpA1;CCHa1-RNAi 25°C | 38 25 | | 0.2298 |
| | Pros-Gal4:Tub-Gal80$^{ts}$ -Gal4 27°C vs Pros-Gal4:Tub-Gal80$^{ts}$ >TrpA1;GFP-RNAi 27°C | 36 25 | | <0.0001 |
| | UAS-TrpA1;UAS-GFP-RNAi 27°C vs Pros-Gal4:Tub-Gal80$^{ts}$ >TrpA1;GFP-RNAi 27°C | 32 25 | | <0.0001 |
| | Pros-Gal4:Tub-Gal80$^{ts}$ -Gal4 27°C vs Pros-Gal4:Tub-Gal80$^{ts}$ >TrpA1;CCHa1-RNAi 27°C | 36 38 | | <0.0001 |
| | UAS-TrpA1;UAS-CCHa1-RNAi 27°C vs Pros-Gal4:Tub-Gal80$^{ts}$ >TrpA1;CCHa1-RNAi 27°C | 36 38 | | <0.0001 |
| | Pros-Gal4:Tub-Gal80$^{ts}$ >TrpA1;GFP-RNAi 27°C vs Pros-Gal4:Tub-Gal80$^{ts}$ >TrpA1;CCHa1-RNAi 27°C | 25 38 | | 0.0295 |
| | Pros-Gal4:Tub-Gal80$^{ts}$ -Gal4 27°C vs | 36 25 | | 0.9862 |

| | | | | |
|---|---|---|---|---|
| | Pros-Gal4:Tub-Gal80ts >CCHa1-RNAi;GFP 27°C | | | |
| | UAS-CCHa1-RNAi;UAS-GFP 27°C vs Pros-Gal4:Tub-Gal80ts >CCHa1-RNAi;GFP 27°C | 32 25 | | 0.5532 |
| | Pros-Gal4:Tub-Gal80ts >CCHa1-RNAi;GFP 27°C vs Pros-Gal4:Tub-Gal80ts >TrpA1;CCHa1-RNAi 27°C | 38 25 | | <0.0001 |
| 5 (a) Basal locomotion | Pros-Gal4:Tub-Gal80ts -Gal4 25°C vs Pros-Gal4:Tub-Gal80ts >TrpA1;GFP-RNAi 25°C | 36 25 | Two-way ANOVA F (6,434) = 3.823 Tukey's multiple comparisons test | 0.0197 |
| | UAS-TrpA1;UAS-GFP-RNAi 25°C vs Pros-Gal4:Tub-Gal80ts >TrpA1;GFP-RNAi 25°C | 32 25 | | 0.9819 |
| | Pros-Gal4:Tub-Gal80ts -Gal4 25°C vs Pros-Gal4:Tub-Gal80ts >TrpA1;CCHa1-RNAi 25°C | 36 38 | | 0.2759 |
| | UAS-TrpA1;UAS-CCHa1-RNAi 25°C vs Pros-Gal4:Tub-Gal80ts >TrpA1;CCHa1-RNAi 25°C | 36 38 | | 0.6124 |
| | Pros-Gal4:Tub-Gal80ts >TrpA1;GFP-RNAi 25°C vs Pros-Gal4:Tub-Gal80ts >TrpA1;CCHa1-RNAi 25°C | 25 38 | | 0.8307 |
| | Pros-Gal4:Tub-Gal80ts -Gal4 25°C vs Pros-Gal4:Tub-Gal80ts >CCHa1-RNAi;GFP 25°C | 36 25 | | 0.9999 |
| | UAS-CCHa1-RNAi;UAS-GFP 25°C vs Pros-Gal4:Tub-Gal80ts >CCHa1-RNAi;GFP 25°C | 32 25 | | 0.9985 |
| | Pros-Gal4:Tub-Gal80ts >CCHa1-RNAi;GFP 25°C vs Pros-Gal4:Tub-Gal80ts >TrpA1;CCHa1-RNAi 25°C | 38 25 | | 0.5142 |
| | Pros-Gal4:Tub-Gal80ts -Gal4 27°C vs Pros-Gal4:Tub-Gal80ts >TrpA1;GFP-RNAi 27°C | 36 25 | | <0.0001 |
| | UAS-TrpA1;UAS-GFP-RNAi 27°C vs Pros-Gal4:Tub-Gal80ts >TrpA1;GFP-RNAi 27°C | 32 25 | | <0.0001 |
| | Pros-Gal4:Tub-Gal80ts -Gal4 27°C vs Pros-Gal4:Tub-Gal80ts >TrpA1;CCHa1-RNAi 27°C | 36 38 | | <0.0001 |
| | UAS-TrpA1;UAS-CCHa1-RNAi 27°C vs Pros-Gal4:Tub-Gal80ts >TrpA1;CCHa1-RNAi 27°C | 36 38 | | <0.0001 |

| | | | | |
|---|---|---|---|---|
| | Pros-Gal4:Tub-Gal80$^{ts}$ >TrpA1;GFP-RNAi 27°C vs Pros-Gal4:Tub-Gal80$^{ts}$ >TrpA1;CCHa1-RNAi 27°C | 25<br>38 | | 0.5114 |
| | Pros-Gal4:Tub-Gal80$^{ts}$ -Gal4 27°C vs Pros-Gal4:Tub-Gal80$^{ts}$ >CCHa1-RNAi;GFP 27°C | 36<br>25 | | 0.9960 |
| | UAS-CCHa1-RNAi;UAS-GFP 27°C vs Pros-Gal4:Tub-Gal80$^{ts}$ >CCHa1-RNAi;GFP 27°C | 32<br>25 | | 0.9700 |
| | Pros-Gal4:Tub-Gal80$^{ts}$ >CCHa1-RNAi;GFP 27°C vs Pros-Gal4:Tub-Gal80$^{ts}$ >TrpA1;CCHa1-RNAi 27°C | 38<br>25 | | <0.0001 |
| 5 (a) Number of sleep bouts | Pros-Gal4:Tub-Gal80$^{ts}$ -Gal4 25°C vs Pros-Gal4:Tub-Gal80$^{ts}$ >TrpA1;GFP-RNAi 25°C | 36<br>25 | Two-way ANOVA $F (6,434) = 22.56$ Tukey's multiple comparisons test | <0.0001 |
| | UAS-TrpA1;UAS-GFP-RNAi 25°C vs Pros-Gal4:Tub-Gal80$^{ts}$ >TrpA1;GFP-RNAi 25°C | 32<br>25 | | 0.5035 |
| | Pros-Gal4:Tub-Gal80$^{ts}$ -Gal4 25°C vs Pros-Gal4:Tub-Gal80$^{ts}$ >TrpA1;CCHa1-RNAi 25°C | 36<br>38 | | <0.0001 |
| | UAS-TrpA1;UAS-CCHa1-RNAi 25°C vs Pros-Gal4:Tub-Gal80$^{ts}$ >TrpA1;CCHa1-RNAi 25°C | 36<br>38 | | 0.0015 |
| | Pros-Gal4:Tub-Gal80$^{ts}$ >TrpA1;GFP-RNAi 25°C vs Pros-Gal4:Tub-Gal80$^{ts}$ >TrpA1;CCHa1-RNAi 25°C | 25<br>38 | | 0.9999 |
| | Pros-Gal4:Tub-Gal80$^{ts}$ -Gal4 25°C vs Pros-Gal4:Tub-Gal80$^{ts}$ >CCHa1-RNAi;GFP 25°C | 36<br>25 | | 0.3283 |
| | UAS-CCHa1-RNAi;UAS-GFP 25°C vs Pros-Gal4:Tub-Gal80$^{ts}$ >CCHa1-RNAi;GFP 25°C | 32<br>25 | | 0.0029 |
| | Pros-Gal4:Tub-Gal80$^{ts}$ >CCHa1-RNAi;GFP 25°C vs Pros-Gal4:Tub-Gal80$^{ts}$ >TrpA1;CCHa1-RNAi 25°C | 38<br>25 | | <0.0001 |
| | Pros-Gal4:Tub-Gal80$^{ts}$ -Gal4 27°C vs Pros-Gal4:Tub-Gal80$^{ts}$ >TrpA1;GFP-RNAi 27°C | 36<br>25 | | <0.0001 |
| | UAS-TrpA1;UAS-GFP-RNAi 27°C vs Pros-Gal4:Tub-Gal80$^{ts}$ >TrpA1;GFP-RNAi 27°C | 32<br>25 | | <0.0001 |
| | Pros-Gal4:Tub-Gal80$^{ts}$ -Gal4 27°C vs | 36<br>38 | | 0.8512 |

| | | | | |
|---|---|---|---|---|
| | Pros-Gal4:Tub-Gal80$^{ts}$ >TrpA1;CCHa1-RNAi 27°C | | | |
| | UAS-TrpA1;UAS-CCHa1-RNAi 27°C vs Pros-Gal4:Tub-Gal80$^{ts}$ >TrpA1;CCHa1-RNAi 27°C | 36<br>38 | | 0.4048 |
| | Pros-Gal4:Tub-Gal80$^{ts}$ >TrpA1;GFP-RNAi 27°C vs Pros-Gal4:Tub-Gal80$^{ts}$ >TrpA1;CCHa1-RNAi 27°C | 25<br>38 | | <0.0001 |
| | Pros-Gal4:Tub-Gal80$^{ts}$ -Gal4 27°C vs Pros-Gal4:Tub-Gal80$^{ts}$ >CCHa1-RNAi;GFP 27°C | 36<br>25 | | 0.9981 |
| | UAS-CCHa1-RNAi;UAS-GFP 27°C vs Pros-Gal4:Tub-Gal80$^{ts}$ >CCHa1-RNAi;GFP 27°C | 32<br>25 | | 0.7089 |
| | Pros-Gal4:Tub-Gal80$^{ts}$ >CCHa1-RNAi;GFP 27°C vs Pros-Gal4:Tub-Gal80$^{ts}$ >TrpA1;CCHa1-RNAi 27°C | 38<br>25 | | 0.9953 |
| 5 (a) Length of sleep bouts | Pros-Gal4:Tub-Gal80$^{ts}$ -Gal4 25°C vs Pros-Gal4:Tub-Gal80$^{ts}$ >TrpA1;GFP-RNAi 25°C | 36<br>25 | Two-way ANOVA $F (6,434) = 2.823$ Tukey's multiple comparisons test | <0.0001 |
| | UAS-TrpA1;UAS-GFP-RNAi 25°C vs Pros-Gal4:Tub-Gal80$^{ts}$ >TrpA1;GFP-RNAi 25°C | 32<br>25 | | 0.0029 |
| | Pros-Gal4:Tub-Gal80$^{ts}$ -Gal4 25°C vs Pros-Gal4:Tub-Gal80$^{ts}$ >TrpA1;CCHa1-RNAi 25°C | 36<br>38 | | 0.0006 |
| | UAS-TrpA1;UAS-CCHa1-RNAi 25°C vs Pros-Gal4:Tub-Gal80$^{ts}$ >TrpA1;CCHa1-RNAi 25°C | 36<br>38 | | 0.0126 |
| | Pros-Gal4:Tub-Gal80$^{ts}$ >TrpA1;GFP-RNAi 25°C vs Pros-Gal4:Tub-Gal80$^{ts}$ >TrpA1;CCHa1-RNAi 25°C | 25<br>38 | | 0.2526 |
| | Pros-Gal4:Tub-Gal80$^{ts}$ -Gal4 25°C vs Pros-Gal4:Tub-Gal80$^{ts}$ >CCHa1-RNAi;GFP 25°C | 36<br>25 | | 0.1065 |
| | UAS-CCHa1-RNAi;UAS-GFP 25°C vs Pros-Gal4:Tub-Gal80$^{ts}$ >CCHa1-RNAi;GFP 25°C | 32<br>25 | | 0.0017 |
| | Pros-Gal4:Tub-Gal80$^{ts}$ >CCHa1-RNAi;GFP 25°C vs Pros-Gal4:Tub-Gal80$^{ts}$ >TrpA1;CCHa1-RNAi 25°C | 38<br>25 | | <0.0001 |
| | Pros-Gal4:Tub-Gal80$^{ts}$ -Gal4 27°C vs Pros-Gal4:Tub-Gal80$^{ts}$ >TrpA1;GFP-RNAi 27°C | 36<br>25 | | <0.0001 |

| | | | | |
|---|---|---|---|---|
| | UAS-TrpA1;UAS-GFP-RNAi 27°C vs Pros-Gal4:Tub-Gal80$^{ts}$ >TrpA1;GFP-RNAi 27°C | 32<br>25 | | 0.0002 |
| | Pros-Gal4:Tub-Gal80$^{ts}$ -Gal4 27°C vs Pros-Gal4:Tub-Gal80$^{ts}$ >TrpA1;CCHa1-RNAi 27°C | 36<br>38 | | <0.0001 |
| | UAS-TrpA1;UAS-CCHa1-RNAi 27°C vs Pros-Gal4:Tub-Gal80$^{ts}$ >TrpA1;CCHa1-RNAi 27°C | 36<br>38 | | <0.0001 |
| | Pros-Gal4:Tub-Gal80$^{ts}$ >TrpA1;GFP-RNAi 27°C vs Pros-Gal4:Tub-Gal80$^{ts}$ >TrpA1;CCHa1-RNAi 27°C | 25<br>38 | | >0.9999 |
| | Pros-Gal4:Tub-Gal80$^{ts}$ -Gal4 27°C vs Pros-Gal4:Tub-Gal80$^{ts}$ >CCHa1-RNAi;GFP 27°C | 36<br>25 | | 0.9824 |
| | UAS-CCHa1-RNAi;UAS-GFP 27°C vs Pros-Gal4:Tub-Gal80$^{ts}$ >CCHa1-RNAi;GFP 27°C | 32<br>25 | | >0.9999 |
| | Pros-Gal4:Tub-Gal80$^{ts}$ >CCHa1-RNAi;GFP 27°C vs Pros-Gal4:Tub-Gal80$^{ts}$ >TrpA1;CCHa1-RNAi 27°C | 38<br>25 | | <0.0001 |
| **Extended Data Figure 6** | | | | |
| 6 (a) Cell activity | Regular vs<br>Regular + galactose | 19<br>17 | One-way ANOVA F (4,78) = 1.177 Tukey's multiple comparisons test | 0.9709 |
| | Regular vs<br>Regular + fructose | 19<br>15 | | >0.9999 |
| | Regular vs<br>Regular + propionic acid | 19<br>19 | | 0.9982 |
| | Regular vs<br>Regular + hexanoic acid | 19<br>13 | | 0.2882 |
| 6 (a) CCHa1 levels | Regular vs<br>Regular + galactose | 19<br>17 | Kruskal-Wallis (5,84) = 9.578 Dunn's multiple comparisons test | 0.2970 |
| | Regular vs<br>Regular + fructose | 19<br>15 | | >0.9999 |
| | Regular vs<br>Regular + propionic acid | 19<br>19 | | >0.9999 |
| | Regular vs<br>Regular + hexanoic acid | 19<br>13 | | >0.9999 |
| 6 (b) | Regular 6h vs<br>Regular + peptone 6h | 27<br>27 | T=test<br>t=3.778 df=52 | 0.0004 |
| | Regular 24h vs<br>Regular + peptone 24h | 22<br>29 | T=test<br>t=6.552 df=49 | <0.0001 |
| 6 (c) | Regular vs<br>Regular + peptone | 32<br>21 | Kruskal-Wallis (28,407) statistic=162.7 | <0.0001 |
| | Regular vs<br>Regular + all aa | 32<br>13 | | 0.0002 |
| | Regular vs | 32 | | 0.0321 |

| | | | | |
|---|---|---|---|---|
| | Regular + essential aa | | Dunn's multiple comparisons test | |
| | Regular vs<br>Regular + non-essential aa | 32 | | 0.0068 |
| | Regular vs<br>Regular + aromatic | 32 | | >0.9999 |
| | Regular vs<br>Regular + negative | 32 | | >0.9999 |
| | Regular vs<br>Regular + non-polar | 32 | | >0.9999 |
| | Regular vs<br>Regular + polar | 32 | | >0.9999 |
| | Regular vs<br>Regular + positive | 32 | | >0.9999 |
| | Regular vs<br>Regular + alanine | 32<br>12 | | >0.9999 |
| | Regular vs<br>Regular + arginine | 32<br>12 | | >0.9999 |
| | Regular vs<br>Regular + aspartic acid | 32<br>11 | | >0.9999 |
| | Regular vs<br>Regular + cysteine | 32<br>11 | | >0.9999 |
| | Regular vs<br>Regular + glutamic acid | 32<br>16 | | 0.4084 |
| | Regular vs<br>Regular + glycine | 32<br>13 | | 0.0347 |
| | Regular vs<br>Regular + histidine | 32<br>12 | | >0.9999 |
| | Regular vs<br>Regular + isoleucine | 32<br>13 | | >0.9999 |
| | Regular vs<br>Regular + leucine | 32<br>11 | | 0.6197 |
| | Regular vs<br>Regular + lysine | 32<br>16 | | >0.9999 |
| | Regular vs<br>Regular + methionine | 32<br>13 | | >0.9999 |
| | Regular vs<br>Regular + phenylalanine | 32<br>13 | | 0.7263 |
| | Regular vs<br>Regular + proline | 32<br>16 | | >0.9999 |
| | Regular vs<br>Regular + serine | 32<br>11 | | >0.9999 |
| | Regular vs<br>Regular + threonine | 32<br>12 | | >0.9999 |
| | Regular vs<br>Regular + tryptophan | 32<br>14 | | >0.9999 |
| | Regular vs<br>Regular + tyrosine | 32<br>15 | | >0.9999 |
| | Regular vs<br>Regular + valine | 32<br>14 | | >0.9999 |

| | | | | |
|---|---|---|---|---|
| 6 (e) Arousability during wake | Regular vs<br>Regular + peptone | 8<br>8 | T=test<br>t=3.378 df=14 | 0.0045 |
| 6 (e) Sleep duration | Regular vs<br>Regular + peptone | 32<br>24 | T-test<br>t=2.66 df=14 | 0.0103 |
| 6 (e) Basal locomotion | Regular vs<br>Regular + peptone | 32<br>24 | T-test<br>t=2.14 df=14 | 0.0368 |
| 6 (e) Number of sleep bouts | Regular vs<br>Regular + peptone | 32<br>24 | T-test<br>t=0.86 df=54 | 0.3955 |
| 6 (e) Length of sleep bouts | Regular vs<br>Regular + peptone | 32<br>24 | T-test<br>t=0.24 df=54 | 0.8094 |
| 6 (f) | Regular vs<br>Regular + peptone | 3<br>3 | T-test<br>t=0.517 df=4 | 0.6324 |
| 6 (g) Arousability during wake | EECG-Gal4 regular vs<br>EECG-Gal4 peptone | 14<br>14 | Two-way ANOVA F (2,79) = 2.14 Tukey's multiple comparisons test | 0.0080 |
| | UAS-CCHa1-RNAi regular vs<br>UAS-CCHa1-RNAi peptone | 14<br>14 | | 0.0480 |
| | EECG>CCHa1-RNAi regular vs<br>EECG>CCHa1-RNAi peptone | 15<br>14 | | 0.9671 |
| 6 (g) Arousability during wake | Pros-Gal4:Tub-Gal80$^{ts}$>GFP-RNAi regular vs<br>Pros-Gal4:Tub-Gal80$^{ts}$>GFP-RNAi peptone | 13<br>13 | Two-way ANOVA F (2,79) = 0.8089 Tukey's multiple comparisons test | 0.0006 |
| | UAS-CCHa1-RNAi regular vs<br>UAS-CCHa1-RNAi peptone | 12<br>12 | | 0.0073 |
| | Pros-Gal4:Tub-Gal80$^{ts}$>CCHa1-RNAi regular vs<br>Pros-Gal4:Tub-Gal80$^{ts}$>CCHa1-RNAi peptone | 16<br>14 | | 0.0645 |
| **Extended Data Figure 7** | | | | |
| 7 (a) | elav>GFP-RNAi vs<br>elav>CCHa1R-RNAi | 5<br>6 | One-way ANOVA F 12,46) = 1.102 Tukey's multiple comparisons test | 0.0001 |
| | UAS-CCHA1R-RNAi vs<br>elav>CCHa1R-RNAi | 11<br>6 | | <0.0001 |
| | Cha>GFP-RNAi vs<br>Cha>CCHa1R-RNAi | 2<br>2 | | 0.9983 |
| | UAS-CCHA1R-RNAi vs<br>Cha>CCHa1R-RNAi | 11<br>2 | | >0.9999 |
| | Tdc2>GFP-RNAi vs<br>Tdc2>CCHa1R-RNAi | 3<br>3 | | >0.9999 |
| | UAS-CCHA1R-RNAi vs<br>Tdc2>CCHa1R-RNAi | 11<br>3 | | 0.9730 |
| | TH>GFP-RNAi vs<br>TH>CCHa1R-RNAi | 3<br>6 | | 0.8573 |
| | UAS-CCHA1R-RNAi vs<br>TH>CCHa1R-RNAi | 11<br>6 | | 0.5973 |
| | Tim>GFP-RNAi vs<br>Tim>CCHa1R-RNAi | 5<br>4 | | >0.9999 |

| | | | | |
|---|---|---|---|---|
| | UAS-CCHA1R-RNAi vs<br>Tim>CCHa1R-RNAi | 11<br>4 | | >0.9999 |
| | vGlut>GFP-RNAi vs<br>vGlut>CCHa1R-RNAi | 6<br>3 | | 0.9886 |
| | UAS-CCHA1R-RNAi vs<br>vGlut>CCHa1R-RNAi | 11<br>3 | | >0.9999 |
| 7 (b)<br>Arousability<br>during wake | UAS-CCHa1R-RNAi vs<br>Ddc>CCHa1R-RNAi | 11<br>11 | One-way<br>ANOVA<br>F (4,47) =<br>5.452<br>Tukey's<br>multiple<br>comparisons<br>test | 0.0331 |
| | Ddc>GFP-RNAi vs<br>Ddc>CCHa1R-RNAi | 8<br>11 | | 0.0070 |
| | UAS-CCHa1R-RNAi vs<br>PAM>CCHa1R-RNAi | 11<br>13 | | 0.2830 |
| | PAM>GFP-RNAi vs<br>PAM>CCHa1R-RNAi | 9<br>13 | | 0.0898 |
| 7 (b) Sleep<br>duration | UAS-CCHa1R-RNAi vs<br>Ddc>CCHa1R-RNAi | 16<br>33 | Kruskal-<br>Wallis<br>(5,158) =<br>14.97<br>Dunn's<br>multiple<br>comparisons<br>test | 0.1137 |
| | Ddc>GFP-RNAi vs<br>Ddc>CCHa1R-RNAi | 36<br>33 | | 0.3113 |
| | UAS-CCHa1R-RNAi vs<br>PAM>CCHa1R-RNAi | 16<br>50 | | 0.0225 |
| | PAM>GFP-RNAi vs<br>PAM>CCHa1R-RNAi | 20<br>50 | | >0.9999 |
| 7 (b) Basal<br>locomotion | UAS-CCHa1R-RNAi vs<br>Ddc>CCHa1R-RNAi | 16<br>33 | One-way<br>ANOVA<br>F (4,150) =<br>6.774<br>Tukey's<br>multiple<br>comparisons<br>test | 0.8683 |
| | Ddc>GFP-RNAi vs<br>Ddc>CCHa1R-RNAi | 36<br>33 | | 0.0337 |
| | UAS-CCHa1R-RNAi vs<br>PAM>CCHa1R-RNAi | 16<br>50 | | 0.9931 |
| | PAM>GFP-RNAi vs<br>PAM>CCHa1R-RNAi | 20<br>50 | | 0.9974 |
| 7 (e)<br>Arousability<br>during wake | $PAM^{MB441B}$>GFP-RNAi vs<br>$PAM^{MB441B}$>CCHa1R-RNAi | 12<br>14 | One-way<br>ANOVA<br>F (2,32) =<br>0.512<br>Tukey's<br>multiple<br>comparisons<br>test | 0.5781 |
| | UAS-CCHa1R-RNAi vs<br>$PAM^{MB441B}$>CCHa1R-RNAi | 10<br>14 | | 0.8690 |
| 7 (e) Sleep<br>duration | $PAM^{MB441B}$>GFP-RNAi vs<br>$PAM^{MB441B}$>CCHa1R-RNAi | 33<br>53 | Kruskal-<br>Wallis (3,157)<br>= 5.07<br>Dunn's<br>multiple<br>comparisons<br>test | >0.9999 |
| | UAS-CCHa1R-RNAi vs<br>$PAM^{MB441B}$>CCHa1R-RNAi | 74<br>53 | | 0.0759 |
| 7 (e) Basal<br>locomotion | $PAM^{MB441B}$>GFP-RNAi vs<br>$PAM^{MB441B}$>CCHa1R-RNAi | 33<br>53 | Kruskal-<br>Wallis<br>3,157=33.95<br>Tukey's<br>multiple | <0.0001 |
| | UAS-CCHa1R-RNAi vs<br>$PAM^{MB441B}$>CCHa1R-RNAi | 74<br>53 | | <0.0001 |

| | | | | |
|---|---|---|---|---|
| | | | comparisons test | |
| 7 (e) Number of sleep bouts | PAM$^{MB441B}$>GFP-RNAi vs<br>PAM$^{MB441B}$>CCHa1R-RNAi | 33<br>53 | Kruskal-Wallis (3,157) = 1.368 Dunn's multiple comparisons test | 0.7585 |
| | UAS-CCHa1R-RNAi vs<br>PAM$^{MB441B}$>CCHa1R-RNAi | 74<br>53 | | >0.9999 |
| 7 (e) Length of sleep bouts | PAM$^{MB441B}$>GFP-RNAi vs<br>PAM$^{MB441B}$>CCHa1R-RNAi | 33<br>53 | Kruskal-Wallis (3,157) = 3.644 Dunn's multiple comparisons test | 0.6125 |
| | UAS-CCHa1R-RNAi vs<br>PAM$^{MB441B}$>CCHa1R-RNAi | 74<br>53 | | 0.1902 |
| 7 (f) | PAM$^{MB441B}$>GFP-RNAi vs<br>PAM$^{MB441B}$>CCHa1R-RNAi | 3<br>3 | One-way ANOVA F (2,6) = 0.863 Tukey's multiple comparisons test | 0.9666 |
| | UAS-CCHa1R-RNAi vs<br>PAM$^{MB441B}$>CCHa1R-RNAi | 3<br>3 | | 0.6075 |
| 7 (g) | PAM$^{MB441B}$ >GFP-RNAi regular vs<br>PAM$^{MB441B}$ >GFP-RNAi peptone | 10<br>10 | Two-way ANOVA F (2,50) = 0.0499 Tukey's multiple comparisons test | 0.0348 |
| | UAS-CCHa1R-RNAi regular vs<br>UAS-CCHa1R-RNAi peptone | 10<br>10 | | 0.0225 |
| | PAM$^{MB441B}$ >CCHa1R-RNAi regular vs<br>PAM$^{MB441B}$ >CCHa1R-RNAi peptone | 10<br>10 | | 0.0384 |
| **Extended Data Figure 8** | | | | |
| 8 (a) Arousability during wake, low amplitude | PAM$^{MB441B}$-Gal4 21°C vs<br>PAM$^{MB441B}$>TrpA1 21°C | 10<br>10 | Two-way ANOVA F (4,78) = 3.051 Tukey's multiple comparisons test | 0.9953 |
| | UAS-TrpA1 21°C vs<br>PAM$^{MB441B}$>TrpA1 21°C | 8<br>10 | | 0.8403 |
| | PAM$^{MB441B}$-Gal4 21°C vs<br>PAM$^{MB441B}$>shi$^{ts}$ 21°C | 10<br>10 | | 0.9830 |
| | UAS- shi$^{ts}$ 21°C vs<br>PAM$^{MB441B}$>shi$^{ts}$ 21°C | 6<br>10 | | >0.9999 |
| | PAM$^{MB441B}$-Gal4 27°C vs<br>PAM$^{MB441}$B>TrpA1 27°C | 10<br>10 | | >0.9999 |
| | UAS-TrpA1 27°C vs<br>PAM$^{MB441B}$>TrpA1 27°C | 8<br>10 | | >0.9999 |
| | PAM$^{MB441B}$-Gal4 27°C vs<br>PAM$^{MB441}$B>shi$^{ts}$ 27°C | 10<br>10 | | 0.1500 |

| | | | | |
|---|---|---|---|---|
| | UAS- $shi^{ts}$ 27°C vs $PAM^{MB441B}$>$shi^{ts}$ 27°C | 6 10 | | 0.0312 |
| 8 (a) Arousability during wake, high amplitude | $PAM^{MB441B}$-Gal4 21°C vs $PAM^{MB441B}$>TrpA1 21°C | 10 10 | Two-way ANOVA F (4,78) = 1.806 Tukey's multiple comparisons test | 0.9997 |
| | UAS-TrpA1 21°C vs $PAM^{MB441B}$>TrpA1 21°C | 8 10 | | >0.9999 |
| | $PAM^{MB441B}$-Gal4 21°C vs $PAM^{MB441B}$>$shi^{ts}$ 21°C | 10 10 | | 0.8307 |
| | UAS- $shi^{ts}$ 21°C vs $PAM^{MB441B}$>$shi^{ts}$ 21°C | 6 10 | | 0.7400 |
| | $PAM^{MB441B}$-Gal4 27°C vs $PAM^{MB441}$B>TrpA1 27°C | 10 10 | | 0.0052 |
| | UAS-TrpA1 27°C vs $PAM^{MB441B}$>TrpA1 27°C | 8 10 | | 0.4960 |
| | $PAM^{MB441B}$-Gal4 27°C vs $PAM^{MB441}$B>$shi^{ts}$ 27°C | 10 10 | | 0.9844 |
| | UAS- $shi^{ts}$ 27°C vs $PAM^{MB441B}$>$shi^{ts}$ 27°C | 6 10 | | >0.9999 |
| 8 (a) Sleep duration | $PAM^{MB441B}$-Gal4 21°C vs $PAM^{MB441B}$>TrpA1 21°C | 26 28 | Two-way ANOVA F (4,213) = 3.455 Tukey's multiple comparisons test | >0.9999 |
| | UAS-TrpA1 21°C vs $PAM^{MB441B}$>TrpA1 21°C | 32 28 | | >0.9999 |
| | $PAM^{MB441B}$-Gal4 21°C vs $PAM^{MB441B}$>$shi^{ts}$ 21°C | 26 22 | | 0.8034 |
| | UAS- $shi^{ts}$ 21°C vs $PAM^{MB441B}$>$shi^{ts}$ 21°C | 39 22 | | <0.0001 |
| | $PAM^{MB441B}$-Gal4 27°C vs $PAM^{MB441}$B>TrpA1 27°C | 26 28 | | 0.0270 |
| | UAS-TrpA1 27°C vs $PAM^{MB441B}$>TrpA1 27°C | 32 28 | | >0.9999 |
| | $PAM^{MB441B}$-Gal4 27°C vs $PAM^{MB441}$B>$shi^{ts}$ 27°C | 26 22 | | 0.0261 |
| | UAS- $shi^{ts}$ 27°C vs $PAM^{MB441B}$>$shi^{ts}$ 27°C | 39 22 | | 0.8203 |
| 8 (a) Basal locomotion | $PAM^{MB441B}$-Gal4 21°C vs $PAM^{MB441B}$>TrpA1 21°C | 26 28 | Two-way ANOVA F (4,213) = 0.811 Tukey's multiple comparisons test | >0.9999 |
| | UAS-TrpA1 21°C vs $PAM^{MB441B}$>TrpA1 21°C | 32 28 | | 0.3733 |
| | $PAM^{MB441B}$-Gal4 21°C vs $PAM^{MB441B}$>$shi^{ts}$ 21°C | 26 22 | | 0.9451 |
| | UAS- $shi^{ts}$ 21°C vs $PAM^{MB441B}$>$shi^{ts}$ 21°C | 39 22 | | 0.0005 |
| | $PAM^{MB441B}$-Gal4 27°C vs $PAM^{MB441}$B>TrpA1 27°C | 26 28 | | 0.9904 |
| | UAS-TrpA1 27°C vs $PAM^{MB441B}$>TrpA1 27°C | 32 28 | | 0.0048 |
| | $PAM^{MB441B}$-Gal4 27°C vs | 26 | | >0.9999 |

| | | | | |
|---|---|---|---|---|
| | $PAM^{MB441}B{>}shi^{ts}$ 27°C | 22 | | |
| | UAS- $shi^{ts}$ 27°C vs<br>$PAM^{MB441B}{>}shi^{ts}$ 27°C | 39<br>22 | | 0.0022 |
| 8 (a) Number of sleep bouts | $PAM^{MB441B}$-Gal4 21°C vs<br>$PAM^{MB441B}{>}$TrpA1 21°C | 26<br>28 | Two-way ANOVA<br>F (4,213) = 0.831<br>Tukey's multiple comparisons test | 0.9943 |
| | UAS-TrpA1 21°C vs<br>$PAM^{MB441B}{>}$TrpA1 21°C | 32<br>28 | | 0.9998 |
| | $PAM^{MB441B}$-Gal4 21°C vs<br>$PAM^{MB441B}{>}shi^{ts}$ 21°C | 26<br>22 | | 0.9854 |
| | UAS- $shi^{ts}$ 21°C vs<br>$PAM^{MB441B}{>}shi^{ts}$ 21°C | 39<br>22 | | >0.9999 |
| | $PAM^{MB441B}$-Gal4 27°C vs<br>$PAM^{MB441}B{>}$TrpA1 27°C | 26<br>28 | | >0.9999 |
| | UAS-TrpA1 27°C vs<br>$PAM^{MB441B}{>}$TrpA1 27°C | 32<br>28 | | 0.9953 |
| | $PAM^{MB441B}$-Gal4 27°C vs<br>$PAM^{MB441}B{>}shi^{ts}$ 27°C | 26<br>22 | | 0.9770 |
| | UAS- $shi^{ts}$ 27°C vs<br>$PAM^{MB441B}{>}shi^{ts}$ 27°C | 39<br>22 | | 0.0793 |
| 8 (a) Length of sleep bouts | $PAM^{MB441B}$-Gal4 21°C vs<br>$PAM^{MB441B}{>}$TrpA1 21°C | 26<br>28 | Two-way ANOVA<br>F (4,213) = 1.956<br>Tukey's multiple comparisons test | 0.9999 |
| | UAS-TrpA1 21°C vs<br>$PAM^{MB441B}{>}$TrpA1 21°C | 32<br>28 | | 0.9949 |
| | $PAM^{MB441B}$-Gal4 21°C vs<br>$PAM^{MB441B}{>}shi^{ts}$ 21°C | 26<br>22 | | >0.9999 |
| | UAS- $shi^{ts}$ 21°C vs<br>$PAM^{MB441B}{>}shi^{ts}$ 21°C | 39<br>22 | | 0.9965 |
| | $PAM^{MB441B}$-Gal4 27°C vs<br>$PAM^{MB441}B{>}$TrpA1 27°C | 26<br>28 | | 0.7757 |
| | UAS-TrpA1 27°C vs<br>$PAM^{MB441B}{>}$TrpA1 27°C | 32<br>28 | | >0.9999 |
| | $PAM^{MB441B}$-Gal4 27°C vs<br>$PAM^{MB441}B{>}shi^{ts}$ 27°C | 26<br>22 | | >0.9999 |
| | UAS- $shi^{ts}$ 27°C vs<br>$PAM^{MB441B}{>}shi^{ts}$ 27°C | 39<br>22 | | 0.5662 |
| 8 (b) Arousability during wake | $PAM^{MB441B}$ >mCherry-RNAi;dcr2 vs<br>$PAM^{MB441B}$ >TH-RNAi;dcr2 | 8<br>10 | One-way ANOVA<br>F (2,24) = 4.392<br>Tukey's multiple comparisons test | 0.3388 |
| | UAS-TH-RNAi;UAS-dcr2 vs<br>$PAM^{MB441B}$ >TH-RNAi;dcr | 9<br>10 | | 0.0179 |
| 8 (b) Sleep duration | $PAM^{MB441B}$ >mCherry-RNAi;dcr2 vs<br>$PAM^{MB441B}$ >TH-RNAi;dcr2 | 59<br>56 | Kruskal-Wallis (2,166) = 27.62<br>Dunn's multiple | 0.3895 |
| | UAS-TH-RNAi;UAS-dcr2 vs<br>$PAM^{MB441B}$ >TH-RNAi;dcr | 54<br>56 | | 0.0010 |

| | | | | |
|---|---|---|---|---|
| | | | comparisons test | |
| 8 (b) Basal locomotion | PAM$^{MB441B}$ >mCherry-RNAi;dcr2 vs PAM$^{MB441B}$ >TH-RNAi;dcr2 | 59<br>56 | One-way ANOVA F (2,166) = 14.5 Tukey's multiple comparisons test | 0.7361 |
| | UAS-TH-RNAi;UAS-dcr2 vs PAM$^{MB441B}$ >TH-RNAi;dcr | 54<br>56 | | 0.0001 |
| 8 (b) Number of sleep bouts | PAM$^{MB441B}$ >mCherry-RNAi;dcr2 vs PAM$^{MB441B}$ >TH-RNAi;dcr2 | 59<br>56 | Kruskal-Wallis (2,166) = 23.55 Dunn's multiple comparisons test | 0.2523 |
| | UAS-TH-RNAi;UAS-dcr2 vs PAM$^{MB441B}$ >TH-RNAi;dcr | 54<br>56 | | 0.0067 |
| 8 (b) Length of sleep bouts | PAM$^{MB441B}$ >mCherry-RNAi;dcr2 vs PAM$^{MB441B}$ >TH-RNAi;dcr2 | 59<br>56 | Kruskal-Wallis (2,166) = 2.206 Tukey's multiple comparisons test | >0.9999 |
| | UAS-TH-RNAi;UAS-dcr2 vs PAM$^{MB441B}$ >TH-RNAi;dcr | 54<br>56 | | 0.4771 |
| **Extended Data Figure 9** | | | | |
| 9 (a) | Wild type 30 °C vs Wild type 40 °C | 4<br>4 | T-test t=4.846 df=6 | 0.0029 |
| 9 (b) peptone sleeping flies | Regular vs Regular + peptone | 10<br>9 | T-test t=0.077 df=17 | 0.9395 |
| 9 (b) peptone awake flies | Regular vs Regular + peptone | 10<br>9 | T-test t=0.3982 df=17 | 0.6954 |
| 9 (b) gut-specific CCHa1 depletion sleeping flies | EECG>GFP-RNAi vs EECG>CCHa1-RNAi | 9<br>9 | One-way ANOVA F(2,24)=0.7927 Tukey's multiple comparisons test | 0.9856 |
| | UAS-CCHa1-RNAi vs EECG>CCHa1-RNAi | 9<br>9 | | 0.5838 |
| 9 (b) gut-specific CCHa1 depletion awake flies | EECG>GFP-RNAi vs EECG>CCHa1-RNAi | 9<br>9 | One-way ANOVA F(2,24)=0.0477 Tukey's multiple comparisons test | 0.9581 |
| | UAS-CCHa1-RNAi vs EECG>CCHa1-RNAi | 9<br>9 | | 0.9995 |

| | | | | |
|---|---|---|---|---|
| 9 (b) MB441B-specific CCHa1R depletion sleeping flies | $PAM^{MB441B}$>GFP-RNAi vs $PAM^{MB441B}$>CCHa1R-RNAi | 10 11 | One-way ANOVA F(2,29)=5.215 Tukey's multiple comparisons test | 0.0201 |
| | UAS-CCHA1R-RNAi vs $PAM^{MB441B}$>CCHa1-RNAi | 11 11 | | 0.9933 |
| 9 (b) MB441B-specific CCHa1R depletion awake flies | $PAM^{MB441B}$>GFP-RNAi vs $PAM^{MB441B}$>CCHa1R-RNAi | 10 11 | One-way ANOVA F(2,29)=2.352 Tukey's multiple comparisons test | 0.0988 |
| | UAS-CCHA1R-RNAi vs $PAM^{MB441B}$>CCHa1-RNAi | 11 11 | | 0.7436 |
| 9 (c) | $PAM^{MB441B}$>CaLexA no temperature vs $PAM^{MB441B}$>CaLexA 28h after temperature | 55 47 | Mann-Whitney | 0.8726 |
| 9 (d) Arousability during wake, low amplitude | $MBON^{MB083C}$-Gal4 21°C vs $MBON^{MB083C}$ >TrpA1 21°C | 10 10 | Two-way ANOVA F (4,78) = 3.028 Tukey's multiple comparisons test | >0.9999 |
| | UAS-TrpA1 21°C vs $MBON^{MB083C}$ >TrpA1 21°C | 8 10 | | 0.8533 |
| | $MBON^{MB083C}$ -Gal4 21°C vs $MBON^{MB083C}$ >$shi^{ts}$ 21°C | 10 10 | | >0.9999 |
| | UAS- $shi^{ts}$ 21°C vs $MBON^{MB083C}$ >$shi^{ts}$ 21°C | 6 10 | | 0.8152 |
| | $MBON^{MB083C}$ -Gal4 27°C vs $MBON^{MB083C}$ >TrpA1 27°C | 10 10 | | 0.0007 |
| | UAS-TrpA1 27°C vs $MBON^{MB083C}$ >TrpA1 27°C | 8 10 | | 0.3325 |
| | $MBON^{MB083C}$ -Gal4 27°C vs $MBON^{MB083C}$ >$shi^{ts}$ 27°C | 10 10 | | 0.0002 |
| | UAS- $shi^{ts}$ 27°C vs $MBON^{MB083C}$ >$shi^{ts}$ 27°C | 6 10 | | 0.0642 |
| 9 (d) Arousability during wake, high amplitude | $MBON^{MB083C}$-Gal4 21°C vs $MBON^{MB083C}$ >TrpA1 21°C | 10 10 | Two-way ANOVA F (4,78) = 0.8467 Tukey's multiple comparisons test | 0.6893 |
| | UAS-TrpA1 21°C vs $MBON^{MB083C}$ >TrpA1 21°C | 8 10 | | >0.9999 |
| | $MBON^{MB083C}$ -Gal4 21°C vs $MBON^{MB083C}$ >$shi^{ts}$ 21°C | 10 10 | | 0.9994 |
| | UAS- $shi^{ts}$ 21°C vs $MBON^{MB083C}$ >$shi^{ts}$ 21°C | 6 10 | | >0.9999 |
| | $MBON^{MB083C}$ -Gal4 27°C vs $MBON^{MB083C}$ >TrpA1 27°C | 10 10 | | 0.9728 |
| | UAS-TrpA1 27°C vs | 8 | | 0.9990 |

| | | | | |
|---|---|---|---|---|
| | MBON$^{MB083C}$ >TrpA1 27°C | 10 | | |
| | MBON$^{MB083C}$ -Gal4 27°C vs<br>MBON$^{MB083C}$ >shi$^{ts}$ 27°C | 10<br>10 | | 0.1988 |
| | UAS- shi$^{ts}$ 27°C vs<br>MBON$^{MB083C}$ >shi$^{ts}$ 27°C | 6<br>10 | | 0.9999 |
| 9 (d) Sleep duration | MBON$^{MB083C}$-Gal4 21°C vs<br>MBON$^{MB083C}$ >TrpA1 21°C | 33<br>53 | Two-way ANOVA<br>F (4,320) = 22.75<br>Tukey's multiple comparisons test | <0.0001 |
| | UAS-TrpA1 21°C vs<br>MBON$^{MB083C}$ >TrpA1 21°C | 35<br>53 | | <0.0001 |
| | MBON$^{MB083C}$ -Gal4 21°C vs<br>MBON$^{MB083C}$ >shi$^{ts}$ 21°C | 33<br>40 | | 0.8776 |
| | UAS- shi$^{ts}$ 21°C vs<br>MBON$^{MB083C}$ >shi$^{ts}$ 21°C | 43<br>40 | | <0.0001 |
| | MBON$^{MB083C}$ -Gal4 27°C vs<br>MBON$^{MB083C}$ >TrpA1 27°C | 33<br>53 | | <0.0001 |
| | UAS-TrpA1 27°C vs<br>MBON$^{MB083C}$ >TrpA1 27°C | 35<br>53 | | <0.0001 |
| | MBON$^{MB083C}$ -Gal4 27°C vs<br>MBON$^{MB083C}$ >shi$^{ts}$ 27°C | 33<br>40 | | 0.0218 |
| | UAS- shi$^{ts}$ 27°C vs<br>MBON$^{MB083C}$ >shi$^{ts}$ 27°C | 43<br>40 | | 0.0576 |
| 9 (d) Basal locomotion | MBON$^{MB083C}$-Gal4 21°C vs<br>MBON$^{MB083C}$ >TrpA1 21°C | 33<br>53 | Two-way ANOVA<br>F (4,320) = 3.759<br>Tukey's multiple comparisons test | 0.9223 |
| | UAS-TrpA1 21°C vs<br>MBON$^{MB083C}$ >TrpA1 21°C | 35<br>53 | | 0.0003 |
| | MBON$^{MB083C}$ -Gal4 21°C vs<br>MBON$^{MB083C}$ >shi$^{ts}$ 21°C | 33<br>40 | | >0.9999 |
| | UAS- shi$^{ts}$ 21°C vs<br>MBON$^{MB083C}$ >shi$^{ts}$ 21°C | 43<br>40 | | <0.0001 |
| | MBON$^{MB083C}$ -Gal4 27°C vs<br>MBON$^{MB083C}$ >TrpA1 27°C | 33<br>53 | | 0.0011 |
| | UAS-TrpA1 27°C vs<br>MBON$^{MB083C}$ >TrpA1 27°C | 35<br>53 | | 0.4863 |
| | MBON$^{MB083C}$ -Gal4 27°C vs<br>MBON$^{MB083C}$ >shi$^{ts}$ 27°C | 33<br>40 | | 0.9997 |
| | UAS- shi$^{ts}$ 27°C vs<br>MBON$^{MB083C}$ >shi$^{ts}$ 27°C | 43<br>40 | | <0.0001 |
| 9 (d) Number of sleep bouts | MBON$^{MB083C}$-Gal4 21°C vs<br>MBON$^{MB083C}$ >TrpA1 21°C | 33<br>53 | Two-way ANOVA<br>F (4,320) = 7.3<br>Tukey's multiple comparisons test | 0.3420 |
| | UAS-TrpA1 21°C vs<br>MBON$^{MB083C}$ >TrpA1 21°C | 35<br>53 | | 0.9499 |
| | MBON$^{MB083C}$ -Gal4 21°C vs<br>MBON$^{MB083C}$ >shi$^{ts}$ 21°C | 33<br>40 | | >0.9999 |
| | UAS- shi$^{ts}$ 21°C vs<br>MBON$^{MB083C}$ >shi$^{ts}$ 21°C | 43<br>40 | | 0.9997 |
| | MBON$^{MB083C}$ -Gal4 27°C vs<br>MBON$^{MB083C}$ >TrpA1 27°C | 33<br>53 | | >0.9999 |

| | | | | |
|---|---|---|---|---|
| | UAS-TrpA1 27°C vs MBON$^{MB083C}$ >TrpA1 27°C | 35 53 | | 0.0295 |
| | MBON$^{MB083C}$ -Gal4 27°C vs MBON$^{MB083C}$ >shi$^{ts}$ 27°C | 33 40 | | 0.9908 |
| | UAS- shi$^{ts}$ 27°C vs MBON$^{MB083C}$ >shi$^{ts}$ 27°C | 43 40 | | 0.0005 |
| 9 (d) Length of sleep bouts | MBON$^{MB083C}$-Gal4 21°C vs MBON$^{MB083C}$ >TrpA1 21°C | 33 53 | Two-way ANOVA $F (4,320) = 12.21$ Tukey's multiple comparisons test | 0.9993 |
| | UAS-TrpA1 21°C vs MBON$^{MB083C}$ >TrpA1 21°C | 35 53 | | <0.0001 |
| | MBON$^{MB083C}$ -Gal4 21°C vs MBON$^{MB083C}$ >shi$^{ts}$ 21°C | 33 40 | | >0.9999 |
| | UAS- shi$^{ts}$ 21°C vs MBON$^{MB083C}$ >shi$^{ts}$ 21°C | 43 40 | | 0.0463 |
| | MBON$^{MB083C}$ -Gal4 27°C vs MBON$^{MB083C}$ >TrpA1 27°C | 33 53 | | 0.0022 |
| | UAS-TrpA1 27°C vs MBON$^{MB083C}$ >TrpA1 27°C | 35 53 | | 0.0324 |
| | MBON$^{MB083C}$ -Gal4 27°C vs MBON$^{MB083C}$ >shi$^{ts}$ 27°C | 33 40 | | 0.9564 |
| | UAS- shi$^{ts}$ 27°C vs MBON$^{MB083C}$ >shi$^{ts}$ 27°C | 43 40 | | 0.1902 |
